# Distinct Quaternary States, Intermediates, and Autoinhibition During Loading of the DnaB-Replicative Helicase by the Phage λP Helicase Loader

**DOI:** 10.1101/2022.12.30.522210

**Authors:** Abhipsa Shatarupa, Dhanjai Brown, Paul Dominic B. Olinares, Jillian Chase, Eta Isiorho, Brian T. Chait, David Jeruzalmi

## Abstract

Replicative helicases require loader proteins for assembly at the origins of DNA replication. Multiple copies of the bacteriophage λP (P) loader bind to and load the *E. coli* DnaB (B) replicative helicase on replication-origin-derived single-stranded DNA. We find that the *E. coli* DnaB•λP complex exists in two forms: B_6_P_5_ and B_6_P_6_. In the 2.66 Å cryo-EM model of B_6_P_5_, five copies of the λP loader assemble into a crown-like shape that tightly grips DnaB. In this complex, closed planar DnaB is reconfigured into an open spiral with a sufficiently sized breach to permit ssDNA to enter an internal chamber. The transition to the open spiral involves λP-mediated changes to the Docking Helix (DH)–Linker Helix (LH) interface. The loader directly stabilizes the open spiral. Unexpectedly, one λP chain in B_6_P_5_ is bound across the breach, precluding entry of replication-origin-derived ssDNA into DnaB’s central chamber. We suggest that the B_6_P_6_ complex is an early intermediate in the helicase activation pathway wherein neither the DnaB helicase nor the λP loader has attained its final form. DnaB in this complex adopts a partially open planar configuration, termed ajar planar. The partially ordered λP loader assembly features a much looser interaction with DnaB. The ssDNA and ATP sites in both complexes are in a configuration ill-suited for binding or hydrolysis. Our work specifies the conformational changes required for the intermediate B_6_P_6_ to transition to B_6_P_5_ on the pathway to recruitment by the initiator protein complex to the replication origin.

**Graphical Abstract:** 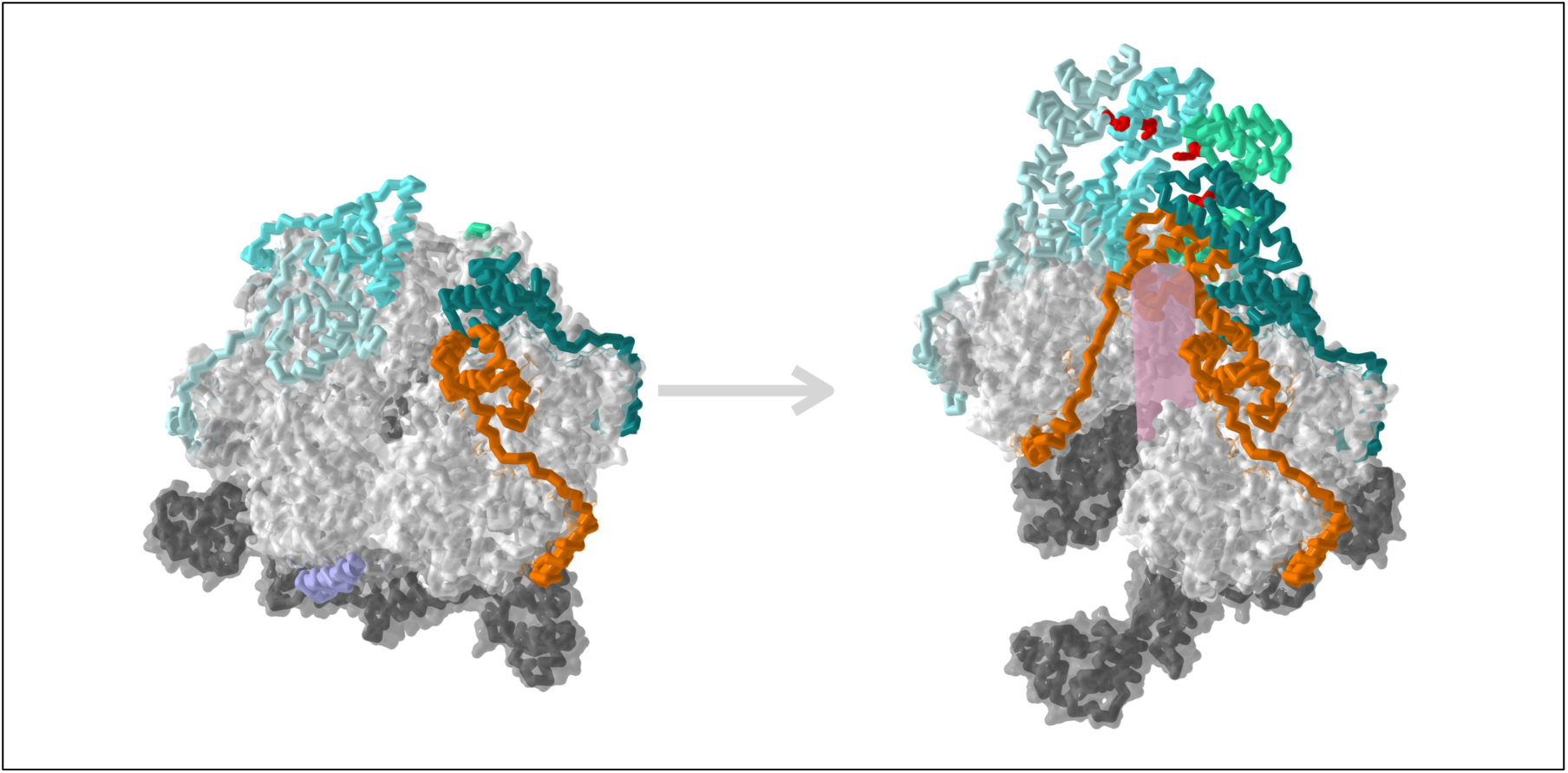

The DnaB helicase loading pathway at the phage λ replication origin populates two intermediate states, distinguished by the number of λP loaders present. The DnaB ring in the B_6_P_6_ complex is planar and partially open. Although it binds six copies of the λP loader, the ajar planar state of DnaB yields inchoate interactions with the loader. During the maturation of the complex, the planar state of DnaB is reconfigured into an open spiral in the B_6_P_5_ complex, which the pentameric λP ensemble now grips tightly. This transition required the eviction of one copy of the loader. Although the breach in the DnaB open spiral is sufficiently sized for entry of ssDNA into the internal chamber, the disposition of one λP chain loader across the single breached interface in DnaB effectively blocks the path to physiological replication origin-derived ssDNA. DnaB is depicted in white/gray ribbon format under a transparent surface. The λP chains are colored orange and shades of blue. The pink cylinder represents the expected path of ssDNA through DnaB.

## Introduction

Efficient separation of duplex DNA into single-strand templates for DNA synthesis is a crucial feature of chromosomal replication systems across all cellular domains of life (1–10). Separation is carried out by the replicative helicase, an oligomeric entity that encircles an internal chamber into which one strand of DNA is loaded while the second strand is excluded. ATP-dependent translocation along the included strand leads to the melting of the duplex. Since bacterial chromosomal DNA is an infinitely long polymer with no free ends, assembling replicative helicases on DNA requires specialized loader proteins.

In bacteria, the replicative helicase (DnaB in most bacteria (henceforth: DnaB), or DnaC in *B. subtilis*) assembles into a two-tiered, ring-shaped homohexameric ensemble around one of the two strands of dsDNA (11–13) (Figure 1 and Supplementary Figure 1 and reviewed in (9, 10, 14–17) and in Supplementary Information). Each DnaB protomer comprises two domains: an amino-terminal domain (NTD) and a carboxy-terminal domain (CTD), linked by a linker helix (LH) element. The NTD and CTD tiers adopt two configurations, termed dilated or constricted, distinguished by the diameter of the central chamber, among other features. Each CTD harbors a helical element, termed the docking helix (DH), which packs against the LH element of an adjacent CTD (11–13). ATP is bound at CTD interfaces, using Walker A and Walker B motifs furnished by one CTD and the arginine finger β-hairpin motif by an adjacent CTD to mediate hydrolysis. Residues from each CTD also contact the phosphate backbone of ssDNA within the inner chamber (18).

**Figure 1.**
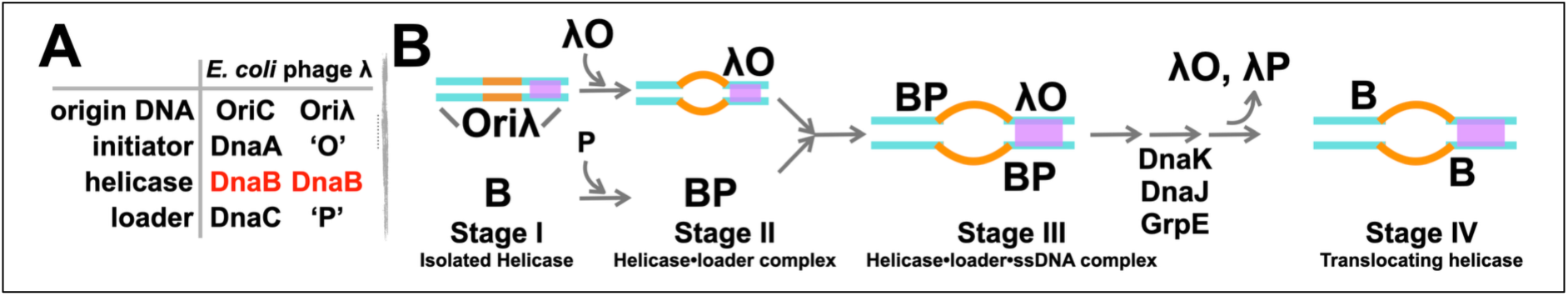
Loading of the bacterial replicative helicase during the initiation of DNA replication. A) Molecules associated with initiating DNA replication in *E. coli* and phage λ. Both systems utilize *E. coli* DnaB (red). B) The loading of the bacterial DnaB helicase onto the origin DNA occurs at the nexus of two pathways: their products merge into the complete initiation complex at the replication origin (17, 111, 112). The first sub-complex is the oligomeric initiator protein complex (*E. coli:* DnaA or phage λ: O) formed at the corresponding replication origin (the four λO sites are colored in purple); a key outcome of this complex is the initial melting of the origin’s DNA unwinding element (DUE, orange). Capture and remodeling of the DnaB helicase (designated with the letter ‘B’) by the helicase loader (*E. coli:* DnaC or phage λ: P) produces the second sub-complex (BP). These two pathways converge with the delivery of the loader-bound helicase to the origin. The resulting initiator-helicase-loader complex at the origin is stabilized by interactions between the initiator, helicase, and loader (1, 8, 113, 114). A series of additional binding and remodeling events, including the engagement of the bacterial chaperones DnaK, DnaJ, and GrpE, culminate in assembling a pair of replication forks (not shown).

An extensive literature provides insights into the structure and function of helicase loaders in bacteria, including *E. coli* DnaC, phage λP (Figure 1), *B. subtilis* DnaI, phage T4 gene 59, phage SPP1 G39P, phage P2 B, DciA, and DopE (1, 7, 19–30). Several schemes for helicase assembly on nucleic acid substrates have been described, referred to here as (1) ring-opening, (2) ring-forming, or (3) ring-closing ((10, 31, 32) and Supplementary Information). Two well-studied bacterial loaders, *E. coli* DnaC and phage λP, are ring-opening loaders; although unrelated in sequence or structure, they implement similar loading mechanisms. Both loaders share an overall architecture, featuring a globular domain attached to an element termed the lasso or grappling hook (15, 33, 34). Both capture and remodel closed planar DnaB into an open right-handed spiral, maintaining both tiers in the constricted conformation. Both suppress DnaB’s ATPase and DNA unwinding activities (35–38). Remodeling creates breaches in DnaB’s two tiers of sufficient size to allow access to the inner chamber by physiological ssDNA substrates produced by the remodeling of the replication origins by the replication initiator protein (*E. coli:* DnaA or phage λ: O) (19, 39–41). The structural resemblance between the two loaders is limited to an alpha helix in the lasso/grappling hook element, positioned between the DH and LH helices of two adjacent DnaB subunits. Remarkably, the protein chain direction of these elements diverges (λP: N-C; DnaC: C-N); this unusual feature reflects convergent evolution (15).

Maturation of the replication origin complex into the replisome requires the eviction of the DnaB-inhibiting loader. DnaC (34, 35, 42) and λP (43, 44) utilize distinct biochemical mechanisms to evict the loader. In phage λ, the tight *E. coli* DnaB-λP loader (BP) complex (37) necessitates the involvement of the host heat shock factors DnaJ, DnaK, and GrpE (40, 45–48) to evict the loader and activate the helicase.

The 4.1 Å single-particle cryogenic-electron microscopy (cryo-EM) structure of the BP complex informed many aspects of the helicase loading reaction (15, 33). However, the limited resolution revealed ∼50% of the structure of the λP loader, as only the carboxy-terminal domains of the five protomers (B_6_P_5_) were visible in the EM map (EMDB: 7076, PDB: 6BBM (33)). Furthermore, the amino acid sequence of the λP loader could not be assigned to its structure, thereby obscuring the helicase and loader interface. As such, the mechanisms of DnaB remodeling, ATPase suppression, and ssDNA entry into the central chamber were incompletely deciphered. We have determined three cryo-EM structures of the BP complex to gain insights into these mechanisms. Two of these structures extend the precision of the B_6_P_5_ complex to 2.66 Å and 2.84 Å; the 2.66 Å structure provides a nearly complete model. In the B_6_P_5_ complex, one copy of λP is entirely resolved, and the other four are nearly so. The resulting pentameric λP ensemble is held together by an extensive interface between its previously invisible domain II sub-structure. In B_6_P_5_, the DnaB NTD and CTD layers are in open spiral configurations, with openings sufficient for ssDNA to access the central chamber. However, unexpectedly, we find that one copy of λP (chain Z) binds across the breach in the DnaB CTD tier to create a significant block to the entry of physiological origin-derived ssDNA into the central chamber of DnaB.

A third structure (3.85 Å) revealed the first view of a BP complex with six copies of the λP loader (B_6_P_6_). The configurations of both DnaB and λP in the B_6_P_6_ entity are strikingly different. Both tiers of DnaB are essentially planar, with the NTD tier topologically closed and the CTD layer ajar relative to B_6_P_5_. The configuration of the six copies of the λP loader and DnaB in B_6_P_6_ implies an inchoate complex that has not reached its final form. We suggest that the ajar planar configuration in B_6_P_6_ is intermediate in the helicase loading pathway between the closed planar and open spiral forms of DnaB.

## Materials and Methods

### Constructs of the λP helicase loader

This work’s biochemical and structural studies relied on a large set of λP constructs (Supplementary Table 1). DNA sequencing (Genewiz/Azenta) was used to verify the integrity of each construct. Extensive efforts identified soluble domains corresponding to the following constructions: amino acids 105:210, 105:205, 105:215, 105:220, 110:210, and 110:215. Each construct included an N-terminal (N-His) or C-terminal (C-His) hexahistidine tag for purification. The construct spanning residues 105:210 provided a sample suitable for X-ray crystallography.

### Protein Biochemistry

Full-length *E. coli* DnaB•λP complex and DnaB were purified using the procedure described in (33), except that 0.2 mM ATP was used instead of 0.5 mM. In addition, full-length *E. coli* DnaB was further chromatographed over a 320 mL Superdex200 size exclusion column equilibrated in 20 mM HEPES-NaOH, pH 7.5, 450 mM NaCl, 5% glycerol (v/v), 2 mM DTT, 0.5 mM MgCl_2_, 0.2 mM ATP. DnaB-containing fractions were concentrated to 23.5 mg/mL, aliquoted, flash-frozen in liquid nitrogen, and stored at −80°C until use.

The *E. coli* DnaB•λP-C-His complex was first purified over nickel-nitrilotriacetic acid beads (Ni-NTA, Qiagen) equilibrated in 50 mM sodium phosphate, pH 7.6, 500 mM NaCl, 10% glycerol (v/v), and 5 mM β-mercaptoethanol (BME). After sequential washing with 25 mM, 50 mM, and 100 mM imidazole in the equilibration buffer, the partially purified DnaB•λP-C-His complex was eluted using successive washes containing 250 mM and 500 mM imidazole in the equilibration buffer. The complex was further purified using methyl hydrophobic interaction chromatography as described in (33). Lastly, the complex was concentrated to ∼9 mg/mL, aliquoted, flash-frozen in liquid nitrogen, and stored at −80°C until use.

A λP construct encompassing residues 105:210 was expressed in *E. coli* BL21 (DE3) cells as described in (33), except the induction proceeded at 16°C for 16 hours. λP105:210 substituted with selenomethionine (SeMet) was prepared for X-ray crystallographic studies as described in (49). Cells expressing N-His-λP105:210 were lysed by sonication (60% amplitude; 3 min; 0.3 secs on, 0.6 secs off, Fisher Scientific Model 500). λP105:210 was recovered in the lysis supernatant, purified over Ni-NTA as described above, and over Superdex200 equilibrated in 100 mM NaCl, 10 mM Tris-HCl pH 7, 5% glycerol (v/v), and 5 mM BME. Fractions containing N-His-λP105:210 were concentrated to 3.5 mg/mL, aliquoted, flash frozen in liquid nitrogen, and stored at −80°C until use.

An N-His-tagged λO construct encompassing residues 156-299 was expressed in *E. coli* BL21 (DE3) cells using ECPM1 medium (50) according to standard techniques. After growth at 37°C to an OD600 of 0.6 – 0.7, IPTG was added to 0.5 mM, and protein expression was allowed to proceed overnight at 25°C. Induced cells were resuspended in 20 mM Tris-HCl, pH 8, 500 mM NaCl, and 5% (V/V) glycerol at a ratio of 5 mL per gram of cells and frozen at −80°C until use.

Cells expressing λO-156-299-N-His were lysed by sonication (70% amplitude; 3 min; 30 secs on, 30 secs off, Fisher Scientific Model 500). The clarified supernatant was chromatographed over Ni-NTA beads equilibrated in 50 mM Tris (pH 7.5), 500 mM KCl, and 5% glycerol (v/v). Following two washes of the Ni-NTA beads with equilibration buffer supplemented with 20 mM and 30 mM imidazole, λO-156-299-N-His was eluted with 250 mM imidazole. Dialysis into 40 mM Tris pH 8, 50 mM NaCl, and 5 mM BME preceded chromatography over Fast Flow Q (Cytiva). The Q column was developed with a gradient from 50 mM to 1 M NaCl in the above buffer. λO-156-299-N-His-containing fractions were precipitated with 70% saturated ammonium sulfate, resuspended, and further chromatographed over a 320 mL Superdex 200 size exclusion column, equilibrated in 20 mM HEPES-KOH pH 7.5, 300 mM NaCl, 5% glycerol, and 5 mM BME. λO-156-299-N-His containing fractions were concentrated to 2 mg/mL, aliquoted, flash-frozen in liquid nitrogen, and stored at −80°C until use.

Protein biochemistry was performed at 4°C.

### X-ray structure determination of λP domain III (residues 105 – 210)

N-His-λP105:210 in 100 mM NaCl, 10 mM Tris-HCl pH 7, 5% glycerol (v/v), and 5 mM BME was thawed on ice and further concentrated to ∼3.5 mg/mL (> 200 μM). High-throughput vapor diffusion (sitting drop) crystallization trials were performed using an Art Robbins Gryphon liquid handling robot in the Macromolecular Crystallization Facility (MCF) of the Structural Biology Initiative at the Advanced Science Research Center, City University of New York. Crystals of λP105:210 were prepared using the sitting drop vapor diffusion method by mixing either 0.1, 0.2, or 0.4 μL of the protein solution and 0.2 μL of a series of commercially available crystallization screens (Qiagen). Conditions that provided initial hits were optimized using bespoke screens prepared at the MCF. Optimized crystals for X-ray diffraction were grown at room temperature in a buffer containing 3 M–4 M NaCl with 0.1 M HEPES-NaOH, pH 7–8, and required 7 days to form. SeMet-derivatized crystals were grown in the same manner. For X-ray diffraction, crystals (>70 μm) were cryoprotected by adding 0.2 μL of LV CryoOil (MiTeGen) directly to the crystallization drop before flash-freezing in liquid nitrogen.

Diffraction data for native and SeMet-derivatized N-His-λP105:210 crystals were recorded at cryogenic temperatures at the 24-ID-E and 24-ID-C (respectively) beamlines at the Northeastern Collaborative Access Team (NECAT) Center for Advanced Macromolecular Crystallography at the Advanced Photon Source at Argonne National Laboratory. Data from native crystals were measured using a wavelength of 0.97918 Å to Bragg spacings of 1.86 Å. Data from SeMet-derivatized crystals were measured at the Se K-edge (12, 662 eV, λ = 0.97918 Å), corresponding to Bragg spacings of 1.86 Å. Both native and derivatized λP105:210 crystallized in space group P3_1_21, with the following cell parameters: a = 49.731, b = 49.731, c = 70.772, alpha = 90°, beta = 90°, and gamma = 120° (Supplementary Table 2). The Matthews coefficient (51) implied the presence of one copy of λP105:210 in the crystallographic asymmetric unit.

Diffraction images were integrated and scaled with HKL2000 (52). Efforts to determine the phases of the λP105:210 structure using molecular replacement and various search models derived from the cryo-EM structure of the DnaB-λP complex did not yield a reliable solution. However, the structure was determined using the single anomalous dispersion (SAD) method on the SeMet-substituted crystals, as implemented in the Autosol feature of the Phenix suite (53). AutoBuild, phenix.refine, and visualization with Coot (54) facilitated the development of the final model, which comprises residues 119 to 192 (residues 105 to 118 and 193 to 210 could not be visualized). The model exhibited a crystallographic R factor, R_work_/R_free,_ of 0.23/0.24.

### Native mass spectrometry (MS)

The *E. coli* DnaB•λP•ssDNA•λO-156-299-N-His complex was assembled by mixing 94.2 µL of 7.96 μM *E. coli* DnaB•λP complex in 20 mM Na-HEPES pH 7.5, 450 mM NaCl, 2 mM DTT, 0.5 mM MgCl_2_, 0.2 mM ATP, 5% glycerol with 15 µL of 100 µM of an aqueous solution of a 43-nucleotide ssDNA (5’-TGACGAATAATCTTTTCTTTTTTCTTTTGTAATAGTGTCTTTT-3’) derived from the DNA unwinding element (DUE) of Oriλ (38) for 20 minutes; the mixture was prepared at a 1:2 molar ratio of BP:ssDNA. 53.1 µL of 212 µM λO in 20 mM HEPES-KOH pH 7.5, 300 mM NaCl, 5% Glycerol, and 5 mM BME was added to this mixture at a 15-fold molar excess relative to BP, and the mixture was incubated for 20 minutes at 4°C. The volume of the reaction mixture was adjusted to 250 µL by adding Milli-Q water and a buffer containing 2.5 M ammonium acetate (pH 7.5) and 2.5 mM magnesium acetate. This yielded a sample containing 3 µM BP, 6 µM ssDNA, and 45 µM λO in 500 mM ammonium acetate, 0.5 mM magnesium acetate, ∼170 mM NaCl, and ∼2% glycerol. The complexes were incubated for an additional 30 minutes at 4°C.

To purify the DnaB•λP•ssDNA•λO-156-299-N-His complex from uncomplexed components, 200 µL of the above mixture was applied to 300 µL of Ni-NTA beads equilibrated in 500 mM ammonium acetate pH 7.5, and 0.5 mM magnesium acetate. After a 45-minute incubation at 4°C, the beads were recovered by centrifugation and washed four times with 300 µL of 500 mM ammonium acetate pH 7.5, 0.5 mM magnesium acetate, and 50 mM imidazole to remove untagged species; each wash included a 30-second incubation at 4°C. Bound complexes were eluted with three washes of 300 µL of 500 mM ammonium acetate (pH 7.5), 0.5 mM magnesium acetate, 250 mM imidazole, and 0.01% Tween-20. SDS-PAGE analysis confirmed the presence of the DnaB, λP, and λO components. Complex-containing fractions were pooled, concentrated over a 100kDa molecular weight cut-off membrane (Spin-X UF Corning Concentrator, 500uL, 431481) at 15, 000 RPM to 0.48 mg/ml, flash-frozen in liquid nitrogen, and stored at −80°C.

A 200 µL volume of the DnaB•λP•ssDNA•λO-156-299-N-His complex was thawed and concentrated using a Microcon centrifugal filter with a 100-kDa molecular weight cut-off (MWCO) (Millipore). To remove residual imidazole, 150 µL of the native mass spectrometry (nMS)-compatible solution (500 mM ammonium acetate, pH 7.5, 0.5 mM magnesium acetate, 0.01% Tween-20, pH 7.5) was then added to the concentrate for another round of centrifugation. To ensure the complete removal of imidazole, the concentrated sample was then buffer-exchanged into an nMS-compatible solution using a Zeba microspin desalting column with a 40-kDa molecular weight cutoff (Thermo Scientific) before nMS characterization.

For nMS analysis, an aliquot (2 – 3 µL) of the buffer-exchanged sample was loaded into a gold-coated quartz capillary tip that was prepared in-house and was electrosprayed into an Exactive Plus EMR instrument (Thermo Fisher Scientific) using a modified static nanospray source (55). The nMS parameters used included: spray voltage, 1.24 kV; capillary temperature, 150 °C; S-lens RF level, 200; resolving power, 8, 750 at m/z of 200; AGC target, 1 × 10^6^; number of microscans, 5; maximum injection time, 200 ms; in-source dissociation (ISD10 V; injection flatapole, 8 V; interflatapole, 4 V; bent flatapole, 4 V; high energy collision dissociation (HCD), 200 V; ultrahigh vacuum pressure, 5.8 × 10^−10^ mbar; total number of scans, 100. Mass calibration in positive EMR mode was performed using cesium iodide. Raw nMS spectra were visualized using Thermo Xcalibur Qual Browser (version 4.2.47). Data processing and spectra deconvolution were performed using UniDec version 4.2.0 (56, 57). The following parameters were used for data processing: Gaussian smoothing, 3; background subtraction, subtract curve 10; smooth charge state distribution, enabled; peak shape function, Gaussian; beta setting (degree of Softmax distribution): 50, Mass range: 50, 000 – 500, 000 Da.

The measured masses are B_6_P_4_: 419, 942 Da, B_6_P_5_: 446, 642 Da, B_6_P_5_ + oriλ ssDNA: 460, 056 Da, and B_6_P_6_: 473, 318 Da. The observed deviations of the measured mass from the expected mass ranged from 0.07% to 0.10% and were due to peak broadening mainly from nonspecific magnesium adduction. In addition, peaks corresponding to DnaB subcomplexes were found, including B (52, 269 Da), B_2_ (104, 551 Da), and B_3_ (156, 899 Da). Although nMS analysis observed several BP assemblies, including a B_6_P_6_ complex and B_6_P_5_-ssDNA species(33), cryo-EM analyses did not reveal the positions of ssDNA or the binding of λO-CTD. PDBO performed all native MS analyses in the laboratory of BTC.

### ATPase rate measurements

An NADH-coupled microplate spectrophotometric assay was employed to measure the rates of ATP hydrolysis (58, 59) by *E. coli* DnaB and to investigate the effect of λP on these rates. DnaB was maintained at a fixed concentration of 100 nM in all experiments. To assess the contribution of unrelated helicases that may have co-purified with the various DnaB entities, we conducted control experiments using a bacterial protein extract prepared according to the exact set of steps described above for DnaB, except that the cells contained the pET24 vector, which did not include either the DnaB or λP genes.

NADH oxidation data were recorded in the presence of 2 mM ATP (GoldBio A-081-100), 2mM phosphoenolpyruvic acid monopotassium salt (PEP), 0.01 U/µL lactate dehydrogenase (Sigma-Aldrich, L1254), 0.002 U/µL pyruvate kinase (Sigma-Aldrich, P7768],) and 0.3 mM NADH in the following buffer: 50 mM HEPES-KOH, pH=7.5, 150 mM potassium acetate, and 8 mM magnesium acetate, 5 mM BME, and 0.25 mg/mL bovine serum albumin (BSA, Fisher #BP9706100). The above components, except for ATP, were combined and placed into wells of a Corning 96-well clear round-bottom UV-transparent Microplate (p/n: 3788). The plate was then warmed to 37 °C for 5 minutes in a pre-warmed SpectraMax M5 plate reader (Molecular Devices). After the warming phase, endpoint readings were taken at a 340 nm wavelength to enable measurement of the path length. This was followed by the addition of 2 mM ATP, mixing through pipetting, and immediately reinserting the plate into the plate reader. Absorbance data were taken at 340 nm every 30 seconds over 30 minutes. Each experiment was performed in triplicate. Raw data was imported into Microsoft Excel for analysis. Each absorbance curve was manually inspected to identify the period during the experiment wherein the slope of the 340 nm absorbance vs. time(s) was constant; in all cases, this period was the complete 30-minute period. The slope of the absorbance curve was obtained by calculating the difference between the final and initial absorbance values and dividing by time in seconds. The resulting slope value was divided by the extinction coefficient of NADH (6, 220 M^-1^ cm^-1^) and the path length (cm) calculated by the plate reader. This calculation produced the rate of ATP hydrolysis in M/s. After correcting for background by subtracting the blank, this value was processed into ATPase activity in turnovers per second by multiplying by the reaction volume (2×10^-4^ L), dividing by the mass of DnaB (g) in the assay, and then multiplying by the molecular weight of monomeric DnaB (52, 390 g/mol).

### Sample Preparation for Single Particle Cryogenic Electron Microscopy

The BP complex in 20 mM HEPES-NaOH pH 7.5, 450 mM NaCl, 2 mM DTT, 0.5 mM MgCl_2_, 0.2 mM ATP, and 5% glycerol (v/v) and λO-156-299-NHis in 20 mM HEPES-KOH pH 7.5, 300 mM NaCl, 5% glycerol (v/v), and 5 mM BME were diluted with glycerol versions of their respective holding buffers to a concentration of ∼0.25%.

Synthetic single-stranded DNA oligonucleotide 5’-TGACGAATAATCTTTTCTTTTTTCTTTTGTAATAGTGTCTTTT-3’ (38) (IDT) was resuspended in Milli-Q water. BP-ssDNA complexes were prepared by mixing protein (1.5 µM) and ssDNA (1.875 µM) at a 1.25 molar excess. BP•λO-156-299-NHis•ssDNA complexes were prepared by mixing BP (1.5 µM) with ssDNA (1.875 µM) and λO-156-299-NHis (2.25 µM) at 1.25 and 1.5 molar excess, respectively.

Protein-DNA complexes were applied to ultra-gold foil grids with holey gold support and 300 mesh size (R 0.6/1, Quantifoil Micro Tools GmbH); the grids were flash-frozen as described (33) and stored in liquid nitrogen until use.

### Acquisition of Cryo-EM Data

This work encompassed three single particle cryo-EM (cryo-EM) data sets: 1) the P45-J50 data set taken from samples that contained the BP complex bound to ssDNA, 2) the P155-J148 data set recorded from the BP•λO-156-299-NHis•ssDNA complex, and 3) the P11-J107 data set from BP complex bound to ssDNA (sample identical to the P45-J50 dataset) (Supplementary Figures 2, 3, 4, 5, 6, 7, Supplementary Information and Supplementary Table 3). The P45-J50 data set produced a 2.66 Å map of the B_6_P_5_ complex. However, no ssDNA was observed in the EM maps. The P155-J148 data set yielded a 3.85 Å model of the B_6_P_6_ complex; however, our maps did not display the λO-156-299-NHis and ssDNA components. The P11-J107 data set produced a 2.84 Å resolution map of the B_6_P_5_ complex (Supplementary Information and Supplementary Methods).

All data were taken on the Titan Krios (FEI, Hillsboro, Oregon) operating at an accelerating voltage of 300 kV and equipped with a Gatan K3 Summit (Gatan, Pleasanton, California) direct electron detector. Data collection was managed by Leginon (60–62). All movies were collected at a pixel size of 1.083 Å. 7, 863 movies were measured for the P45-J50 BP-ssDNA complex with a defocus range of −1 µm to −2.5 µm, 6, 479 for the P155-J148 BP•λO-156-299-NHis•ssDNA complex with a defocus range of −0.6 µm to −2.7 µm, and 25, 595 un-tilted movies and ∼6661 tilted movies for the P11-J107 BP-ssDNA complex with a defocus range of −1 µm to −2.5 µm and 1 µm to −4.5 µm, respectively. The 25°-tilted data set was collected from the same grid to mitigate the expected preferred specimen orientation (63); the tilted data set was measured using similar imaging parameters to those of the un-tilted collection. The dose rate was set to 25.51 e-/Å²/s for P45-J50, 25.59 e-/Å²/s for P155-J148, 25.51 e^-^/Å^2^/s for P11-J107 un-tilted collection, and 25.78 e^-^/Å^2^/s for P11-J107 for 25°-tilted collection, resulting in accumulated doses of 51.01 e-/Å², 51.19 e-/Å², 51.01 e^-^/Å^2^, and 51.55 e^-^/Å^2^, respectively. Forty frames were recorded per movie at 0.05-second intervals over a total duration of 2 seconds. The 2.84 Å B_6_P_5_ model, derived from the P11-J107 data set, is discussed in the Supplementary Information, as it is less complete than the 2.66 Å model from the P45-J50 data set.

Unless otherwise indicated, cryo-EM data sets were processed in CryoSPARC (64–68). Processing was initiated for each dataset by applying patch motion correction and estimating the contrast transfer function (CTF).

### Cryo-EM Image Processing of the P45-J50 Data Set (B_6_P_5_, 2.66 Å)

The 7, 863 movie frames were submitted to particle selection using the ‘Blob Picker’ tool (Supplementary Figure 2). Inspection of the resulting 7, 250, 518 particles in the “inspect picks” job trimmed the data set to 3, 762, 436 particles. This set was extracted with a 300-pixel box size and 2x binning to produce 3, 193, 376 particles. 2D classification with a total of 200 classes yielded 2, 448, 342 particles. 2D templates prepared from this set were used in template picking to produce 10, 556, 526 particles. Another round of inspection in the “inspect picks” job reduced the number to 4, 331, 766 particles. Extraction with a larger box (400 pixels) and 2x binning produced 3, 709, 955 particles. These particles underwent a second round of 2D classification with 200 classes to produce 2, 798, 195 particles. *Ab initio* reconstruction with C1 symmetry enforced produced four classes, A-00, A-01, A-02, and A-03, with ∼780K, ∼473K, ∼996K, and ∼547K particles. Heterogeneous refinement of these classes resulted in four classes termed B-00, B-01, B-02, and B-03 (grey, red, pink, and blue in Supplementary Figure 2), with the following particle counts and map resolutions ∼480K/4.40 Å, ∼1.5 million/4.40 Å, ∼283K/7.83 Å, and ∼528K/4.40 Å. Class B-01, with the largest number of particles, was pushed forward to non-uniform refinement with the ‘optimize per-particle defocus’ parameter enabled to produce a 4.44 Å map. Particles from this map and the B-01 volume then went through a second round of non-uniform refinement with ‘optimize per-group CTF params’ enabled to produce a second 4.44 Å map. The particles from the second round of non-uniform refinement were re-extracted with a larger and unbinned box size of 400 pixels to provide a new collection of ∼1.5 million particles. This revised set of particles and the class B-01 volume above were submitted to a third round of non-uniform refinement. This calculation produced a 2.66 Å map (class D, green in Supplementary Figure 2). Finally, class D underwent sharpening using DeepEMhancer (69). DeepEMhancer generates maps by applying three algorithms: tight target, wide target, and high-resolution. Inspection of these maps suggested that the wide target was the best option regarding map quality (class E, magenta in Supplementary Figure 2). Additional details about the final volume appear in Supplementary Figure 3. Further processing of Class B-00 produced a 2.75 Å map (class C). However, this map proved to be of lower quality than the 2.66 Å map described above; as such, it was not further pursued.

### Cryo-EM Image Processing of the P155-J148 Data Set (B_6_P_6_, 3.85 Å)

The 6, 479 movie frames were submitted to particle extraction using Topaz (70) and the pre-trained ResNet8 (64 units) model (Supplementary Figure 4). The extracted particles were processed using the “inspect picks” job, resulting in 6, 317, 003 particles. They were then extracted with a 320-pixel box size and 2x binning, yielding 5, 694, 258 particles. 2D classification with 200 classes produced 3, 165, 479 particles. *Ab initio* reconstruction with these particles, with enforcement of C1 symmetry, resulted in five classes named I-00 to I-04, with 580K, 600K, 560K, 840K, and 585K particles in the various classes. Class I-03, with the highest particle count, was submitted to Topaz training using the ResNet8 model architecture. The trained Topaz model was then used to re-extract 2, 315, 940 particles using a 200-pixel box size (binned/Fourier cropped).

2D classification, and *ab-initio* reconstruction with C1 symmetry produced 4 classes: class II-00, II-01, II-02, and II-03 with ∼422K, ∼312K, ∼315K, and ∼472K particles, respectively. Heterogeneous refinement of the above 4 classes produced: class III-00, III-01, III-02, and III-03, with ∼325K (4.40 Å), ∼338K (4.40 Å), ∼264K (4.40 Å), and ∼590K (4.40 Å) particles/resolutions, respectively. Class III-02 was not further processed because of its comparatively low particle count and a distorted volume that appeared stretched in all directions and which could not be reconciled with known structures of DnaB or the BP complex. Class III-03 volume appeared as a distinct two-tiered DnaB structure, whereas maps from classes III-00 and III-01 resembled DnaB but lacked structural information. Consequently, only the III-03 volume was selected for further refinement.

To refine the map further, 1, 256, 507 particles from all three classes (III-00, 01, and 03) were selected for the next round of heterogeneous refinement. Since heterogeneous refinement requires multiple input maps of the same or different types, we included the class III-03 volume three times. We chose the Class III-03 map because it was the most well-defined among all the Class III volumes and had the highest number of particles. Heterogeneous refinement yielded three classes, characterized by the following numbers of particles and resolutions: IV-00 (∼354, 000 particles, 4.40 Å), IV-01 (∼581, 000 particles, 4.40 Å), and IV-02 (∼322, 000 particles, 4.40 Å). These volumes are grey, pink, and green in Supplementary Figure 4. Since multiple particle sets can be used, but only a single volume can be provided for non-uniform refinement, particles from classes IV-00, IV-01, and IV-02 were combined, incorporating the class IV-01 volume with the highest particle count, and submitted to non-uniform refinement. The resulting volume (4.44 Å) was designated class V (blue in Supplementary Figure 4).

Refined particles from class V were used to extract 1, 250, 415 particles at a full box size of 400 pixels. Non-uniform refinement produced a 3.00 Å map (Class VI, red in Supplementary Figure 4). The 3.00 Å resolution, based on the Gold-Standard Fourier shell correlation (GSFSC) curve, was a positive outcome (not shown). However, the observation that the FSC curve did not drop to zero implied a potential problem, possibly due to the presence of duplicate particles. To address this potential problem, a “remove duplicates” job was applied, which retained ∼573K particles while discarding ∼678K particles. The retained particles were refined against the class V volume (blue), with the ‘optimize per-particle defocus’ parameter enabled to yield a 3.90 Å map. Non-uniform refinement and class V volume, with the ‘optimize per-group CTF params’ and the “fit tilt” and “fit trefoil” parameters, yielded a 3.91 Å map. A last round of non-uniform refinement using the refined particles from the 3.91 Å map and the class V volume, with the ‘optimize per-group CTF params’ and “fit spherical aberration” and “fit tetrafoil” provided a 3.85 Å resolution map (Class VII, purple in Supplementary Figure 4). The class VII volume was sharpened using DeepEMhancer (69). Inspection of the maps suggested that the tight target (class VIII, as shown in Supplementary Figure 4) had the best map quality. Additional details about the final volume appear in Supplementary Figure 5.

### Model Building

#### P45-J50 (B_6_P_5_, 2.66 Å)

To build a model of the B_6_P_5_ complex for refinement, we leveraged the availability of high-resolution crystal structures of the amino-terminal domain of DnaB• (PDB: 1B79, (71)), domain Ill of the λP (1.86 Å, this work), as well as Alphafold models of both *E. coli* DnaB (72) and λP ((73, 74) and Supplementary Information). We reasoned that a model constructed from high-resolution components would provide an excellent starting stereochemistry and perform best in a restrained real-space refinement against our 2.66 Å cryo-EM map in Phenix (75, 76).

Components were placed with MOLREP (77, 78) or guided by an earlier B_6_P_5_ structure (PDB: 6BBM, (33)). Notably, five λP domain II segments, missing from earlier models, could be built into the P45-J50 map. Several segments of DnaB and λP (DnaB residues: 25:30, 166:181, 500:501, and λP residues: 1:23, 78:82, 99:100, 120:122, 186:212, 226:232) could not be placed by superposition; these were built by hand in COOT (79) using the Alphafold models as guides. AlphaFold predicted a straight helix for the N-terminal lasso/domain I. However, this conformation did not fit the density in our EM map, so the helix had to be bent after Glu-14.

The final B_6_P_5_ model for the 2.66 Å P45-J50 map encompasses DnaB residues (chain A: 25:173 and 201:468, chain B: 16:468, chain C: 19:468, chain D: 18:468, chain E: 19:468, and chain F: 24:468) and λP residues (chain V: 40:232, chain W: 40:232, chain X: 40:232, chain Y: 37:232, and chain Z-2:231). Our model includes six ADP molecules bound to six magnesium ions. Although included in the sample submitted to cryo-EM, the P45-J50 map showed no density that could be interpreted as ssDNA.

#### P155-J148 (B_6_P_6_, 3.85 Å)

The P155-J148 map revealed a partially ordered B_6_P_6_ complex. A model of the complex was constructed using the approach described above for the P45-J50 map. The availability of the B_6_P_5_ model also guided model building. While the complete structure of each DnaB chain could be seen, only portions of the six λP loader chains were ordered. In four chains (V, W, Y, and Z), only domains III/IV were visualized; for two chains (U and X), only the C-terminal lasso/domain IV was seen. For all λP chains, the N-terminal lasso/domain I and domain II were disordered. The final model encompassed DnaB residues (chain A: 24:468, chain B: 24:468, chain C: 24:468, chain D: 24:468, chain E: 24:468, and chain F: 24:468) and six protomers of λP (U: 211:232, V: 119:232, W: 119:232, X: 210:233, Y: 119:232, Z: 119:231). Each nucleotide binding site in B_6_P_6_ is filled with ADP/Mg, though the occupancies are not identical. Although the C-terminal domain of λO (residues 156-299) and ssDNA were included in the sample submitted to cryo-EM, the P155-J148 maps showed no density for these entities.

### Model Refinement

Models of the BP complexes described in this work were refined using the RealSpaceRefine tool in Phenix (75, 76) against either maps produced by CryoSPARC (64–68) and/or DeepEMhancer (69). Refinement encompassed cycles with minimization global, NQH flips, and the atomic displacement parameters (adp) options and leveraged Ramachandran and secondary structure restraints. The rigid body option was applied in the first round of refinement. The rigid bodies for all DnaB chains in all the refinements were defined as follows: residues 1:173 (NTD), 174:200 (LH), and 201:468 (CTD). For λP, the rigid bodies were defined as follows: residues 1:40 (domain I), 41:118 (domain II), 119:192 (domain III), and 193:233 (domain IV), and applied only when the domain in question was present in a particular model. The refinement of B_6_P_6_ performed better when the 2.66 Å B_6_P_5_ (P45-J50) model was included as the reference model. We also performed two rounds of model rebuilding with high-resolution components, as described above, followed by re-refinement. This procedure produced high-quality models with excellent statistics, particularly in the less well-defined segments.

### Model Analysis and Visualization

Structural analysis was carried out using the CCP4 software package (80), Coot (79, 81), the Uppsala software suite (82–85), UCSF-CHIMERA (86), Phenix (53, 75, 76, 87–91), PyMOL (92), and as described previously (33). Distances and angles within and between protein structures were calculated using the PyMol Python scripts titled “pairwise_dist” (https://pymolwiki.org/index.php/Pairwise_distances) and “angle_between_domains” (https://pymolwiki.org/index.php/Angle_between_domains). Molecular graphics and figures were generated using UCSF-CHIMERA (86) and PYMOL (92). The software used in this study was sourced from the SBGrid Consortium (93).

## Results

### Two States (B_6_P_5_, B_6_P_6_) of the *E. coli* DnaB Helicase–λP Loader Complex

This work aimed to obtain cryo-EM (94–97) structures of the *E. coli* DnaB– λP Helicase Loader (BP) complex bound to ssDNA and, separately, bound to ssDNA and the carboxy-terminal domain of the λO initiator protein (λO-CTD). We have determined three cryo-EM structures at 2.66 Å, 2.84 Å, and 3.85 Å (Methods, Supplementary Figures 2, 3, 4, 5, 6, 7, Supplementary Information, Supplementary Methods, and Supplementary Table 3). These analyses yielded EM maps of higher resolution than our previous 4.1 Å study of the B_6_P_5_ entity (33) and provided a first look at a second oligomeric form of the BP complex (B_6_P_6_). Native mass spectrometry (nMS) analysis of these samples showed the presence of a B_6_P_5_ ssDNA assembly (33) and a B_6_P_6_ species (Supplementary Figure 8). However, cryo-EM analyses did not reveal the positions of ssDNA in any of the complexes.

Our earlier study provided limited insights into the λP loader, as only the carboxy-terminal domains (domains III and IV) were visible in our maps. We were further limited by a model that lacked protein side chains, since the amino acid sequence could not be assigned to the map. The 2.66 Å cryo-EM map of the B_6_P_5_ complex revealed a nearly complete structure of five copies of the λP loader (Figure 2 and Supplementary Figure 9). One λP chain was entirely seen (chain Z; residues 1:233, Supplementary Figure 10), and four chains were nearly completely visible (Domains II, III, IV, chains V, W, X, and Y; residues ∼40:233). Our maps did not resolve four instances of λP domain I (chain V, W, Y, and X; residues 1:∼39). In the nearly complete model, the B_6_P_5_ complex presents as a four-tier entity with its λP and DnaB components arranged in a right-handed open spiral configuration (Figure 3) whose architecture differs from the translocating form of DnaB (18). A prominent breach is visible between the top and bottom of the open DnaB spiral (Figure 2 and Supplementary Figure 11). The two additional tiers constitute domains I/II and III/IV of λP; the remaining two are the NTD and CTD of DnaB (Figure 2). The NTD and CTD tiers of DnaB in the B_6_P_5_ complex are in the constricted configuration (Supplementary Figure 1 and Supplementary Information). In the B_6_P_5_ complex, the six nucleotide sites on DnaB are filled with ADP, although the site on chain B at the top of the spiral appears to have lower occupancy (Supplementary Figure 12). As noted previously (15, 33), the open spiral geometry of DnaB within the B_6_P_5_ complex leaves the ATP hydrolytic catalytic machinery in positions sub-optimal for catalysis (Supplementary Information and Supplementary Figure 13). The 2.84 Å B_6_P_5_ structure superimposes with the 2.66 Å instance with an RMSD of 0.7 (3043 Cα) and is more fully described in the Supplementary Information.

**Figure 2.**
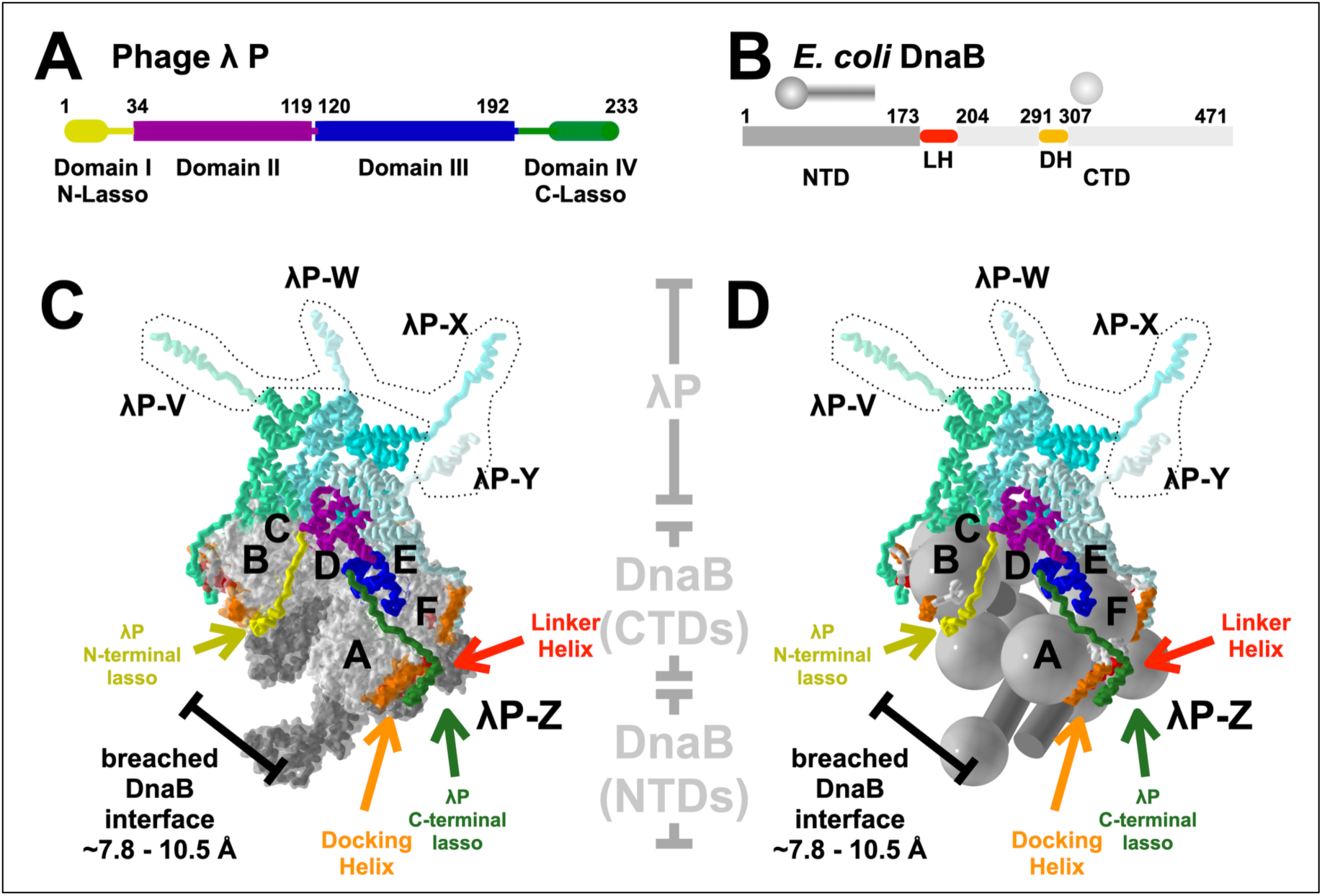
The Structure of the B_6_P_5_ Form of the *E. coli* DnaB – λP Loader. Linear domain architecture of λP (A) and *E. coli* DnaB (B). The amino (NTD) and carboxy-terminal (CTD) layers of DnaB are depicted using a design language that presents the CTDs as large spheres and the NTDs as small spheres/cylinders. C) The DnaB–λP complex, with its five loader molecules (labeled V, W, X, Y, and Z), is depicted in the ribbon representation and colored in shades of blue, save for chain Z, which is colored by domain as illustrated in panel A. The right-handed open spiral of DnaB is depicted as a surface. The DnaB NTD tier is colored in dark gray, and the CTD layer is in light gray, with the DH and LH elements colored and labeled in orange and yellow. The positions of λP domain I for chains V, W, X, and Y (outlined with a dashed line shape), which are not visible in our maps, are modeled for reference. D) Same as panel C, except that DnaB is depicted usingcthe above design language. The CTD spheres of each subunit carry labels corresponding to the underlying chain.

**Figure 3.**
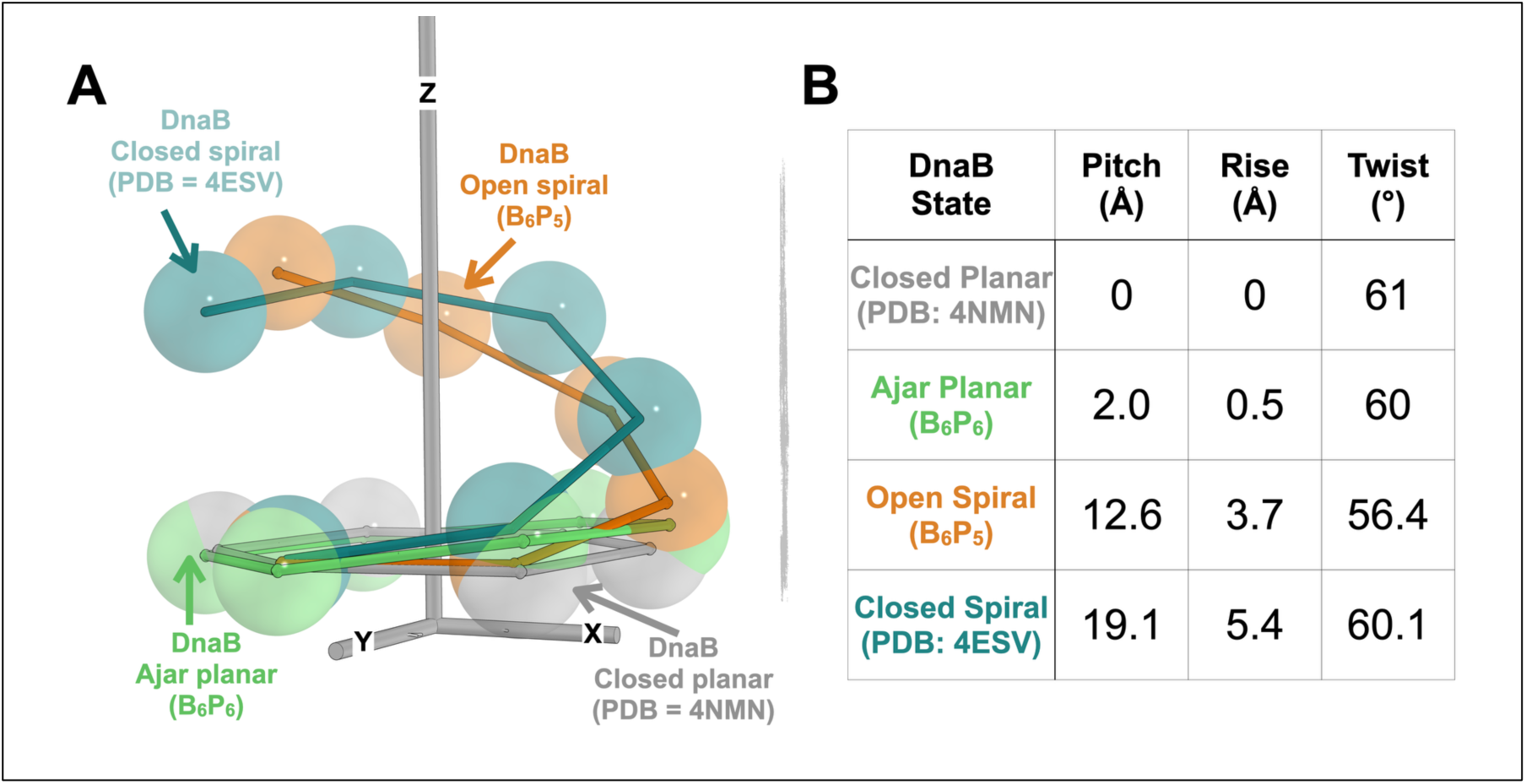
States Populated by the DNA-Binding CTD Tier of DnaB. A) The spirals of the CTD layer of the closed planar form (PDB: 4NMN (12), grey), the ajar planar (B_6_P_6_, this work, green), the open spiral (B_6_P_5_, this work, orange), and the closed spiral (PDB:4ESV (18), deep teal) are displayed. Individual CTDs, depicted as spheres centered around the gravity of the CTD, are interconnected. The underlying coordinates of each DnaB structure were superimposed onto the CTD of the closed planar structure at the bottom of the spiral. The plane of the CTD tier of the closed planar structure is oriented perpendicular to the Z-axis. B) Helical parameters (pitch, rise, and twist) for the specified states of the DnaB CTD.

Previously, we used nMS to demonstrate that preparations of the *E. coli* DnaB–λP helicase loader contained a second oligomeric form with a B_6_P_6_ stoichiometry ((33) and Supplementary Figure 8); our 3.85 Å cryo-EM map revealed the atomic structure of this complex. Models corresponding to domain III/IV were built for four of the six λP chains (V, W, Y, and Z); for the remaining chains (U and X), only the domain IV-C-terminal lasso was visible in our maps. In contrast to the open spiral seen in the B_6_P_5_ complex, DnaB in the B_6_P_6_ complex is nearly planar, with no breach. Below, we provide evidence that DnaB in this complex is partially open, with four of its CTDs further away from their neighbors than in closed, planar DnaB. Unlike B_6_P_5_, the NTD DnaB in the B_6_P_6_ complex is in the dilated configuration, while the CTD tier is constricted. The six nucleotide-binding sites in B_6_P_6_ are filled with ADP, although judging from the density quality, the occupancy of each site differs (Supplementary Figure 12).

### The Pentameric λP Ensemble in B_6_P_5_ Keeps DnaB’s Inner Chamber Sealed

In the nearly complete 2.66 Å B_6_P_5_ complex, five copies of λP assemble in a crown-like shape on top of the CTD layer to effectively seal the breach in the DnaB spiral, thereby preventing entry by physiological replication origin ssDNA (Figure 2). Each monomer of λP features four domains: two compact domains (domain II: residues ∼30 to ∼119; domain III: residues ∼120 to ∼192) flexibly linked (Supplementary Figure 10). The amino (domain I, residues 1 to ∼29) and carboxy termini (domain IV, 193 to 233) of λP are likely intrinsically disordered in solution but fold into an alpha helix linked to domains II and III by an extended segment; we refer to these elements as the N-terminal and C-terminal lasso/grappling hooks. The five previously unseen copies of the λP domain II assemble into an open spiral, displaying an extensive interface that stabilizes the pentameric λP ensemble on the B_6_P_5_ complex. The λP domain II average of ∼1000 Å^2^ of surface area for a total of ∼4000 Å^2^ for the complete ensemble. These values are significant when considered in the context of the NTD (average: ∼1900 A^2^) and CTD (average: ∼2300 A^2^) interfaces of DnaB. The five λP domain IIs circumscribe a volume continuous with DnaB’s inner chamber and exhibit modest flexibility relative to the corresponding domain III position (Supplementary Figure 14). Five domains III/IV of the λP loader ensemble bind at DnaB subunit interfaces; notably, five C-terminal lasso/grappling hooks bind at the nexus of DnaB’s DH and LH elements. As predicted (15, 33), there are no contacts between the B_6_P_5_ complex’s five copies of λP domain III. Superposition suggests that domains III/IV are relatively rigid due to their contact with DnaB (Figure 5, Supplementary Figure 14, and below).

One λP chain (Z) is entirely visible in our maps and, remarkably, adopts a configuration that the other chains cannot access. λP chain Z is positioned to span and effectively block the breach in the open DnaB spiral through its contacts to the A, B, and F subunits. The N-terminal lasso (domain I) of λP binds to chain B at the top of the open DnaB spiral (in the pose in Figure 2). A single breach in DnaB implies that only domain I of one chain can adopt the configuration of the λP chain Z; the remaining domain Is, disordered in our structure, must populate distinct configurations. Notably, Tyr 27 in λP domain I makes a close contact with the ADP in the chain B CTD; the role, if any, of nucleotide in stabilizing the B_6_P_5_ complex requires clarification. Domain II spans the breach between subunits B and A at the top and bottom of the spiral, respectively. Finally, Domain III/IV binds at the interface between the CTDs of subunits A and F. Crucially, the position of the breach-blocking domain II of λP chain Z is maintained through the extensive interface between the five domain IIs of the ensemble (chains V, W, X, Y, and Z). Thus, even though openings in the NTD and CTD tiers of DnaB provide entrances for ssDNA of sufficient size (NTD: ∼7.8 Å; CTD: ∼10.5 Å; Figure 2 and Supplementary Figure 11), the disposition of chain Z, along with the extensive interface between the five domain IIs, effectively seals entry into DnaB’s inner chamber to a physiological ssDNA bubble-shaped substrate at a replication origin (Figure 1).

Comparisons of the higher resolution B_6_P_5_ model with the *E. coli* DnaB•DnaC (B_6_C_6_) complex (15, 33, 34) show that the single-ordered N-terminal lasso/grappling hook (domain I) of λP-chain Z positions the N-terminal helix of this element against the DH helix of DnaB chain B; this position <u>overlaps</u> with the lasso/grappling hook of DnaC, which is bound at the same site (Figure 4 and Supplementary Figure 15). Notably, although the binding modes of λP’s five C-terminal lassos are identical and resemble those of DnaC, save for the change in protein chain direction, the N-terminal lasso binds to the DH helix via a binding mode that differs from that seen with the C-terminal lasso (Supplementary Figure 16). Interestingly, one λP protomer (chain Z) binds at the top and bottom of the breach (in the pose in Figure 2), using the N-terminal and C-terminal lassos, respectively. However, in the DnaC complex, these functions are carried out by two distinct protomers, chain G and chain L, at the top and bottom of the spiral. For both loaders, positioning cognate elements on the DnaB chain B at the top of the spiral could sterically suppress the re-closure of the helicase. This finding represents another example (15) wherein the unrelated λP and DnaC helicase loaders have evolved to converge on the same solution: positioning a loader helix in a similar but distinct binding mode on the DH element at the top of the DnaB spiral.

**Figure 4.**
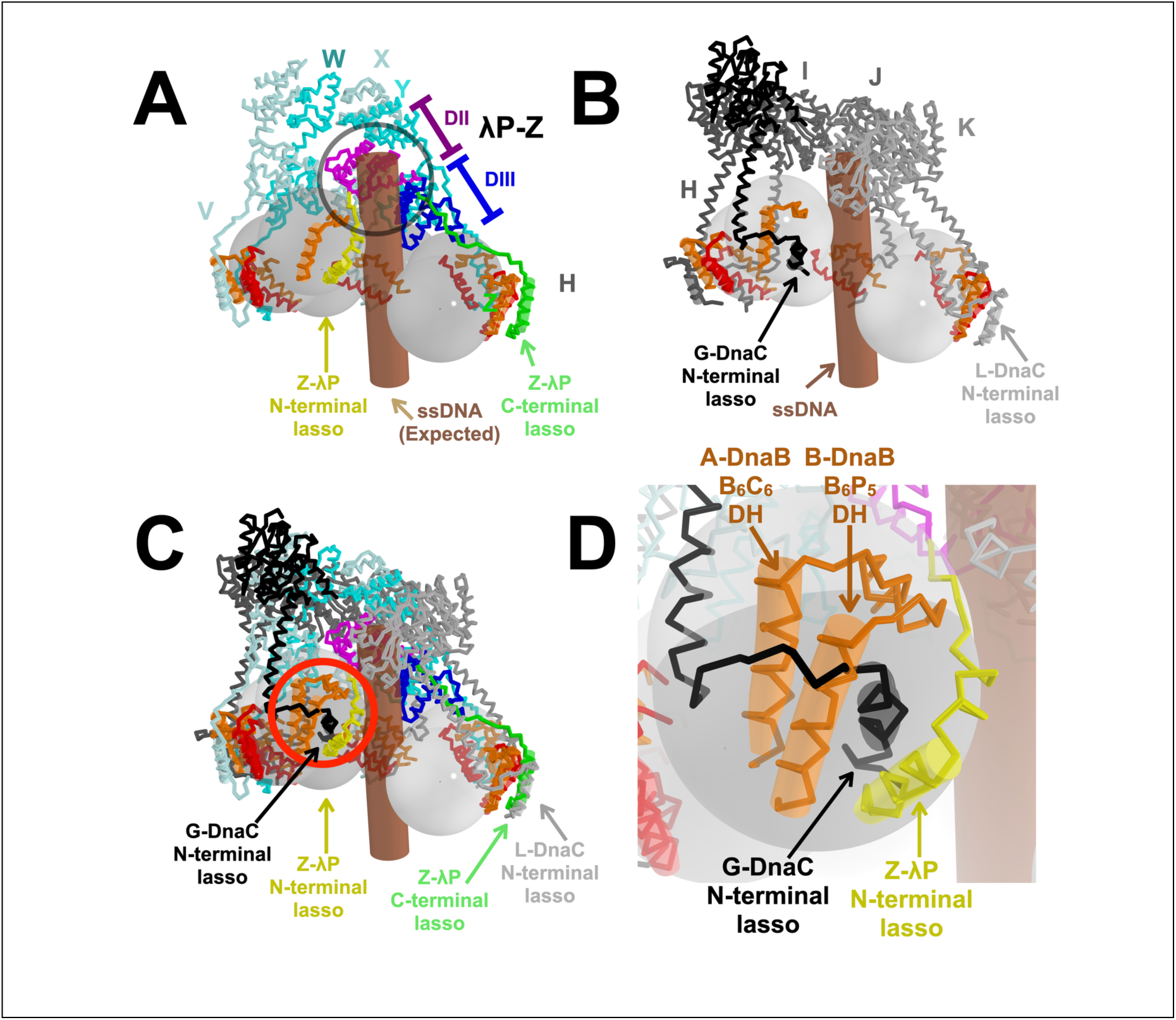
Asymmetry and Convergent Evolution in the λP ensemble. A) In contrast to the other chains, which bind to two DnaB subunits, λP chain Z binds to three subunits (chains B, A, and F), two (chains A and B) located on either side of the breached DnaB interface. λP chain Z is colored by domain (domain I, yellow; domain II, light purple; domain III, blue; domain IV, green), and the remaining λP chains are colored in shades of cyan. The brown cylinder represents the expected position of ssDNA based on the DnaB-DnaC-ssDNA complex (PDB: 6QEM, (34)). The DnaB CTDs are depicted as grey spheres with labels corresponding to the underlying chain; the NTD domains are not shown for clarity. The DnaB DH and LH helices are colored in red and orange. This disposition of the amino-terminal domain (domains I/II) of λP chain Z effectively precludes entry of a physiological origin ssDNA into the central chamber of DnaB (highlighted by the molecular clash, which is circled in gray). B) The DnaB-DnaC-ssDNA complex (PDB: 6QEM) is posed as superimposed on chain A of the BP complex and depicted in the same style as the B_6_P_5_ complex in panel A. Individual DnaC protomers are colored in shades of gray. C) The B_6_P_5_ and B_6_C_6_ loader complexes in panels A and B are superimposed. Each of the C-terminal lassos from λP superimposes on a DnaC lasso. However, the single visible λP N-terminal lasso overlaps with the position of the DnaC lasso (chain G) at the top of the spiral (circled in red). Thus, one λP subunit (chain Z) forms contacts made by two DnaC loader subunits (chains G and L). D) Close-up of the red-circled segment in panel C.

Our 2.66 Å model of the B_6_P_5_ complex forces reconsideration of the mechanism of helicase loading. The finding that one λP protomer binds across the DnaB breach and blocks access of ssDNA to the inner chamber implies a previously unrecognized autoinhibited configuration. Our results predict that remodeling of the B_6_P_5_ complex to clear the path for ssDNA into the DnaB hexamer must accompany the recruitment of the B_6_P_5_ complex to the replication origin.

### The Pentameric λP Loader Ensemble Grips DnaB Tightly

In the B_6_P_5_ complex, the pentameric λP loader ensemble grips the DnaB helicase at five discrete and widely dispersed sites (Figure 5); each site comprises a DnaB-specific insertion into the RecA-fold (Supplementary Figure 17). Owing to the asymmetry of the complex, not every λP instance uses every site. The five sites on DnaB are 1) the DH element (residues 291:307) on chain B of the breached interface (its partner LH element from chain A is not visible in our maps); the N-terminal lasso of chain Z is the only λP element to make this contact (BSA: ∼1900 Å^2^). This contact explains the finding that deletion of λP residues 9:85 disrupts the complex (98). 2) A short DnaB helix comprised of residues 387:391 on the CTD at the breached interface; this contact is only made by domain II of λP chain Z (BSA: ∼400 Å^2^). 3) A wide patch that spans two adjacent DnaB CTDs (residues 424:434, 456 on one CTD and residues 391:395, 429:430, 446:449, and the C-terminal residue 468 from the adjacent CTD) and includes contacts (residues 447:450) on the arginine finger β-hairpin of each CTD; domain III of each λP monomer contacts this patch (BSA: ∼1600 Å^2^). The involvement of the extreme C-terminus of DnaB in λP contacts was anticipated by genetic experiments (99). 4) a site comprised of the DH (residues 290:307) and LH (residues 182:199) elements of two adjacent DnaB subunits; the C-terminal lasso of each λP monomer makes this contact (BSA: ∼1400 Å^2^). Notably, the nature of the DnaB•λP interfaces at sites 1 (λP N-terminal lasso) and 4 (λP C-terminal lasso), both of which include DnaB DH elements, diverge considerably; indeed, the N-terminal lasso occupies the same space as that of the chain B LH element (Figure 4 and Supplementary Figure 16). 5) A patch on the NTD of DnaB (residues 76:77, 183:195); two λP’s (chain Y and chain W) make this contact (BSA: ∼350 Å^2^). Our structure provides a partial rationale for the finding that specific DnaB mutants (residues V256I/E426K and G338E/E426K (98)) exhibit resistance to inhibition by λP, as residue 426 is located at interfacial site #3. Residues 256 and 338 are packed against one another in the CTD but are distant from the B_6_P_5_ interface; we speculate that mutations at these sites disrupt the CTD’s structure. The λP domain IIs of four chains (V, W, X, and Y) make no contact with DnaB.

**Figure 5.**
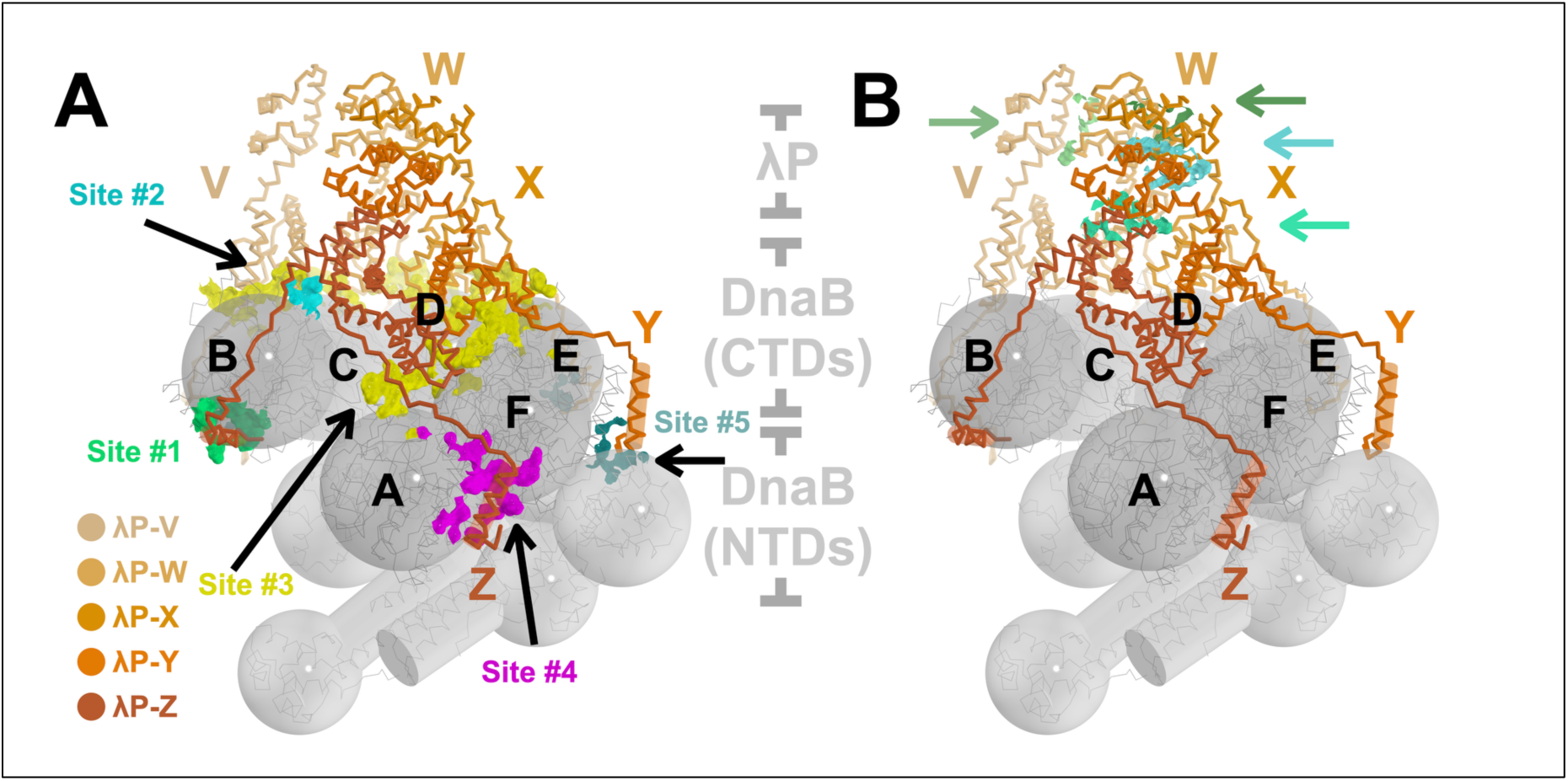
Five Widely Dispersed Sites Comprise the Interface between λP and *E. coli* DnaB. **A)** The five interaction sites (#1-#5, each uniquely colored) between the loader and helicase subunits. These interaction sites are depicted as surfaces, with points on the surface indicating positions within 4.5 Å between the helicase and the loader. Due to the asymmetry of the B_6_P_5_ complex, not every loader protomer utilizes each site. The λP ensemble is shown in ribbon format, colored in shades of orange as indicated in the legend. The DnaB hexamer, labeled from A to F, is presented in ribbon, sphere, and cylinder design language. Labels distinguish the CTD spheres. B) The B_6_P_5_ complex is depicted as in panel A, except that the interfaces between λP loader subunits are highlighted and indicated with arrows in the same color as the surface. Interaction sites between loader subunits are represented as surfaces, with points on the surface indicating positions within 4.5 Å between pairs of loader subunits.

The extensive interface between the λP ensemble and DnaB contrasts with the considerably more modest interface formed by the DnaC loader assembly (15, 33, 34). The six N-terminal lassos/grappling hooks of DnaC make identical interactions with the DH-LH interface of each DnaB subunit: this binding site overlaps with site #4 of the BP complex. Unlike domain III of λP, which slightly overlaps with the position of the AAA ATPase domains of DnaC, the latter domain makes no contact with DnaB. In this context, it is noteworthy that λP’s higher affinity to DnaB, which can displace DnaC from the BC complex (37), may be explained by the multitude of contacts to DnaB made by the λP ensemble, especially chain Z.

### The λP Ensemble Stabilizes the DnaB Spiral in the B_6_P_5_ Complex

An extensive interface between helicase and loader stabilizes the right-handed open DnaB spiral in the B_6_P_5_ complex (Figures 2, 3, 5, 6, and Supplementary Figure 15). Domain III of each λP loader contacts two consecutive DnaB subunits (interaction sites #2 and # 3). λP residues: 145:162 contact DnaB loops (residues: 424:432) on one subunit, while λP residues: 131:141 and 169:174 of the same chain contact a DnaB loop (residues: 391:398) and a strand (residues: 447:449, 454:456, and 468) on a neighboring subunit.

Although the DnaC and λP loaders are unrelated in structure, the DnaB portion of the B_6_C_6_ (34) and B_6_P_5_ (33) complexes adopts very similar right-handed open spiral configurations (15). However, the underlying arrangement of the CTDs is not identical; this divergence is reflected in the distinct helical parameters of each CTD spiral (Figure 3, B_6_P_5_: average helical pitch (12.6 Å), rise (3.7 Å), and twist (56.4°); B_6_C_6_: average helical pitch (15.8 Å), rise (4.2 Å), and twist (55.4°). This distinct architecture arises from wedging apart the two CTDs in each B_2_P_1_ sub-structure by ∼3° relative to the arrangement seen in the BC complex; this translates into small shifts in CTD positions (Figure 6). The disposition of DnaB subunits in the B_6_P_5_ complex directly results from the unique interface made with λP. The pentameric λP ensemble features an extensive interface with the helicase, and the contacts mediated by λP domain III mediate the wedging apart of DnaB subunits. On the other hand, the globular domains of the hexameric AAA+ DnaC ensemble make no contacts with DnaB whatsoever; the absence of contacts permits a closer approach of individual DnaB subunits to each other in the BC complex than is possible in the B_6_P_5_ ensemble.

**Figure 6.**
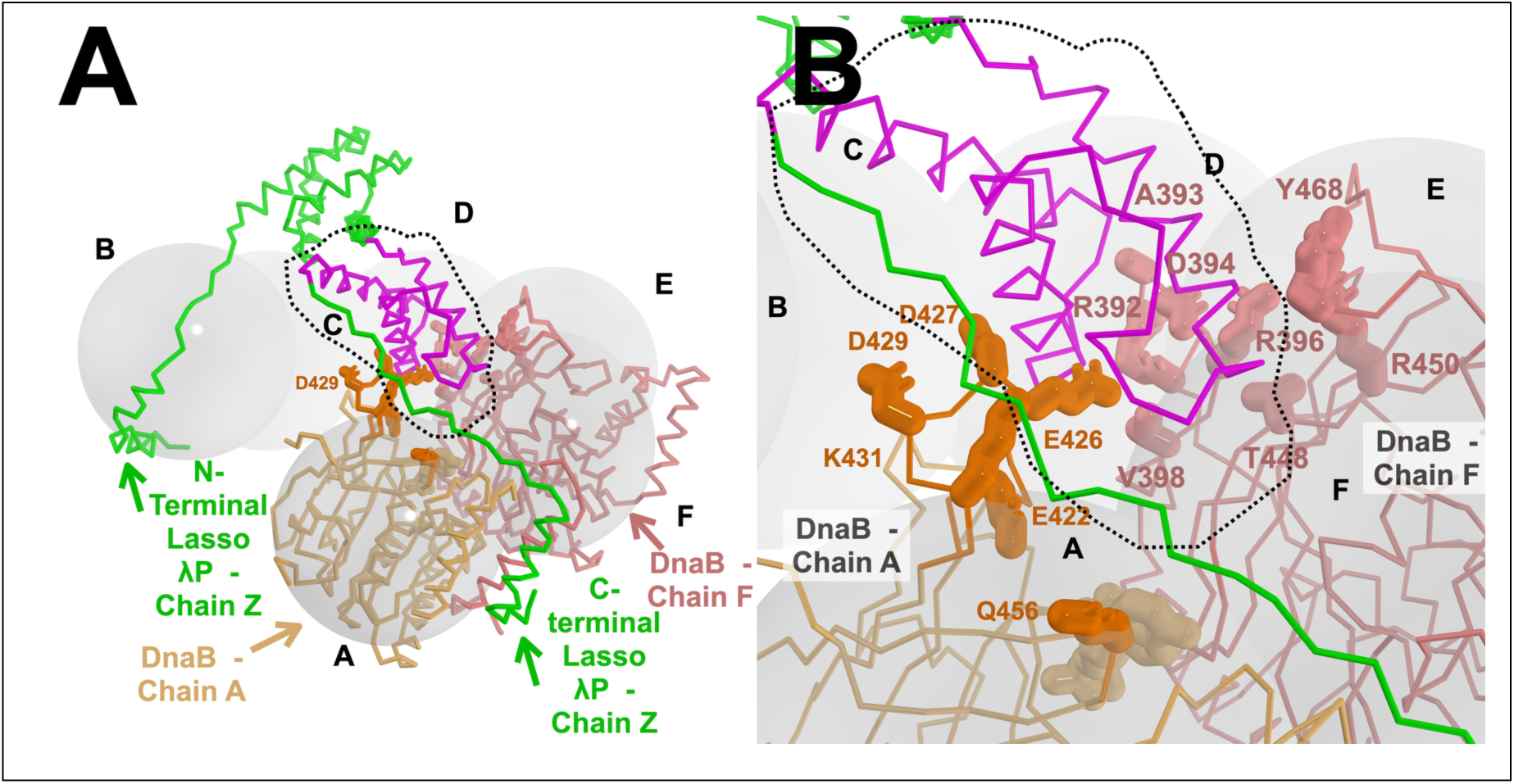
Stabilization of the DnaB Spiral by Domain III of λP in the B_6_P_5_ complex. A) The open spiral of DnaB in the B_6_P_5_ complex is stabilized through interactions with domain III of the λP loader. The CTDs of DnaB within the B_6_P_5_ complex are represented by transparent, labeled gray spheres. Chain Z of the λP ensemble (green) is depicted in the PyMol ribbon representation, along with the CTDs from chain A (light orange) and chain F (salmon). Residues in the DnaB CTD from chain A (light orange) and chain F (salmon) that interact with domain III of λP (outlined) are shown using the stick representation. The corresponding interfacial residues on chain λP Z are not displayed for clarity. The interface from the DnaB-DnaC complex (PDB: 6QEL, (34) and Supplementary Figure 15) does not exhibit such features. B) Close-up view of the area outlined in panel A.

The observation of similar helicase configurations in the B_6_P_5_ and B_6_C_6_ complexes led us to previously suggest that the open spiral might be an intrinsic DnaB state since it is observed in complexes with a loader that forms an extensive interface (λP) and one that does not (DnaC) (15, 33, 34). The 2.66 Å B_6_P_5_ complex structure teaches that the open spiral configuration may be intrinsic; however, the interaction with λP also drives changes in the spiral that are not seen when DnaC binds to DnaB. Whatever the source, an overall consequence of the inability of DnaB subunits in the B_6_P_5_ or B_6_C_6_ complexes to approach each other optimally has profound implications for both ATP hydrolysis and ssDNA binding (Supplementary Information and Supplementary Figures 13 and 18).

### The Carboxy-Terminal Lasso of the λP Ensemble Shears the LH-DH Interface of Each DnaB Monomer in the B_6_P_5_ Complex

During the transition from the closed planar to the open spiral form (B_6_P_5_), each C-terminal lasso helix of the λP ensemble binds alongside 5 DH - LH elements to create a three-helical bundle (interaction sites #4 and #5), binding shears and nearly completely disrupts the underlying DH-LH interface (Figures 5 and 7). Notably, the DH and LH elements are unique to the DnaB family of helicases compared to other RecA family members ((100), Supplementary Figure 17). As noted above, the N-terminal lasso of λP binds to the sixth DH element (chain B) in DnaB (top of the spiral as in the pose in Figure 2); this DH element lines the breach in the hexamer (interaction site #1). Binding wholly expels the partner LH element (chain A) since the N-terminal lasso-helix takes its place; indeed, the sixth chain A LH helix is not seen in our EM maps.

**Figure 7.**
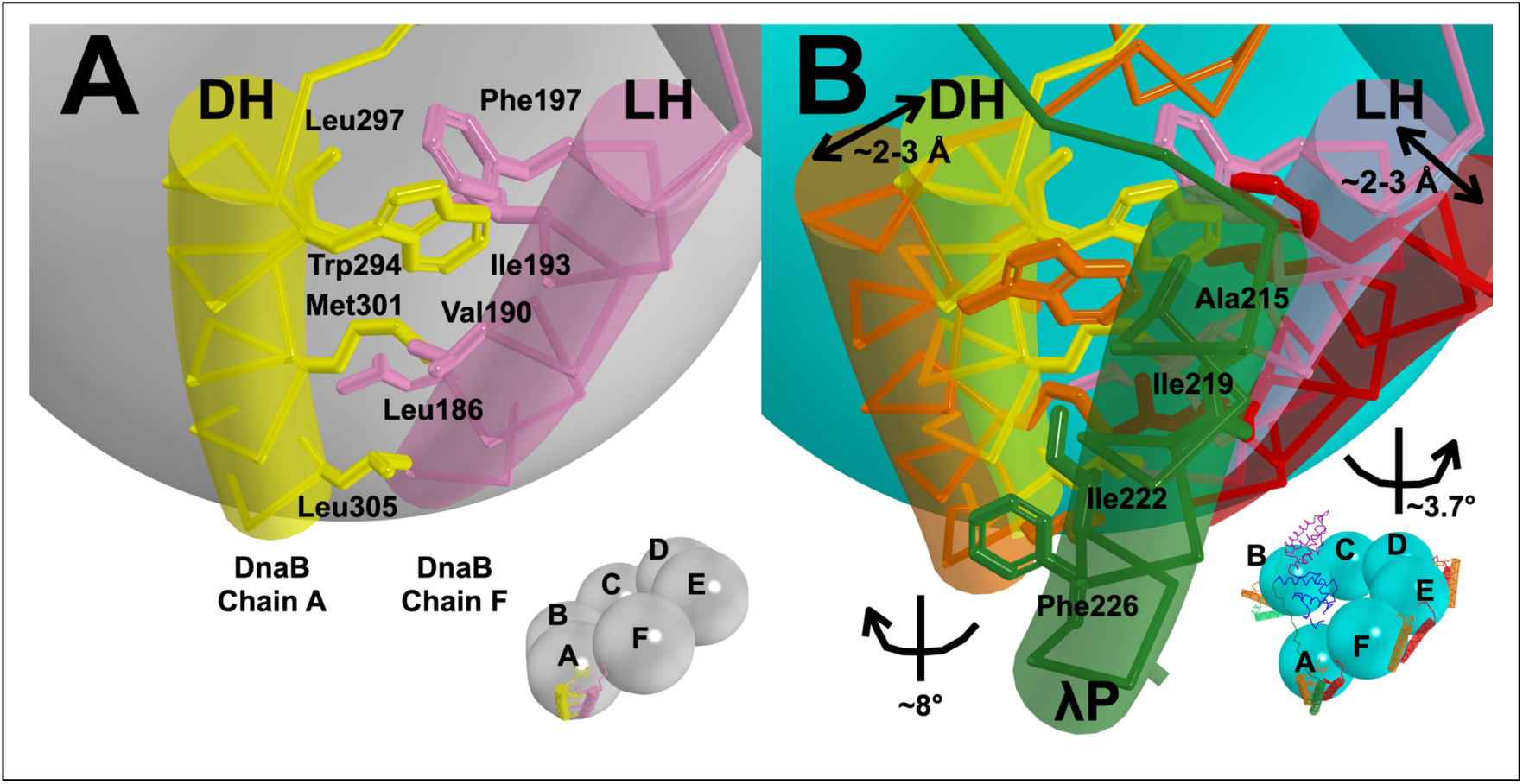
The Carboxy-Terminal Lassoes of the λP Ensemble in the B_6_P_5_ Complex Shear the DH-LH Elements. A) Knobs and holes packing between the docking helix (DH) and linker helix (LH) in the closed planar form of DnaB. The above model was constructed by superimposing DH and LH helices from the Alphafold model of *E. coli* DnaB (72) on the closed planar DnaB model (PDB: 4NMN, (12)). Residues 291:306 (DH) and 182:199 (LH) are depicted in the PyMol ribbon and transparent cylindrical cartoon representation. Interacting residues are shown as sticks and labeled. B) The C-terminal lasso of the λP loader disrupts the DH–LH interaction during the transition from the closed planar to the open spiral form of DnaB in the B_6_P_5_ complex. The DH and LH helices rotate an average of 8° and 3.7° away from the dimer interface as they move apart by 2-3 Å during the transition. The closed planar DH–LH interaction is disrupted and replaced by new interactions (residues shown as sticks and labeled) with the C-terminal lasso helix of λP. These interactions are found at each λP C-terminal lasso. Panel insets depict the relevant DnaB hexamer with each CTD drawn as a sphere; for clarity, the NTD domains are not shown. The letters on each inset represent protein chains of the DnaB hexamer.

In the closed planar form, the DH element of each DnaB subunit packs against the LH element of an adjacent monomer in an antiparallel fashion via an interface whose Cα atoms are, on average, ∼15.2 Å apart (the closed planar forms have the following values: 4NMN (constricted): 14.7 Å, 2R6D (dilated): 15.4 Å, 3BGW (constricted): 15.5 Å). In closed planar *E. coli* DnaB (modeled on the 4NMN structure (12)), the underlying contacts between the DH and LH elements are mediated primarily by hydrophobic interactions (Figure 7). For example, DH residue Trp 294 packs into a pocket formed by LH residues Val 190, Ile 193, and Phe 197 on an adjacent DnaB subunit. DH residue Met 301 packs in a pocket comprised of LH residues Leu 186 and Val 190, and DH residue Leu 305 binds in a pocket lined by Ala 183 and Leu 186.

Our prior work showed that the DH-LH interfaces in both the B_6_C_6_ (34) and the B_6_P_5_ complexes (15) are disrupted by the DnaC and λP loaders. In the 2.66 Å B_6_P_5_ structure, we observe that the helical segment (residues 211:227) of each C-terminal lasso of λP is positioned alongside the DH-LH elements. Compared to the closed planar form, the DH and LH helices are rotated by an average of +8.0° and ∼-3.7 °, respectively, and translated by ∼2.4 Å from ∼14.7 Å in the closed planar form to ∼17.1 Å in the λP-induced open spiral (Figure 7). A critical overall consequence of λP binding is the near-complete abrogation of interactions between the DH and LH elements in the DnaB open spiral. The repositioned DH and LH elements combine with the λP C-terminal lasso helix to produce a three-helix assembly; the disrupted DH-LH interface is replaced with distinct contacts between each component and residues in the λP C-terminal lasso helix. In the repositioned DH element, DnaB Trp 294 packs against λP residues Ala 215, Lys 218, and Ile 219; DH residue Met 301 resides next to λP residues Ile 222 and Phe 226; DH residue Leu 305 contacts λP residue Phe 226 as it maintains contacts with LH residue Ala183 (the sole contact between DH and LH in the loader complex). The repositioned DnaB LH elements also contact the λP C-terminal lasso helix. LH residue Glu 194 contacts Ile 219. Notably, the DH-LH-λP three-helix bundle interaction is nearly identical in all five instances of B_2_P_1_ in the B_6_P_5_ complex (DnaB CTD: RMSD = 0.2 Å on 200 Cα atoms, Supplementary Figure 14). We found that, although not included in the superposition, the second CTD DnaB in the B_2_P_1_ sub-structure and domain III/IV of the λP superimpose well; this suggests a relatively rigid DnaB-CTD_2_-λP substructure in B_6_P_5_ (Supplementary Figure 14).

Unlike the four λP protomers (chains V, W, X, and Y), which contact two DnaB subunits, chain Z contacts three DnaB chains (B, A, and F). The interaction with chain B at the top of the spiral (as in the pose in Figure 2) is mediated by the N-terminal lasso helix of λP, which is positioned adjacent to the DH element that lines the breach in the hexamer. In contrast to the three-helical DH-LH-λP bundle seen with the five instances of the B_2_P_1_ sub-structure, a two-helical bundle is observed (Figure 2 and Supplementary Figure 16). In this species, the LH element (chain A) has been displaced and is not visible in our maps; in its place, we find the λP N-terminal lasso helix. Notably, the same DnaB (chain B) DH residues highlighted above (Trp 294, Ile 297, Met 301, and Leu 305) participate in the divergent interaction with the N-terminal lasso helix of λP. DnaB DH residue Trp 294 contacts λP residue Met 16; DnaB residue Ile 297 packs against Gln 15; DnaB residue Met 301 packs into a pocket formed from λP residues Met 8, Phe 11, Asp 12. Finally, DnaB residue Leu 305 is found in a pocket formed by Met 8 and Val 9. The resolution of our maps is relatively low in the volume encompassed by site #1; as such, a description of the underlying interface should be considered tentative.

Examining the configuration of various DnaB structures suggests that the DH-LH interface is an essential nexus of structural changes during helicase loading. Measurements of the distances between DH and LH elements in the closed planar forms of Stage I (dilated: 2R6D (13) and constricted: 3BGW (11) and 4NMN (12)) reveal a ∼15.2 Å separation. In the Stage II DnaC loader bound conformers, with and without ssDNA, the distance between DH and LH elements increases to ∼17.4 Å and 17.8 Å; these values closely follow the ∼17.1 Å distance seen in the 2.66 Å B_6_P_5_ complex structure. With the expulsion of the loader and transition to the translocating Stage IV, the DH and LH elements move closer together (4ESV (18): 15.5 Å, 7T20:16.0 Å (101), 7T21:16.1 Å (102), 7T22: 16.1 Å (103), and 7T23: 16.2 Å (104)), but perhaps, not as close as the Stage I structures. Notably, in *V. cholerae* (VC), isolated (PDB: 6T66, (105)) and DciA-bound DnaB (PDB: 8A3V, (26)) diverge from the above trends. The average DH–LH distance in isolated closed planar VC DnaB is ∼17 Å, close to values extracted from loader-bound open spiral forms of DnaB. Furthermore, this distance appears to contract (15.6 Å) in the DciA-bound closed planar form of DnaB. Divergences in distances between DH–LH elements may suggest substantially different loader mechanisms between the DnaC/λP and the DciA helicase loading systems. Notably, the moderate resolution (2.8–3.8 Å) of the available structures limits our analysis.

Our work establishes the helical DH—LH DnaB elements as important moving parts in the loading pathway. The binding of the λP loader disrupts the hydrophobic packing seen in the DH—LH element of the closed planar configuration, yielding two new interfaces between the same surfaces of each component and the C-terminal lasso. A distinct interaction is seen between the DH elements and the λP N-terminal lasso.

### Where Does the B_6_P_5_ Complex Bind ssDNA?

DnaB and the λP loader are known to interact with ssDNA (18, 38, 106); however, the mechanism by which these contacts are integrated within the helicase loader complex remains unknown. We had previously shown that the ssDNA binding site on DnaB is disrupted compared to the translocating complex (15, 33) (Supplementary Information and Supplementary Figure 18). The 2.66 Å structure of the B_6_P_5_ complex teaches that not only is the ssDNA binding site disrupted in the loader complex, but two other architectural features sever access to the central chamber. First, as described above, one λP monomer (chain Z) binds to two DnaB subunits that span the breach, and in doing so, utilizes λP domains I/II to physically block the entry of ssDNA into the chamber. Second, owing to contacts between λP domain III and DnaB, the central chamber cannot achieve the active form while the loader remains bound (Supplementary Figure 19). As noted above, for DnaB in the B_6_P_5_ complex to achieve the configuration in the ssDNA complex, each CTD must rotate by ∼15° toward the central chamber. However, this structural change is precluded due to the steric clash between each λP domain III and pairs of DnaB CTDs in the ssDNA complex. This finding starkly contrasts with the B_6_C_6_ complex, where DnaC’s divergent binding mode places no restriction on the adoption by DnaB of the conformation in the ssDNA complex. Indeed, in the B_6_C_6_-ssDNA complex (PDB: 6QEM, (34)), DnaB adopts a configuration closely related to that in the ssDNA-bound translocating form (PDB: 4ESV, (18)). Notwithstanding its internal chamber’s distorted shape and geometry, the B_6_P_5_ complex is known to bind ssDNA robustly (Supplementary Figure 8 and (33, 38)).

Little is known about the binding modes that λP deploys to bind ssDNA. Although λP contains no known ssDNA binding domains, one λP residue (F99) has been shown to crosslink to ssDNA (38, 106). In the B_6_P_5_ complex, the five instances of λP F99 trace a winding path in the inner chamber formed by the λP domain IIs (Figure 8). This finding implies that, notwithstanding the restrictions to entry and binding of ssDNA in DnaB’s central chamber, the λP loader contacts nucleic acid in its inner chamber at some point during the helicase loading pathway. Since linear ssDNA was used for crosslinking, little light is shed on the process of loading onto the initiator-produced bubble at a replication origin.

**Figure 8.**
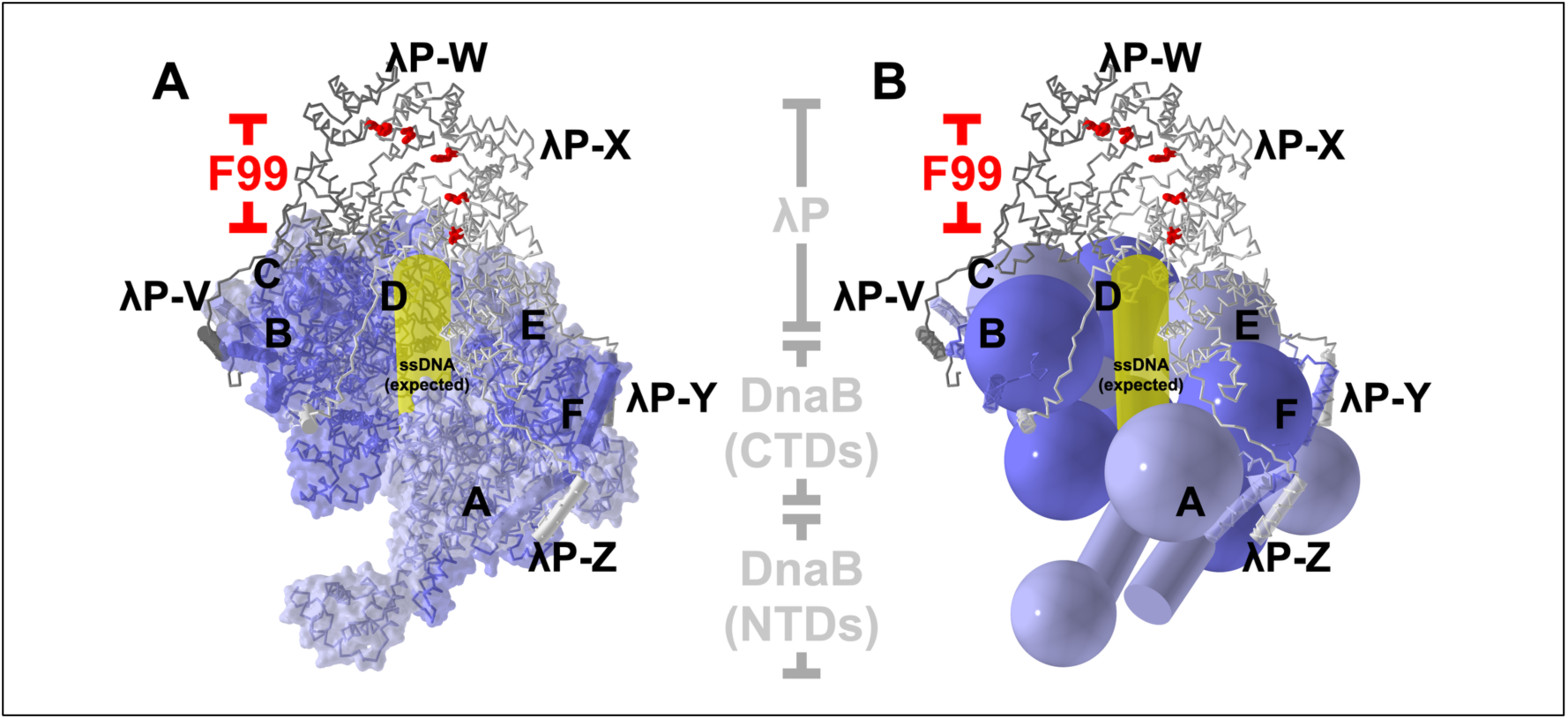
Path of ssDNA through the B_6_P_5_ complex. λP Phe 99 (red) is known to crosslink with ssDNA (38, 106). Within the B_6_P_5_ complex, the five instances of λP Phe 99 circumscribe a volume enclosed by the five copies of λP domain II. The constellation of λP Phe 99 residues is above the volume anticipated to be occupied by ssDNA (yellow), based on the DnaB•DnaC-ssDNA complex (PDB: 6QEM (34)). A) The B_6_P_5_ complex is illustrated in a ribbon representation. DnaB is shown with a transparent surface colored in alternating blue hues. B) The same orientation of the B_6_P_5_ complex, but with DnaB depicted using the sphere and cylinder design language.

Our structural findings raise several intriguing possibilities: 1) the λP loader, while bound to DnaB, contributes most, if not all, of the significant contacts to ssDNA, and 2) ssDNA is bound by DnaB but not in the central chamber. Such a distinct ssDNA binding mode is hinted at by the blocked entry path into the central chamber of the B_6_P_5_ complex and its distorted central chamber shape. Of course, the B_6_P_5_-ssDNA complex may blend both modes. Supporting the notion that DnaB interacts with surfaces outside the central chamber during the initial ssDNA encounter complex, prior work by Trakselis and colleagues provides ample evidence for the existence of such contacts (107, 108). Clarifying the path of ssDNA through the B_6_P_5_ complex and whether it changes during the loading reaction is an urgent priority for future studies.

### Six Copies of λP Program a DnaB Intermediate between Closed Planar and Open Spiral

In the B_6_P_6_ assembly, the DnaB and the λP loader components adopt significantly different arrangements than in B_6_P_5_ (Figure 9). DnaB exhibits two main configurational changes relative to its counterpart in B_6_P_5_. First, in contrast to the open spiral in B_6_P_5_, the CTD tier in B_6_P_6_ is essentially planar (B_6_P_6_: average helical pitch (∼2.0 Å), rise (∼0.5 Å), and twist (∼60°) (Figure 3). Moreover, although individual CTDs are ∼2 Å further apart in B_6_P_6_ (average = 35.2 +/- 0.8 Å) than in B_6_P_5_ (average = 33.1 +/- 0.1 Å), the CTD layer features no breach (Figure 9). We refer to this configuration as the ‘ajar planar’ configuration. The higher standard deviation in the distance measurement implies that DnaB in B_6_P_6_ is transitioning between the relatively rigid closed planar and open spiral forms (Supplementary Figure 20). Second, the NTD tier in B_6_P_6_ is found in the closed planar dilated configuration (Figure 9), in contrast to the constricted arrangement in B_6_P_5_. The NTD tier from B_6_P_6_ resembles those from 2R5U (109) (root mean square deviation (RMSD) = ∼2.0 Å) and 7T20 (101) (RMSD = ∼1.0 Å).

**Figure 9.**
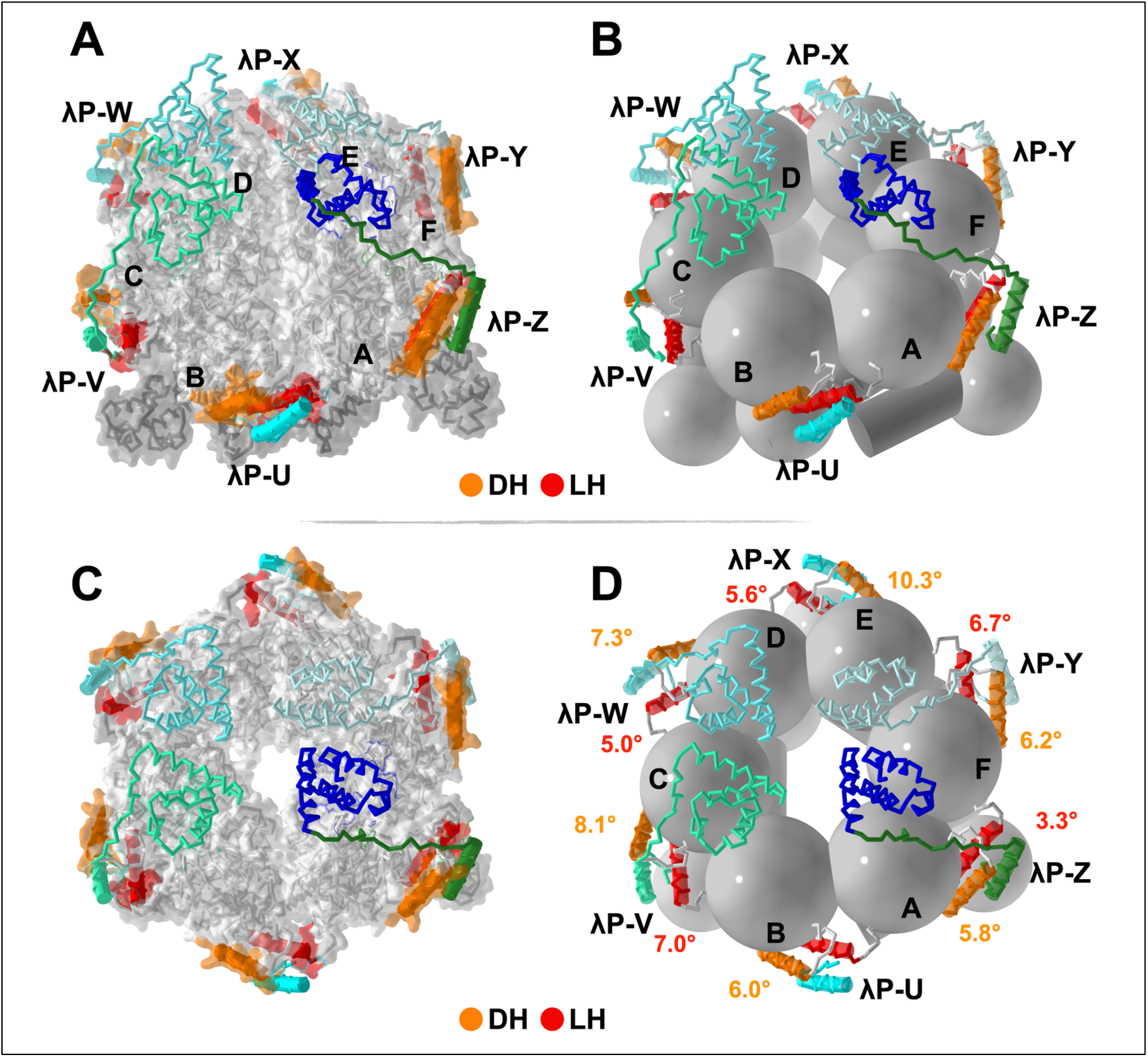
The Structure of the B_6_P_6_ Form of the *E. coli* DnaB–λP loader. In all the panels, the B_6_P_6_ complex, with its six loader molecules (labeled U, V, W, X, Y, and Z), is depicted in the ribbon representation; domains III and IV of λP chain Z are colored in blue and green; the other chains in hues of blue. DnaB in panels A and C is shown as a surface. Panels B and D illustrate DnaB using the sphere and cylinder design language. The DH and LH elements in each panel are shown in the ribbon and cylinder format and are colored orange and yellow. Panels A and B are rotated by 45° along the horizontal X-axis compared to the pose in panels C and D. The orange and red values in panel D represent rotation angles experienced by the DH and LH elements in the transition from closed planar to ajar planar.

The ajar planar form of DnaB in B_6_P_6_, like B_6_P_5_, exhibits distortions in the interfacial nucleotide binding sites and the ssDNA binding site in the internal central chamber (Supplementary Figures 13 and 18). The six nucleotide binding sites in B_6_P_6_ are filled with ADP/Mg^2+^, although the occupancy of the sites on chains B, C, and D is lower than that of the other chains (Supplementary Figure 12). As noted above, pairs of DnaB subunits are further apart, a finding upheld by the average distance between the Walker A lysine from one subunit and the arginine finger β-hairpin from the adjacent subunit (B_6_P_6_: ∼15 Å; B_6_P_5_: ∼12 Å; 4ESV: 11.4 Å). Therefore, the interfacial NTP sites on B_6_P_6,_ like those on B_6_P_5_, are not positioned appropriately for hydrolysis (Supplementary Figure 13). Indeed, our preparations of the BP complex, a mixture of B_6_P_4_, B_6_P_5_, and B_6_P_6_ (33), displayed no detectable traces of ATP hydrolytic activities (Supplementary Figure 21).

Compared to their positions in the closed planar, open spiral, and closed spiral ssDNA complexes, the seven DnaB residues in contact with ssDNA are significantly displaced in B_6_P_6_ (Supplementary Figure 18). The distances between one of these positions (*E. coli* Arg 403) in B_6_P_6_ and the corresponding residue (*B.st* 4ESV: Arg 381) in the translocating form are shifted between 6.3 Å and 39.1 Å (Supplementary Figure 18B). Like the B_6_P_5_ ensemble, the ssDNA binding surface in B_6_P_6_ diverges substantially from its configuration in the ssDNA complex.

### The Hexameric λP-DnaB Complex is Inchoate

The six copies of λP in the B_6_P_6_ complex (Figures 9 and 10) are differentially ordered in our EM maps. Domains I and II of each chain are entirely disordered. Only domains III and IV of the four copies (chains Z, Y, W, and V) are visible in our maps. The arrangement of these two domains in B_6_P_5_ is nearly identical to that in the five copies of λP in B_6_P_5_. For chains U and X, only the lasso domain IV is visible. The ajar planar DnaB configuration in B_6_P_6_ explains the pattern of ordered and disordered domain IIIs. The chain U λP domain III is disordered in our maps, and efforts to position it by superposition reveal a steric clash with corresponding domains from chain V and chain Z; similarly for chain X, which would clash with chains W and Y (Figure 10 and Supplementary Figure 22). These clashes arise from the planar nature of DnaB in B_6_P_6,_ which restricts the space available to accommodate the domains III seen in B_6_P_5_. We also speculate that disorder in the λP protomers may indicate that the loader ensemble is inchoate and has not yet formed the stabilizing interfaces seen in the B_6_P_5_ entity.

**Figure 10.**
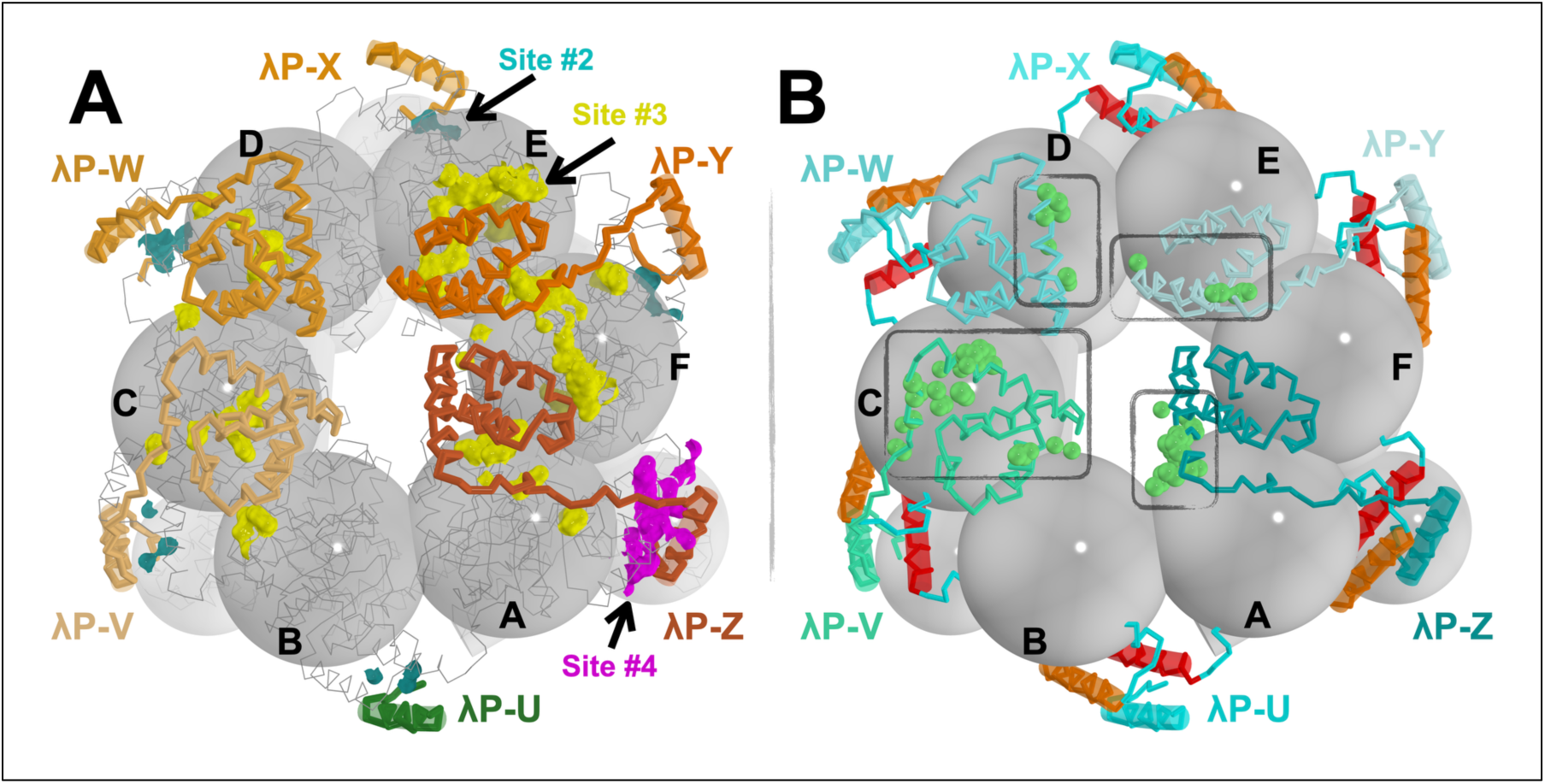
The Helicase Loader Interface in the B_6_P_6_ Complex is Inchoate. **A)** The interaction sites (#2, #3, and #4, each uniquely colored) between DnaB and λP. As in Figure 5, the interaction sites are depicted as surfaces, with points on the surface indicating positions within 4.5 Å between the helicase and the loader. Sites #3 (yellow) and #4 (purple) encompass different amounts of buried surface area in the various interfaces. The λP ensemble is shown in ribbon format, colored in shades of orange. The DnaB hexamer, labeled from A to F, is presented in a ribbon and the sphere/cylinder design language. The CTD spheres of each subunit carry labels corresponding to the underlying chain. B) Incompatibility of the disordered domain IIIs of chain U and X with the B_6_P_6_ complex is indicated by the clash (close approach; within 1.5 Å) with chains W and Y, and V and Z. The light-green-colored spheres show atomic positions in the above chains within 1.5 Å of chains U and X. The complete superposition appears in Supplementary Figure 23.

The disorganized state of the λP protomers in B_6_P_6_ extends to the interfaces formed with DnaB (Figure 10A). First, the lack of a breach in the DnaB CTD layer obscures binding site #1 for the chain Z N-terminal lasso/domain I. The disordered domain IIs do not permit the population of site #2, and changes to the NTD layer in B_6_P_6_ also abolish site #5. Only λP interfaces at sites #3 and #4 are formed. Moreover, the inchoate nature of B_6_P_6_ programs the burial of diverse surface area depending on the λP protomer. The four ordered λP domain IIIs each bury less surface area than in B_6_P_5_ (B_6_P_6_: ∼1200 Å^2^ vs. B_6_P_5_: ∼1600 Å^2^), and the amounts buried vary by chain (Y > Z > V >W). Similarly, for the λP domain IV interfaces, chain Y buries more surface area (∼1430 Å^2^) than the other chains (W=V=Z: ∼1040 Å^2^, X: ∼620 Å^2^, U: ∼610 Å^2^), but still less, on average, than in B_6_P5 (B_6_P_6_: ∼1000 Å^2^ vs. B_6_P_5_: ∼1400 Å^2^). Both findings indicate a lack of complete engagement of the various λP chains with DnaB. Second, comparing the six B_2_P_1_ sub-complexes in B_6_P_6_ reveals a much less rigid arrangement than in B_6_P_5_ (Supplementary Figure 20). Unlike the corresponding superposition with B_6_P_5_ (Supplementary Figure 14), the DnaB chain excluded from the calculation shows a considerably larger RMSD (B_6_P_6_: 5.4 Å vs B_6_P_5_: 0.7 Å). Moreover, while the characteristic rotations/translations of the sheared DH and LH elements are evident in B_6_P_6_, the underlying values differ considerably between chains compared to B_6_P_5_ (Figure 9D). In considering the significance of these changes, the current 3.85 Å resolution of B_6_P_6_ should be kept in mind.

Taken together, analysis of the architecture of DnaB and λP, nucleotide, ssDNA binding sites, and helicase and loader interfaces suggests that B_6_P_6_ represents an early, partially open, inchoate intermediate that can be assigned to a position between the closed planar and open spiral forms in the reaction trajectory (Figure 11).

**Figure 11.**
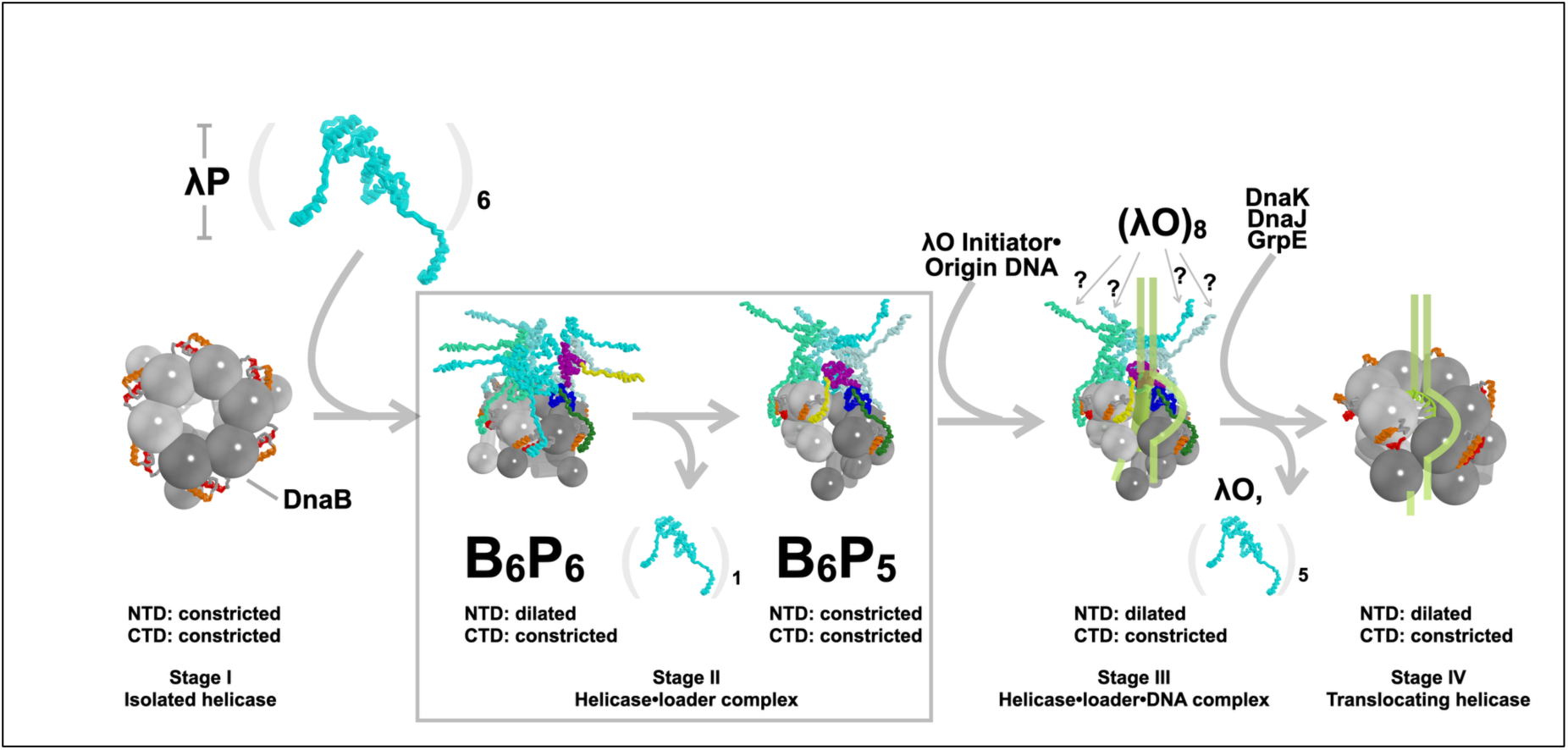
Model for loading the DnaB-λP loader complex onto the λO-Oriλ. Assembly stages of the DnaB helicase at the phage λ origin. This work focused on the two oligomeric forms of the BP complex (boxed) that populate Stage II: B_6_P_6_ and B_6_P_5._ Six copies of the λP loader bind to the closed planar ring (Stage I) of *E. coli* DnaB, creating the ajar-planar B_6_P_6_ complex. In this complex, the λP domains I (chain Z), II, and III have not adopted the configuration in B_6_P_5_. Loss of one copy of the λP loader accompanies reconfiguration into the B_6_P_5_ open spiral. This entity features openings in the NTD and CTD layers that allow ssDNA to enter the central chamber. However, the λP chain-Z binds on both sides of the breached interface in DnaB, stabilized by a substantial interface between the domain IIs of the λP pentamer. This configuration effectively blocks the entry of replication-origin-derived ssDNA into DnaB’s inner chamber. Despite the blockage, the B_6_P_5_ complex can still bind ssDNA; it is suggested that this binding may occur outside of DnaB’s inner chamber in Stage III. The disordered domain Is of λP chains V, W, X, and Y are proposed to bind to the λO-Oriλ initiator-replication origin complex. The expulsion of the λP ensemble, assisted by *E. coli* chaperone proteins, facilitates the transition to Stage IV, the translocating form. The yellow cylinder represents the expected path of ssDNA inferred from the DnaB•DnaC-ssDNA complex (PDB: 6QEM, (34)). The docking helix and linker helix elements are colored red and orange.

## Discussion

The mechanisms accompanying the loading and activation of the bacterial DnaB helicase at a replication origin are not yet fully understood. To gain insights into these mechanisms, we determined cryo-EM structures of the *E. coli* DnaB• λP helicase loader complex in two oligomeric states, B_6_P_5_ and B_6_P_6_. The B_6_P_5_ complex extends an earlier effort at a much lower resolution (33) by shedding light on structural elements that undergo conformation switching during helicase ring opening. The configuration of the pentameric λP ensemble in B_6_P_5_ also hints at a previously unrecognized autoinhibited intermediate for ssDNA loading. In contrast to the open spiral B_6_P_5_ entity, the B_6_P_6_ complex is essentially planar, but with the DnaB CTDs further apart; we refer to this arrangement as the ‘ajar planar’ configuration. The most economical scheme to sequence these states in the helicase loading pathway is to position them in the following order: a) closed planar DnaB, b) ajar planar B_6_P_6_, and c) open spiral B_6_P_5_, followed by other steps (Figure 11). We discuss these states in this order.

Closed planar DnaB exists in two conformational states, with the NTD and CTD tiers in either dilated or constricted configurations (reviewed in (9, 10, 14–17) and in Supplementary Information), with evidence for a hybrid state (NTD: dilated, CTD: constricted (102)). We observed that six copies of the λP helicase loader bind to closed planar DnaB, stabilizing its CTD tier in the constricted configuration (Figure 11). In contrast, the NTD tier remains dilated. In B_6_P_6_, both tiers are planar. However, the DnaB CTDs are positioned further apart than in the closed planar form, but not enough to create a breach that exposes the central chamber. The NTD tier is closed. The lack of a breach in ajar planar B_6_P_6_ DnaB explains the disordered λP N-terminal lasso, whose binding site, which only forms in the open spiral, is blocked (Supplementary Movies 1 and 2). Notably, the crystal structure of the *Bacillus* DnaB-family helicase bound to its DnaI helicase loader also features a 6:6 stoichiometry and a partially open planar DnaB species (110); the relationship between the two complexes, if any, remains to be elucidated.

Several lines of evidence suggest that the DnaB and λP components in B_6_P_6_ are still developing and have not reached their final form. First, the six λP protomers exhibit varying order levels in our maps. Each has an ordered C-terminal lasso domain, but two chains are missing domain III, and all chains lack the λP ensemble-stabilizing domain II found in B_6_P_5_ and the N-terminal lasso/domain I. Six C-terminal lassoes are ordered in B_6_P_6,_ indicating that this segment is likely the first to bind stably to DnaB. Second, the nexus between the DnaB DH and LH elements, representing the binding sites for the C-terminal lassos, exhibits the characteristic rotations/translations seen in the well-defined B_6_P_5_ entity, but these changes vary by subunit. Moreover, this finding suggests that disruption of the DH-LH interface precedes the breach of one interface and the transition to the open spiral observed in B_6_P_5_. Third, four of the six λPs in B_6_P_6_ feature ordered domain IIIs bound at CTDs from consecutive DnaB subunits. However, the nature, number of contacts, and buried surface area vary by binding site. Concomitantly, individual CTDs display greater configurational flexibility than in B_6_P_5_. We presume that the two disordered domain IIIs populate diverse conformations since binding to DnaB is precluded by steric clashes with the ordered domain IIIs. Fourth, the extensive domain II interface in the B_6_P_5_ λP ensemble cannot be accommodated in B_6_P_6,_ leading us to conclude that these interfaces are unformed and the underlying domain IIs adopt flexible conformations. Lastly, mismatches in spiral configurations between B_6_P_6_ and the translocating open spiral (PDB = 4ESV, (18)) suggest undeveloped ssDNA and nucleotide-hydrolytic sites.

Transition to the B_6_P_5_ open spiral complex requires considerable remodeling of the DnaB and λP entities (Supplementary Movies 1 and 2) and eviction of one of the λP chains. Both the NTD and CTD tiers are reconfigured. The planar NTD tier is breached and assumes an open spiral configuration. The ajar planar CTD tier is remodeled into an open spiral. Remodeling both tiers creates the complete set of five sites on DnaB for contacts with λP. Breach of the DnaB ring reveals λP binding site #1. Reconfiguring the DnaB CTDs optimizes sites #2 and #3 for binding, opening space for two additional λP domain IIIs to bind. The λP C-terminal lassoes engage more fully with open spiral DnaB at site #4. Remodeling of the NTD and CTD tiers creates site #5. Although there is room in the pentameric λP ensemble for a sixth domain II to engage, its corresponding domain III would have no binding site since binding occurs at DnaB subunit interfaces, and all the interfaces are occupied. As such, the chain Z N-terminal lasso binds near the DH helix on the DnaB CTD, where the sixth C-terminal lasso (chain U) might have been expected to sit. Notably, λP chain Z’s multivalent contacts could also block non-productive DnaB reclosure to the closed planar form. The finding that the λP loader harbors lasso elements at both the amino and carboxy termini creates conceptual links with homologs of the DciA loader, which also feature similar architectures at both termini (7, 27); however, a mechanistic relationship between λP and these DciA loaders remains to be established. Remodeling the DnaB CTD tier into the open spiral destroys the DH-LH binding site for the chain U C-terminal lasso. We speculate that the lack of binding sites for chain U domains III and the C-terminal lasso destabilizes its presence in the complex, causing it to be evicted as B_6_P_6_ matures into B_6_P_5_.

Although DnaB in B_6_P_5_ exhibits openings in the NTD and CTD tiers of sufficient size, the combination of contacts by the N-terminal and C-terminal lasso domains of λP-chain Z and the extensive interface formed by the five λP domain IIs precludes entry of physiological origin-derived ssDNA into the inner chamber. Deepening the mystery is the finding that the B_6_P_5_ complex still binds ssDNA. We speculate that ssDNA may bind to the λP ensemble (38) or to the exterior of DnaB, as previously suggested (107, 108), or both, before the necessary conformational changes that enable loading of ssDNA in DnaB’s inner chamber.

Prior studies have shown that B_6_P_5_ must undergo additional remodeling as the helicase loading and activation reaction progresses. The complex must engage with the octameric λO initiator protein – Oriλ DNA complex (40, 41). We speculate that four disordered λP N-terminal lassoes (chains V, W, X, and Y) may have a role in engaging with the λO ensemble, where each lasso engages with two λO subunits in the λO-λP-DnaB complex. We speculate that the autoinhibition will be relieved as the B_6_P_5_ complex is recruited to the λO•Oriλ complex at the replication origin (Figure 9) and may require intervention by the bacterial heat shock/chaperone proteins: DnaK, DnaJ, and GrpE (45–48). Among other questions to be addressed are: Does ssDNA populate a secondary binding site on the surface of DnaB before entering the central chamber? How and when does ssDNA enter the central chamber of DnaB? How is the λO-λP-DnaB complex further reconfigured to activate the helicase for unwinding? The critical roles played by the *E. coli* chaperones DnaK, DnaJ, and GrpE in additional remodeling and eviction events remain to be clarified.

## Supporting information

Supplementary sections

## Data Availability

Coordinates from the X-ray structure of λP – domain III and the cryo-EM structure of the *E. coli* DnaB–λP complex are available from the RCSB Protein Data Bank under accession codes 8V9S (λP – domain III), 8V9T (B_6_P_5_-2.84 Å), 9OA1 (B_6_P_5_-2.66 Å), and 9AO2 (B_6_P_6_-3.85 Å). The underlying primary data, X-ray detector diffraction images, structure factors, and cryo-EM maps are available from the Protein Data Bank, the SBGrid Data Bank, and the EMBD under the accession codes EMD-43086 (B_6_P_5_-2.84 Å), EMD-70269 (B_6_P_5_-2.66 Å), and EMD-70271 (B_6_P_6_-3.85 Å).

## Author Contributions

DB, AS, and DJ conceptualized the study. DB and AS designed and prepared all the proteins. DB and EI performed crystallization and X-ray crystallography. AS performed all the ATPase assays. AS and JC acquired and analyzed the cryo-EM data. All nMS analyses were performed by PDBO in the BTC lab. DB, AS, JC, and DJ jointly built and analyzed the structures and drafted the manuscript.

## Acknowledgments

We thank the Jeruzalmi lab members, the biophysics group at City College, the CUNY Advanced Science Research Center, and the New York Structural Biology Center for scientific and technical advice. We dedicate our work on the DnaB–λP complex to the memory of Professor Roger McMacken, on whose shoulders we stand and who inspired us in so many ways.

