## Supplementary sections for "Distinct Quaternary States, Intermediates, and Autoinhibition During Loading of the DnaB-Replicative Helicase by the Phage λP Helicase Loader"

### Supplementary Information

#### Structure of the DnaB Replicative Helicase

Hexameric *E. coli* DnaB is a two-tiered, ring-shaped ensemble with an internal chamber where ssDNA binds during translocation (1–3). Each DnaB protomer is formed out of two domains: an amino-terminal domain (NTD) and a carboxy-terminal domain (CTD); the NTD and CTD of each DnaB subunit are linked by a linker helix (LH) element. In the assembled hexamer, pairs of amino-terminal domains engage in a dimeric arrangement, and three of these dimers assemble into a trimeric configuration that represents one of DnaB's layers; the NTD layer circumscribes the DNA binding channel and provides binding sites for DNA primase during the elongation phase of DNA replication. The second of DnaB's layers is formed out of six carboxy-terminal domains (CTD), which exhibit a pseudo-sixfold arrangement in contrast to the NTD layer. The CTD tier comprises six CTDs of DnaB, each of which harbors a RecA-like ATPase domain. Each CTD harbors a helical element, termed the docking helix (DH), which packs against the LH element of an adjacent CTD. DnaB, a superfamily 4 helicase, binds ATP at CTD interfaces; the Walker A and Walker B motifs for this composite site are furnished by one CTD and the arginine finger β-hairpin motif by the adjacent CTD. The residues that contact the phosphate backbone of ssDNA within the inner chamber of DnaB are also found on the CTD (4). It has been suggested that the interplay between the arginine finger-beta-hairpin motif and the ssDNA binding loops links ssDNA binding with ATP hydrolysis (4). The NTD and CTD layers are found in two arrangements, termed dilated and constricted, in addition to differently sized inner chambers. These two states feature distinct arrangements that give rise to divergent inter-subunit contacts (Supplementary Figure 1).

#### Schemes for helicase assembly on nucleic acid substrates

Three schemes for helicase loading have been described, termed here as (1) ring-opening, (2) ring-forming, or (3) ring-closing (5–7). These schemes are organized around the structure of the helicase before loading. In the ring-opening scheme, the hexameric helicase exhibits a closed ring configuration. During helicase loading, one of the six interfaces is breached to permit the entry of ssDNA into the internal chamber. The *E. coli* DnaC and bacteriophage λP loaders (8–10) implement this scheme. In the ring-forming scheme, DnaB subunits are assembled around DNA by the *B. subtilis* DnaI loader into the active helicase (11). The ring-closing scheme is observed with the Rho and MCM2-7 helicases, wherein the hexameric rings are pre-opened; ring closure occurs upon the entry of the nucleic acid substrate into the inner chamber and involves additional factors (6, 12, 13).

#### The Complete Structure of the λP Helicase Loader

To enhance our previous analysis of the BP complex (8), we employed protein engineering, experimental structural analysis, and AlphaFold structure prediction (14, 15) to determine the complete structure of λP. Using the structure of λP within the BP complex (PDB = 6BBM, (8)) as a guide, we isolated a soluble fragment of λP (residues 105 to 210, domain III). We next determined its X-ray crystal structure (residues 119 to 192) at 1.86 Å (Supplementary Figure 10 and Supplementary Table 2). λP domain III adopts a three-helical structure resembling that seen in the earlier cryo-EM analysis; further, the crystal structure confirmed prior inferences about λP's protein chain direction and the approximate side chain mappings to the structure (8).

We also calculated the complete model of λP with the Google CoLab implementation of AlphaFold in ChimeraX (14, 15). AlphaFold provided five predicted conformers of λP; each conformer harbored four domains: two compact domains (domain II: residues ~30 to ~119; domain III: residues ~120 to ~192) and two flexibly linked structural domains at the amino (residues 1 to ~29) and carboxy (193 to 233) termini with a single alpha helix, termed the N-terminal and C-terminal lasso/grappling hooks. The AlphaFold-predicted structure was used to interpret our cryo-EM maps. The crystal structure of λP domain III superimposed well on the AlphaFold predicted structure (root mean square deviation (RMSD) of 0.328 Å over 478 atoms). AlphaFold predicted a four helical bundle for the amino-terminal domain (domain I/II) that terminated in a lasso-like amino-terminal helix. Notably, the diverse configuration of the AlphaFold predicted conformers hints at flexibility in the arrangement of the four domains.

##### The 2.84 Å Cryo-EM Structure of the *E. coli* DnaB–λP Helicase Loader

This work aimed to obtain a cryo-EM (16–19) structure of the $B_6P_5$ complex bound to ssDNA. Although the results of native mass spectrometry analysis revealed the clear presence of a $B_6P_5$-ssDNA species (8) and are described below, the 2.84 Å map we obtained did not reveal the position of the ssDNA. Although devoid of ssDNA, our EM maps had a higher resolution than our prior study (8). The $B_6P_5$ 2.84 Å model is very similar to the 2.66 Å model except that it is less complete; four chains (V, W, X, and Y) lack density for a λP domain II. As such, our description here is limited.

#### Nucleotide Hydrolysis by DnaB is Directly Suppressed by λP in $B_6P_5$.

Although the sample submitted to cryo-EM analysis was supplemented with an excess of ATP (0.2 mM), each site in the $B_6P_5$ 2.66 Å model was filled with ADP. In the $B_6P_5$ 2.84 Å model, only 5 of the 6 sites on DnaB in the complex were found to be filled with ADP (Supplementary Figure 12A); the nucleotide site on the subunit that lines the breach (chain B) remains unfilled. In the sites filled with ADP, in addition to a pocket formed out of the conserved GKT (residues 236-238) Walker A and Walker B (residue D343) motifs, DnaB residues Gln281, Thr282, and Arg285 surround the six amino group of the adenine group, with the potential to form weak hydrogen bonds (~3.8 Å), however, essentially all these interactions could also be made by the 6 oxo group of guanine; the smaller pyrimidine bases could not form these interactions. There are no contacts to the ribose sugar. As such, the lack of specificity for ribonucleotide bases revealed by solution studies, including the lower affinity for purine ribonucleotides (20–22), is represented in our structure.

The open spiral configuration of DnaB in the $B_6P_5$ complex rearranges the composite ATP sites into configurations ill-suited for hydrolysis and/or ADP release (Supplementary Figure 13 and Figure 21). ATP binding and hydrolysis on DnaB take place at composite sites with structural elements that span two subunits; the Walker A/P-loop and Walker B elements are found on one subunit, and the β-hairpin arginine finger, which completes the site, resides on an adjacent subunit (4). Prior work ((23) and Supplementary Figure 21) found that DnaB's ATPase activity is entirely suppressed in the BP mixture of $B_6P_4$, $B_6P_5$, and $B_6P_6$. Each nucleotide binding site in the structure of the $B_6P_5$ complex is disrupted relative to the presumed active configuration in the translocating form (PDB: 4ESV, (4)). Inspection of the $B_2P_1$ superposition described above suggests that relative to the translocating form, the CTDs of DnaB in $B_6P_5$ are rotated outward away from the central chamber by ~16°. Notably, Lys *B.st* 440/*E. coli* 418 and *B.st* Arg 442/*E. coli* 420 are implicated in ATP hydrolysis in DnaB family members (24, 25). Changes in CTD configurations between $B_6P_5$ (R442) and translocating DnaB (R420) result in average shifts of 1.7 Å (Lys) and 1.4 Å (Arg) in the β-hairpin arginine finger sub-structure (Supplementary Figure 13). Shifts of this magnitude could explain the suppression of ATPase activity in the DnaB•λP complexes.

Outward rotation of the CTDs is directly linked to loader binding since domain III of λP makes contacts with two adjacent DnaB CTDs, including the arginine finger β-hairpin of each CTD (site #3), as noted above (Supplementary Figure 19). Close inspection of the superposition shows that the position of λP-domain III is incompatible with the position of the DnaB CTDs in the translocating form. As such, the λP loader stabilizes a DnaB configuration that precludes the formation of nucleotide binding sites competent for hydrolysis and/or ADP release and, thus, neatly accounts for the ATPase biochemical data ((23) and Supplementary Figure 21). Support for the idea that suppression of ATPase activity arises from shifts in the arginine finger β-hairpin and the direct role played by λP in specifying the shift is supported by comparisons with the analogous BC complex, which also lacks ATPase activity. The BC complex's architecture of the ATP sites resembles those seen in the $B_6P_5$ complex, although positional shifts in the arginine-finger β-hairpin are not as extensive (Supplementary Figure 13).

DnaC, which is unrelated in structure to λP, adopts a distinct architecture when bound to DnaB that completely lacks contact with the arginine-finger β-hairpin. The absence of these contacts may explain the modest shift of the β-hairpin in the BC complex. For the reasons presented above, we conclude that surfaces from domain III of λP promote and stabilize the shifted configuration of the arginine finger β-hairpin and, concomitantly, suppress ATP hydrolytic activity.

#### Distorted Geometry of the ssDNA Binding Sites in $B_6P_5$

We have previously described how the shape and geometry of the ssDNA binding site in the central chamber of DnaB differ significantly between the $B_6P_5$ complex (State II, (8)) and translocating states (State IV, PDB: 4ESV, (4)). In the translocating DnaB from *B.st*, each DnaB subunit projects a DNA binding loop into the central chamber to provide contacts by three residues (PDB: 4ESV; *G.st*: R381, E382, G384; *E. coli*: R403, E404, G406) per subunit to ssDNA. These contacts are also seen in the analogous *E. coli* complex (PDB: 7T20) but include four additional residues per subunit (*E. coli*: N356, T358, N386, R387). Thus, a total of 7 DnaB residues are in contact with ssDNA. In 4ESV, the pitch of DnaB's spiral matches ssDNA's, providing equivalent contacts between each helicase subunit and the nucleic acid backbone. However, the pitch and twist of the closed spiral DnaB (PDB: 4ESV, ~19.1 Å, ~60°) differ significantly from values (~12.6 Å, 56.4°) observed in the $B_6P_5$ complex. Divergently configured DnaB spirals critically displace the positions of ssDNA binding residues in the loader complex between the $B_6P_5$ 2.84 Å structure from values in the translocating form (A: 5.6 Å; B: 2.5 Å; C: 4.0 Å; D: 10.2 Å; E: 15.8 Å; F: 19.8 Å, and Supplementary Figure 18). A comparison of the 7T20 E. coli – ssDNA complex reveals similar shifts. Although the architecture of the ssDNA binding central chamber is altered in the $B_6P_5$ complex and might preclude ssDNA binding, it is formally possible that a different ssDNA binding mode may mediate binding.

### Supplementary Methods

#### Cryo-EM Image Processing of P11-J107 ($B_6P_5$-ssDNA, 2.84 Å)

To facilitate computational handling, the 25,595 movies were divided into four groups, designated as A, B, C, and D; identical computational procedures were applied to each group (Supplementary Figures 6 and 7, and Supplementary Table 3). The frames were aligned using motion correction and a B-factor of -500 Å$^2$, followed by the estimation of the Contrast Transfer Function (CTF) using the CTF fitting tool in WARP (26). The WARP program was also used to extract approximately 12 million particles from 25,959 frames using the BoxNet2Mask_20180918 deep convolutional neural network and the following parameters: diameter = 150 Å, box size = 320 pixels, pixel size = 1.083 Å, dose per frame = accumulated dose of 51.01 $e^-$/Å$^2$ / 40 frames = 1.28 $e^-$/Å$^2$ / frame. The WARP-extracted particles were imported into cryoSPARC (version 3.3.2) (27–29) and then subjected to 2D classification and *ab initio* reconstruction with C1 symmetry enforced. Each reconstruction produced 4 classes A0 to A3 (~360,000, ~360,000, ~790,000, ~480,000); B0 to B3 (~200,000, ~160,000, ~320,000, ~170,000); C0 to C3 (~370,000, ~720,000, ~490,000, ~310,000), D0 to D3 (~380,000, ~600,000, ~330,000, ~330,000 particles). Inspection of these 12 classes for high-resolution features and particle numbers suggested that the A2, B2, and C1 classes, which included ~1.83 million particles (790,000, 320,000, and 720,000 particles, respectively), be prioritized for template-based heterogeneous refinement. The class C3 volume, which did not resemble the $B_6P_5$ complex, was also included in the heterogeneous refinement procedure to exclude low-quality particles. Template-based heterogeneous refinement was performed using the published 4.1 Å EM density map (EMD-7076) (8) as the template. To avoid a calculation with 3 closely resembling volumes, we excluded the C1 volume owing to its similarity to the volumes in classes A2 and B2. The heterogeneous refinement step produced 4 classes: E0, E1, E2, and E3. Class E0 (Supplementary Figure 3S, red volume), which featured the most significant number of particles (~1.13 million) and provided the highest resolution map (4.41 Å), was carried forward to homogeneous refinement and non-uniform refinement. These procedures yielded EM maps with resolutions of 2.88 Å (Volume J, green in Supplementary Figure 6) and 2.84 Å (Volume K, purple in Supplementary Figure 6), respectively. The 2.84 Å map was considered to exhibit a higher overall quality. Inspection of both maps after sharpening with DeepEMhancer (30) upheld this judgment. The 2.84 Å EM sharpened map (Volume M, cyan in Supplementary Figure 6) represented our highest-quality map and was used to guide model building (below).

Although the map derived from the E1 class (Yellow in Supplementary Figure 6) produced a map with a similar resolution (4.41 Å) to that from the E0 class, we recovered a smaller number (~432,000) of particles. Nevertheless, we submitted this class to homogeneous refinement and obtained a map with a resolution of 3.62 Å (Volume L, magenta in Supplementary Figure 6). Model building into this map revealed a $B_6P_5$ complex that lacked density for the λP chain

Z domain II. Classes E2 (Gray in Supplementary Figure 6) and E3 (Orange in Supplementary Figure 6) harbored ~100,000 particles each, at 9 Å and 8.55 Å, respectively. The low quality of all these maps precluded meaningful analysis.

To evaluate potential biases in the above procedure, the A2, B2, and C1 classes, along with their corresponding volumes, and the C3 class volume (from above) were also subjected to heterogeneous refinement, but in this case, without a template. This procedure generated 4 classes (F0, F1, F2, and F3). We elected to proceed with the F0 class because it featured the highest resolution (4.41 Å) map and the largest number of particles (~1 million); homogeneous refinement of this class yielded a map with a resolution of 2.95 Å (Volume G in Supplementary Figure 6). A second effort encompassed classes F0, F1 (~380, 000 particles), and F2 (~260K particles), which featured volumes with resolutions of 4.41 Å (above), 4.41 Å, and 4.50 Å (the F3 class produced a volume with a resolution of ~9 Å; the ~110,000 particles in this class were not pursued in this study). The combined set of particles from the F0, F1, and F2 classes (1.7 million particles) was submitted to a second round of heterogeneous refinement. This procedure produced 4 classes (H0, H1, H2, and H3). The highest resolution volume (4.41 Å) and the most significant number of particles (~1 million) were recovered in class H0. Homogeneous refinement of these particles provided a 2.93 Å map (Volume I in Supplementary Figure 6). The 2.95 Å and 2.93 Å maps obtained from the non-template approach did not yield maps of higher quality than those obtained when a template was used in refinement. Classes H1, H2, and H3 harbored smaller numbers of particles, ~270,000, ~220,000, and ~175,000, respectively, and produced maps with resolutions 4.41 Å, 4.41 Å, and 6.74 Å, respectively. These maps, even after sharpening, did not reveal information not already seen in the high-quality 2.84 Å EM map described above.

The four classes recovered from group D (D0, D1, D2, and D3) were subjected to heterogeneous refinement, yielding four classes (N0, N1, N2, and N3). The N1 class, with the highest resolution volume (4.41 Å) and the most significant number of particles (~860,000), was moved forward; the other classes N0 (~6 Å with ~320,000), N2, and N3 (both at ~8 Å with ~230,000) particles were abandoned since they produced volumes with no resemblance to the BP complex. Homogeneous refinement, a second round of 2D classification, *ab initio* modeling, and heterogeneous refinement, applied to class N1, produced four classes (O0, O1, O2, and O3). The O2 class, with the highest resolution map (3.92 Å) and the largest number of particles (~450,000), surpassed the recovery in classes O0, O1, and O3 at resolutions below 4.1 Å, which each had approximately 100,000 particles. Consequently, O2 advanced to the next stage in the refinement step. To explore whether the O2 class would produce a higher-quality reconstruction, we combined the particles in O2 with those in K, G, and I to obtain 3.7 million particles. CryoSPARC then culled duplicate particles to yield a set of 1.7 million particles. This particle set underwent *ab initio* reconstruction, heterogeneous refinement, homogeneous refinement, and non-uniform refinements. Homogeneous and non-uniform refinement procedures produced two maps, each with a resolution of 2.81 Å (Volumes P and Q in Supplementary Figure 6).

Notwithstanding the resolution, the quality of these density maps was substantially lower than the ones described above.

The tilted dataset was processed in the same manner as described above, producing a final map with a resolution of 3.30 Å. Combining this map with the 2.84 Å described above did not yield a higher-quality map, so the tilted data set was not pursued further.

#### Model Building

**P11-J107 ($B_6P_5$, 2.84 Å).** Model building leveraged high-resolution crystal structures of the amino-terminal domain of *E. coli* DnaB (PDB: 1B79 (31)), domain III of the λP (1.86 Å, this work), as well as AlphaFold (14, 15) models of both *E. coli* DnaB (32) and λP. Components were placed using MOLREP (33, 34) or guided by an earlier $B_6P_5$ structure (PDB: 6BBM) (8).

The final model encompassed the following DnaB residues (chain A: 25:173 and 201:468, chain B: 16:468, chain C: 19:468, chain D: 18:468, chain E: 19-468, and chain F: 24:468) and λP residues (chain V: 119:232, chain W: 119:232, chain X: 119:232, chain Y: 119:232, and chain Z: 2:231). The model also included 5 ADP molecules bound to 5 magnesium ions in the five DnaB chains (A, C, D, E, and F); the nucleotide site corresponding to chain B is unfilled. The P11-J107 map showed no density that could be interpreted as ssDNA, even though it was included in the sample submitted to cryo-EM.

#### Refinement

**P11-J107 ($B_6P_5$, 2.84 Å).** The 2.84 Å model of the $B_6P_5$ complex was refined using the real_space_refine tool in Phenix against either maps produced by CryoSPARC or DeepEMhancer. Refinement encompassed cycles with the minimization global, NQH flips, and the atomic displacement parameters (adp) options, and leveraged the Ramachandran and secondary structure restraints. The rigid body option was applied only in the first round of refinement. The rigid bodies for all DnaB chains were defined as follows: residues 1:173 (NTD), 174:200 (LH), and 201:468 (CTD). For λP, the rigid bodies were defined as follows: residues:1:40 (domain I), 41:118 (domain II), 119:192 (domain III), and 193:233 (domain IV) and applied only when the domain in question was present in a particular model. We also performed two rounds of model re-building with high-resolution components, as described above, followed by re-refinement. This procedure produced high-quality models with excellent statistics, particularly in the less well-defined segments.

#### Supplementary Figures/Supplementary Legends

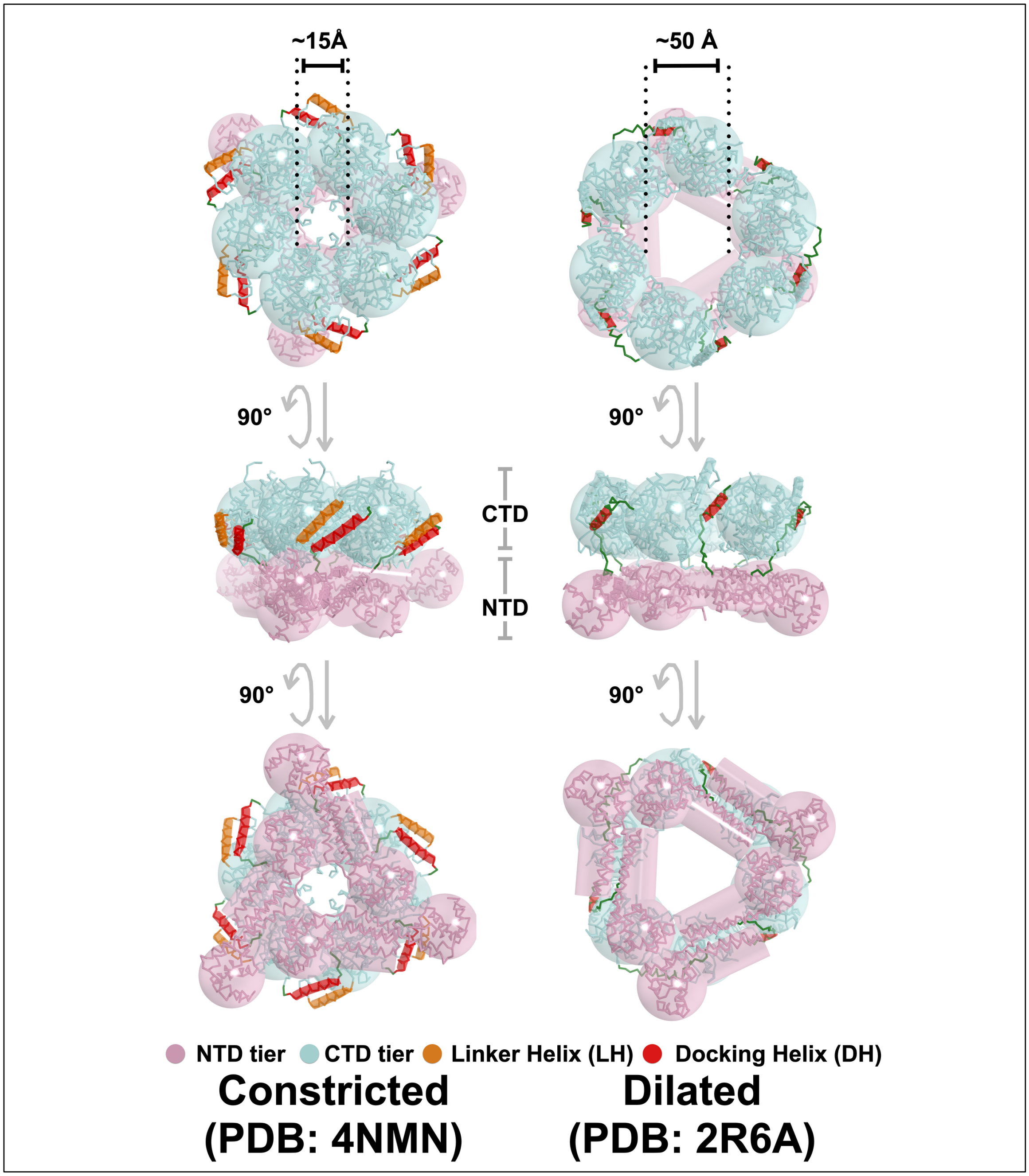

**Supplementary Figure 1. Architecture of the DnaB Replicative Helicase.** DnaB adopts distinct configurations known as constricted (PDB: 4NMN, left) and dilated (PDB: 2R6A, right) (1–3). The arrangement of both the NTD and CTD tiers differs between the two forms. Additionally, DnaB structures have been identified with a hybrid tier configuration, where the NTD is dilated while the CTD is constricted (35). DnaB is illustrated in a PyMol ribbon representation with superimposed sphere and cylinder design language elements. The NTD layers are pink, while the CTD layer is light blue. The DH and LH elements are presented in the PyMol ribbon and cylinder formats, colored red and orange.

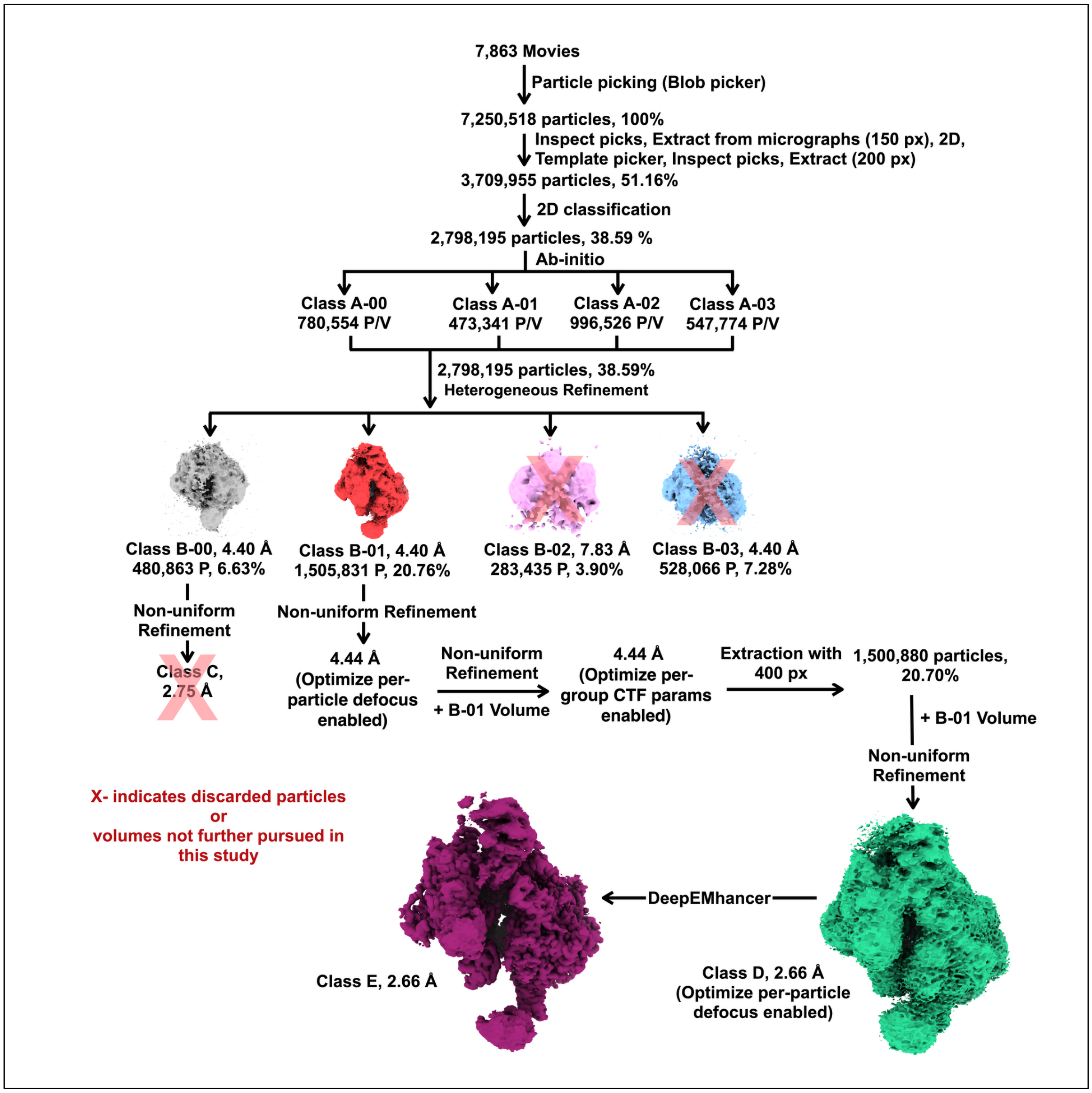

**Supplementary Figure 2. Processing of the P45-J50 Cryo-EM Data Set ($B_6P_5$, 2.66 Å).** The calculations are detailed in the Methods section. The volumes are distinguished by color. The 2.66 Å volume (green) represents the highest-resolution EM density map we obtained. Additional map improvement was achieved by processing with DeepEMhancer ((30), shown in a deep-purple-colored volume). The percentages are calculated based on the total number of particles picked.

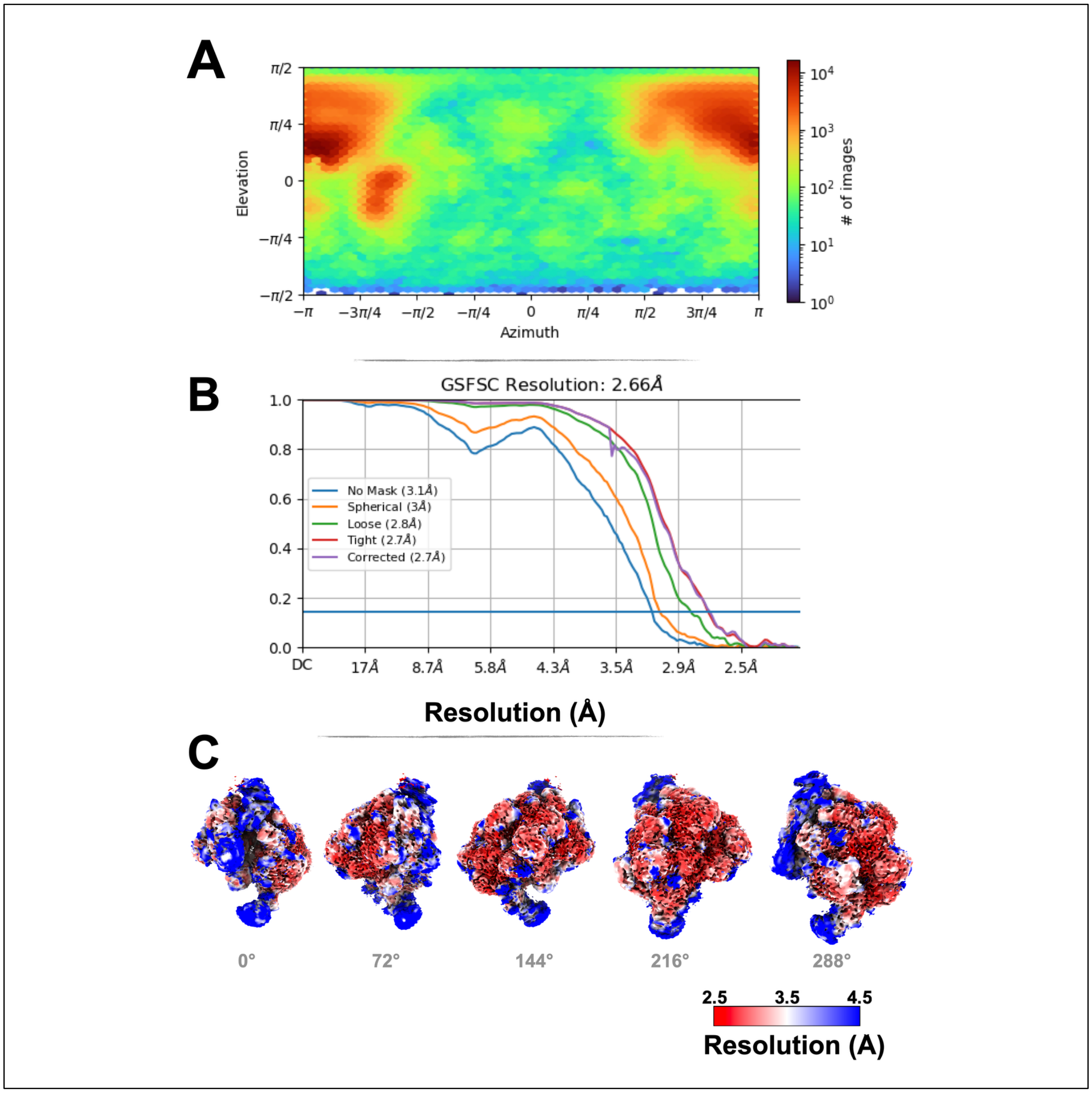

**Supplementary Figure 3. EM Statistics, Fourier Shell Correlation, and Local Resolution for the P45-J50 Data Set ($B_6P_5$, 2.66 Å).** A) Visual representation from CryoSPARC of the number of particles against the elevation and azimuthal angles of the $B_6P_5$ particles that produced the 2.66 Å reconstruction described in the text. The plot is colored from $10^0$ (blue) to $10^4$ (red) particles. We noted some evidence of a preferred orientation (red clusters, at + or - π), however, this did not affect our ability to interpret the map. B) Five Fourier shell correlation (FSC) plots as implemented in CryoSPARC. These plots were calculated by applying no mask, a spherical mask, a loose mask, a tight mask, and a tight mask with correction around the $B_6P_5$ complex (27). The resolution of our map is 2.66 Å at the FSC value of 0.143 for the tight mask with correction plot. C) Local resolution of the 2.66 Å $B_6P_5$ EM map plotted from 2.5 Å (red) to 4.5 Å (blue). The images are related by 72° (0°, 72°, 144°, 216°, 288°). The NTD of DnaB chain A, domain I of λP (chain Z), and all the λP domains II, colored in blue, represent the lower resolution portions of our map. The rest of the map is colored red and captures the better-resolved segments.

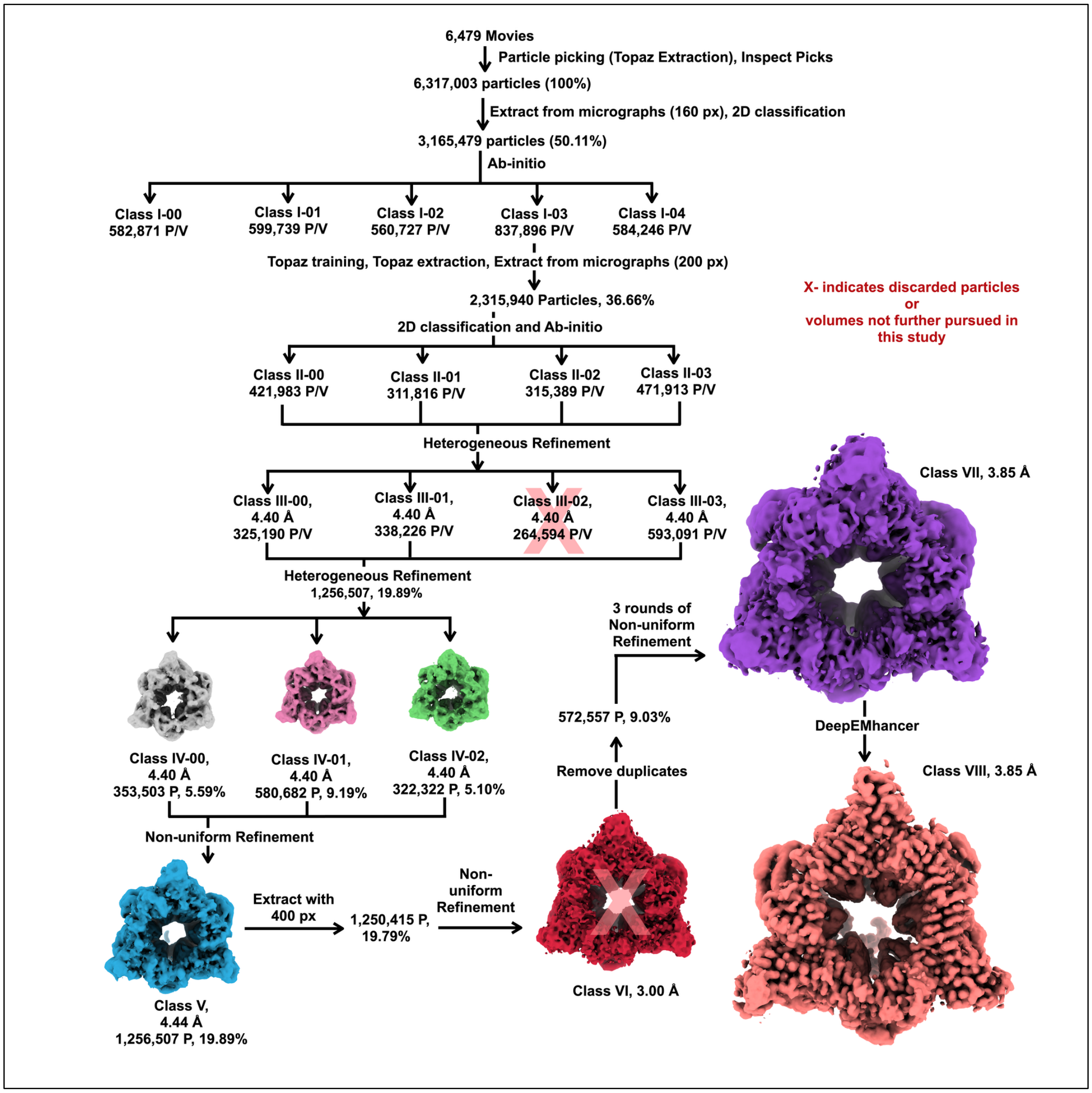

**Supplementary Figure 4. Processing of the P155-J148 Cryo-EM Data Set ($B_6P_6$, 3.85 Å).** The calculations are detailed in the Methods section. The volumes are distinguished by color. The 3.85 Å volume (purple) represents the highest-resolution EM density map we obtained for the $B_6P_6$ complex. Additional map improvement was achieved by processing with DeepEMhancer ((30), salmon-colored volume). The percentages are calculated based on the total number of particles picked.

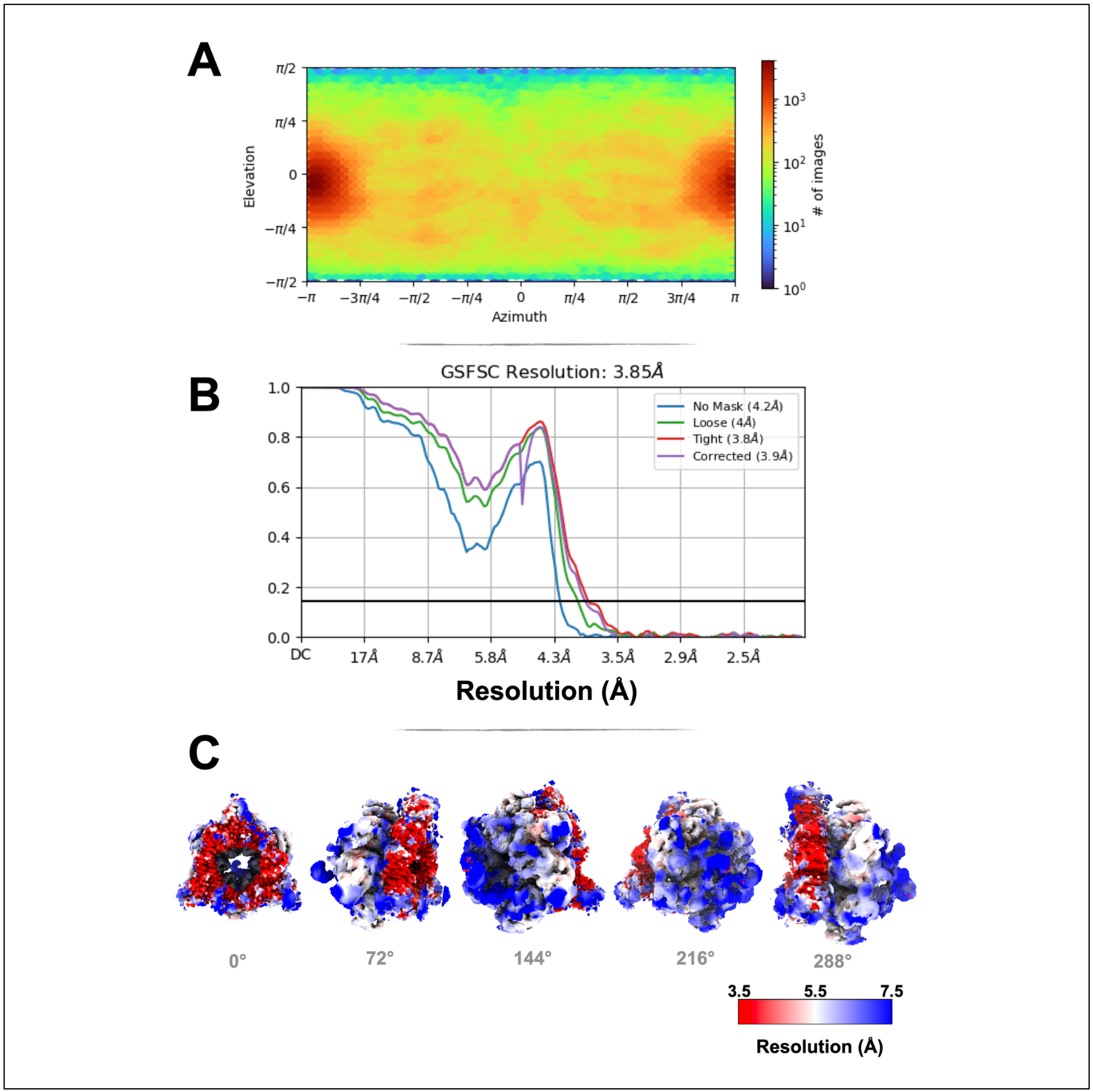

**Supplementary Figure 5. EM Statistics, Fourier Shell Correlation, and Local Resolution for the P155-J148 Data Set ($B_6P_6$, 3.85 Å).** A) Visual representation from CryoSPARC of the number of particles against the elevation and azimuthal angles of the $B_6P_6$ particles that produced the 3.85 Å reconstruction described in the text. The plot is colored from $10^0$ (blue) to $10^4$ (red) particles. Some preferred orientation in the particles (red clusters, at + π or - π) was evident, however, this did not affect our ability to interpret the map. B) Five Fourier shell correlation (FSC)

plots as implemented in CryoSPARC. These plots were calculated by applying no mask, a spherical mask, a loose mask, a tight mask, and a tight mask with correction around the $B_6P_6$ complex (27). The resolution of our map is 3.85 Å at the FSC value of 0.143 for the tight mask with correction plot. C) Local resolution of the 3.85 Å $B_6P_6$ EM map plotted from 3.5 Å (red) to 7.5 Å (blue). The images are related by 72° (0°, 72°, 144°, 216°, 288°).

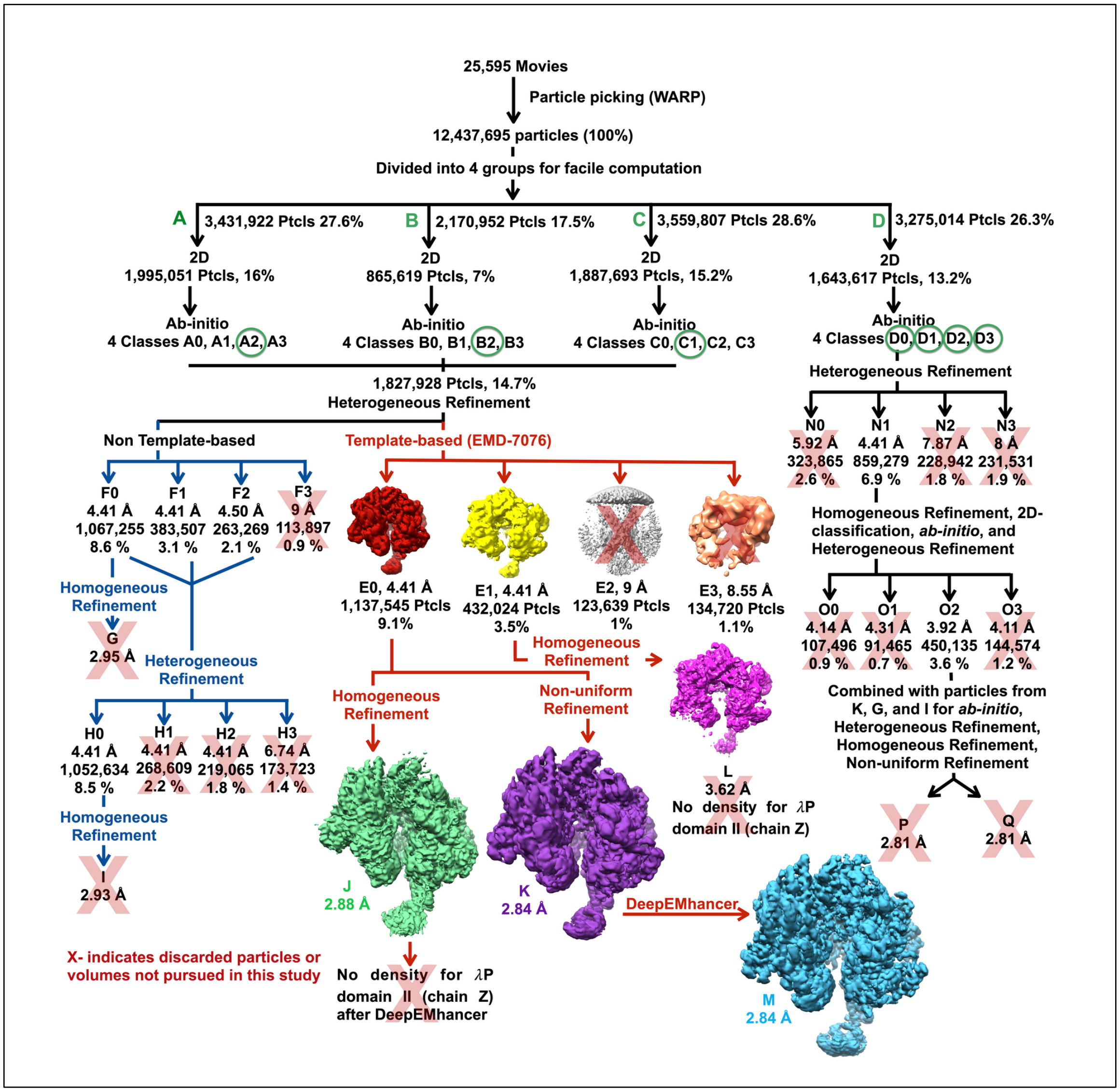

**Supplementary Figure 6. Processing of the P11-J107 Cryo-EM Data Set ($B_6P_5$, 2.84 Å).** The calculations are described in the Supplementary Methods section. The volumes are colored to distinguish them. The 2.84 Å volume (purple) represents this data set's highest-resolution EM density map. Additional map improvement was achieved by processing with DeepEMhancer ((30), cyan volume). The percentages are calculated based on the total number of particles picked.

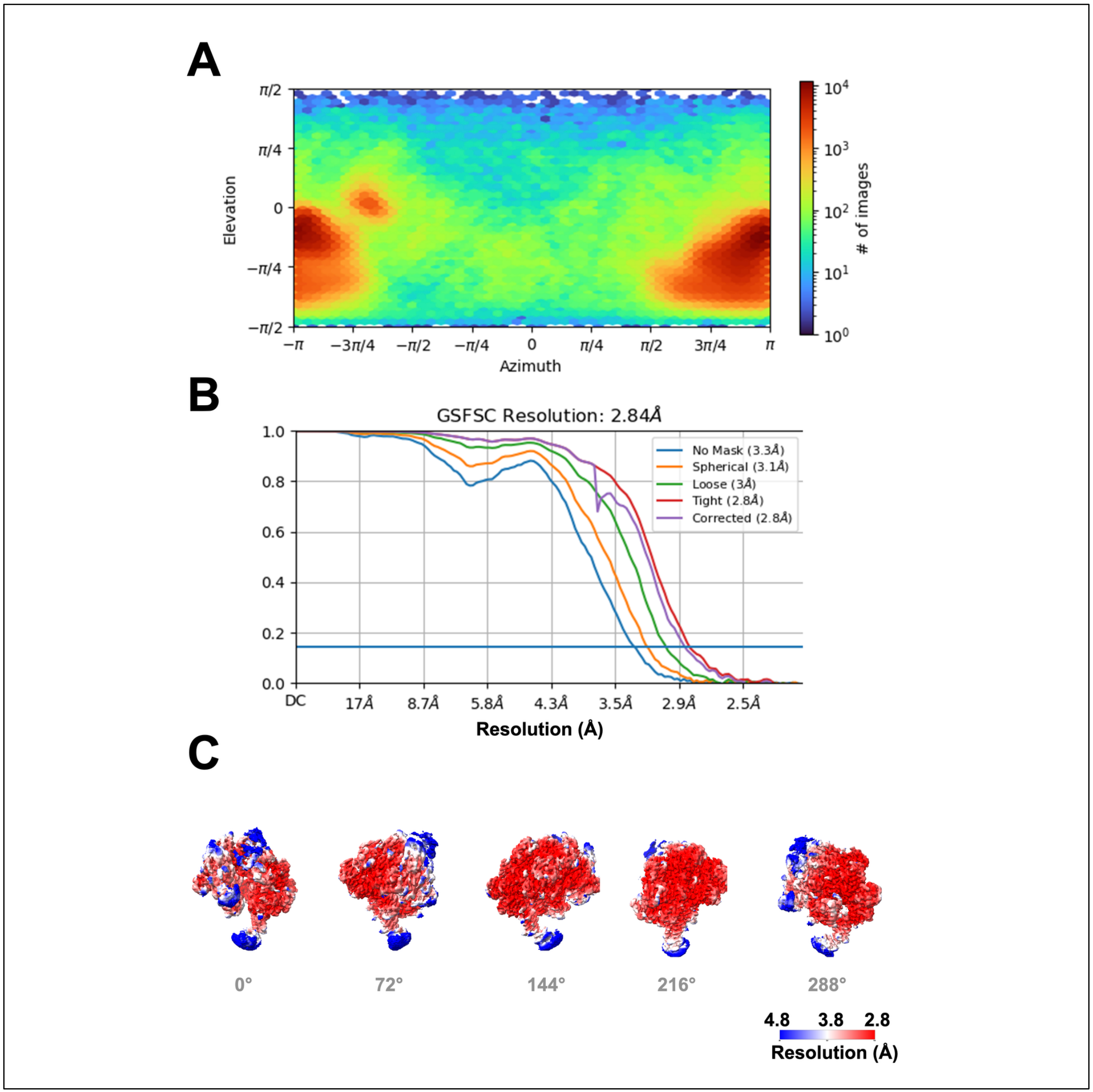

**Supplementary Figure 7. EM Statistics, Fourier Shell Correlation, and Local Resolution for the P11-J107 Data Set ($B_6P_5$, 2.84 Å).** A) Visual representation from CryoSPARC of the elevation and azimuthal angles of the BP particles that led to the 2.84 Å reconstruction described in the text. The plot is colored from $10^0$ (blue) to $10^4$ (red) particles. Some preferred orientation in the particles (red clusters, at + π or - π) was evident, however, this did not affect our ability to interpret the map. B) Five Fourier shell correlation (FSC) plots as implemented in

CryoSPARC. These plots were calculated by applying no mask, a spherical mask, a loose mask, a tight mask, and a tight mask with correction around the BP complex (27). The resolution of our map is 2.84 Å at the FSC value of 0.143 for the tight mask with correction plot. C) Local resolution of the 2.84 Å BP EM map plotted from 4.8 Å (blue) to 2.8 Å (red). The images are related by 72° (0°, 72°, 144°, 216°, 288°).

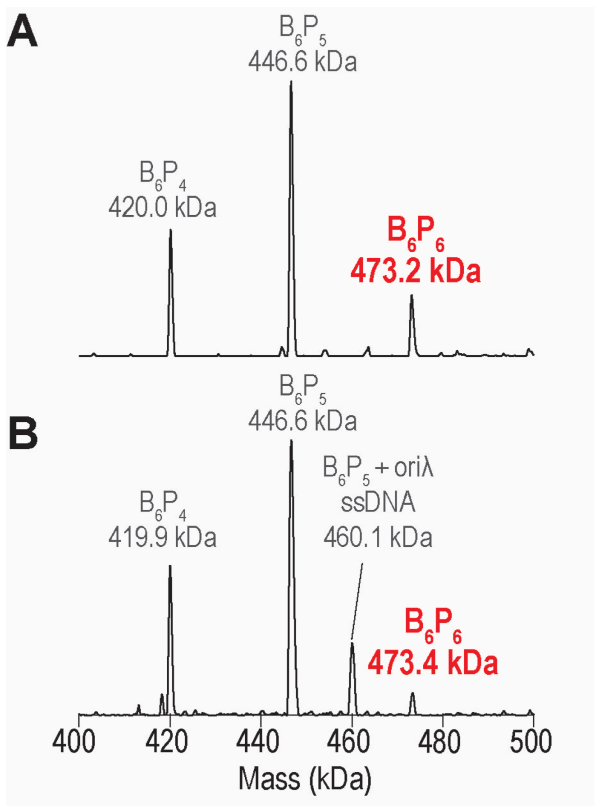

**Supplementary Figure 8. Native Mass Spectrometric (nMS) analysis of DnaB-λP samples shows a subpopulation of $B_6P_6$ assemblies.** (A) Deconvolved native MS spectrum of a representative BP sample with the predominant peak corresponding to the $B_6P_5$ assembly. A subpopulation of $B_6P_6$ complex was also observed (labeled in red) (8). (B) Deconvolved native MS spectrum for a sample containing BP, 43-mer Oriλ ssDNA and λO CTD. A $B_6P_5$-ssDNA complex was observed, but no assemblies containing BP, ssDNA and λO CTD protein were detected. Again, a subpopulation of $B_6P_6$ complex was found. The two samples are in 500 mM ammonium acetate, 0.5 mM magnesium acetate, 0.01% Tween-20 pH 7.5.

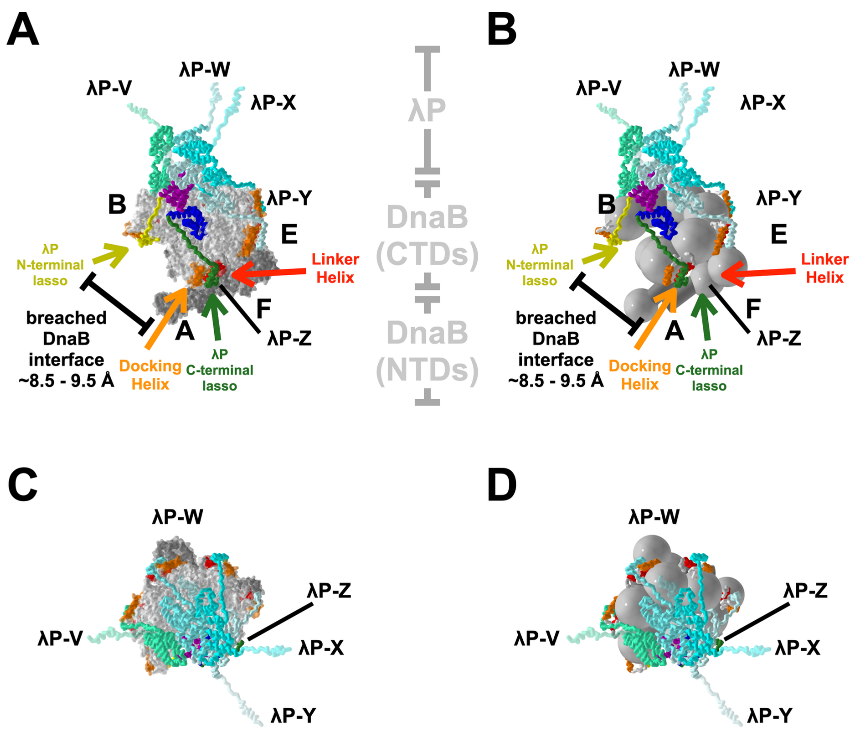

**E**

| $B_6P_5$ | DH (°) | LH (°) |
|---|---|---|
| Z | 7.9 | 5.3 |
| Y | 8.1 | 3.3 |
| X | 8.4 | 4.8 |
| W | 7.9 | 2.3 |
| V | 7.7 | 3.0 |
| Avg | 8.0 | 3.7 |

**Supplementary Figure 9. Additional Poses of the $B_6P_5$ Complex.** The $B_6P_5$ complex is represented in the same manner as in Figure 2. Panels A and B are rotated by 30° along the vertical Z axis from the pose in Figure 2. The pose in panels C and D is rotated by 90° along the horizontal X axis from the pose in Figure 2. Panel E lists the rotation angles experienced by the DH and LH elements in the transition from the closed planar to the open spiral forms.

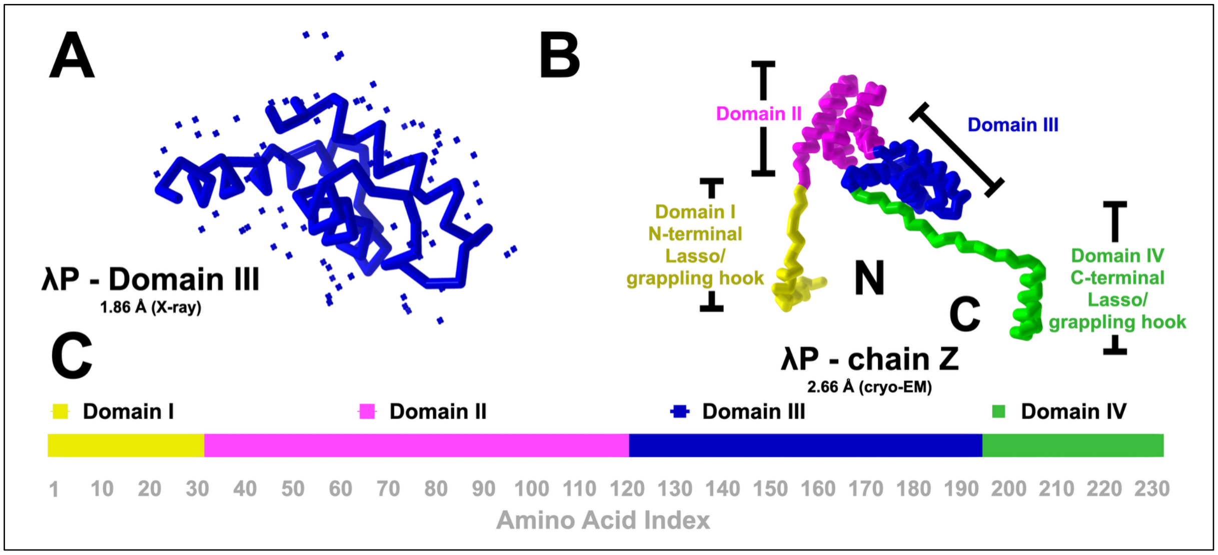

**Supplementary Figure 10. The Complete Structure of the λP Helicase Loader.** A) PyMol ribbon representation of the 1.86 Å crystal structure of λP domain III. Water molecules in the structure are depicted as plus (+) signs. B) Model of the complete structure of the complete structure of λP (chain Z) from the 2.66 Å $B_6P_5$ complex. C) Linear domain map of λP. The models in panels A and B are colored as shown in panel C.

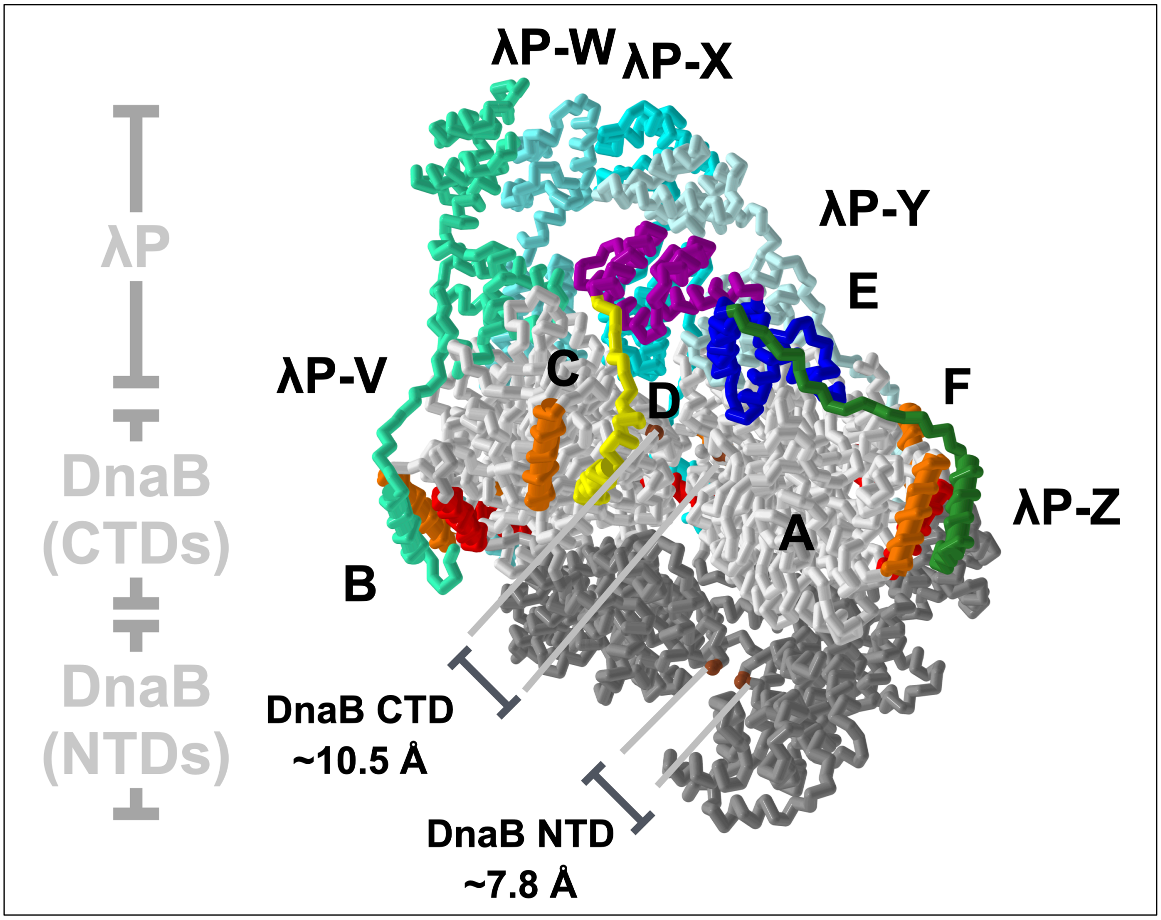

**Supplementary Figure 11. Breach in the NTD and CTD layers of the Open Spiral $B_6P_5$ Complex.** The pose and representation of the $B_6P_5$ complex are identical to those in Figure 2. Positions on the NTD and CTD, colored in dark red, represent points of closest approach in the breach and, therefore, the length where the breach was measured. The DnaB (chains A: F) NTD tier is colored in dark gray, and the CTD layer is in light gray, with the DH and LH elements colored and labeled in orange and red, respectively. The five λP loader molecules (labeled V, W, X, Y, and Z) are depicted in the ribbon representation and colored in shades of blue, save for chain Z, which is colored by domain (domain I: yellow, domain II: purple; domain III: blue, and domain IV: green).

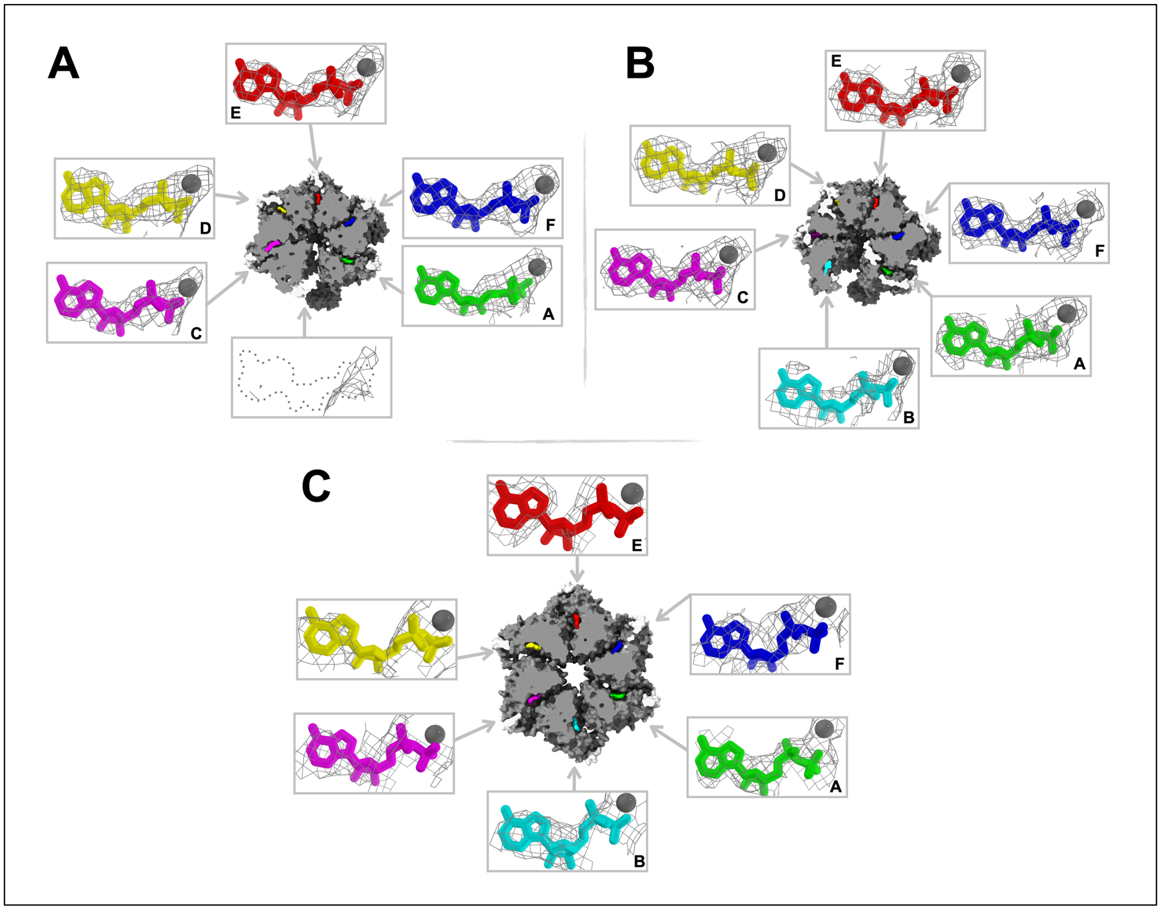

**Supplementary Figure 12. Nucleotide Occupancy of the $B_6P_5$ and $B_6P_6$ complexes.** EM density corresponding to the nucleotide-binding sites in DnaB within the $B_6P_5$ (A: $B_6P_5$; P11-J107, 2.84 Å; B: $B_6P_5$; P45-J50, 2.66 Å;) and $B_6P_6$ (C: $B_6P_6$; P155-J148, 3.85 Å) complexes. Density maps (produced with the PyMol "carve=2.5" command) are superimposed on the modeled ADP and Mg ions. DnaB is shown looking down the CTD tier in the clipped surface representation to illustrate the positions of each nucleotide binding site. A) In the 2.84 Å $B_6P_5$ structure (P11-J107), five sites show ADP density contoured at 11; the sixth site on chain B exhibits no density even at a lower contour value of 7. The outlined shape marks the position of the missing ADP. B) All six sites show ADP density in the 2.66 Å $B_6P_5$ structure (P45-J50). Except for chain B, which is contoured at

7, all sites are contoured at 11; this implies chain B has a lower occupancy. C) All six sites show density for ADP in the 3.85 Å $B_6P_6$ structure (P155-J148). The density for all $B_6P_6$ sites is contoured at 5 to enable comparison. The density quality at the ADP sites on chains B, C, and D is lower than the other sites, indicating a lower occupancy.

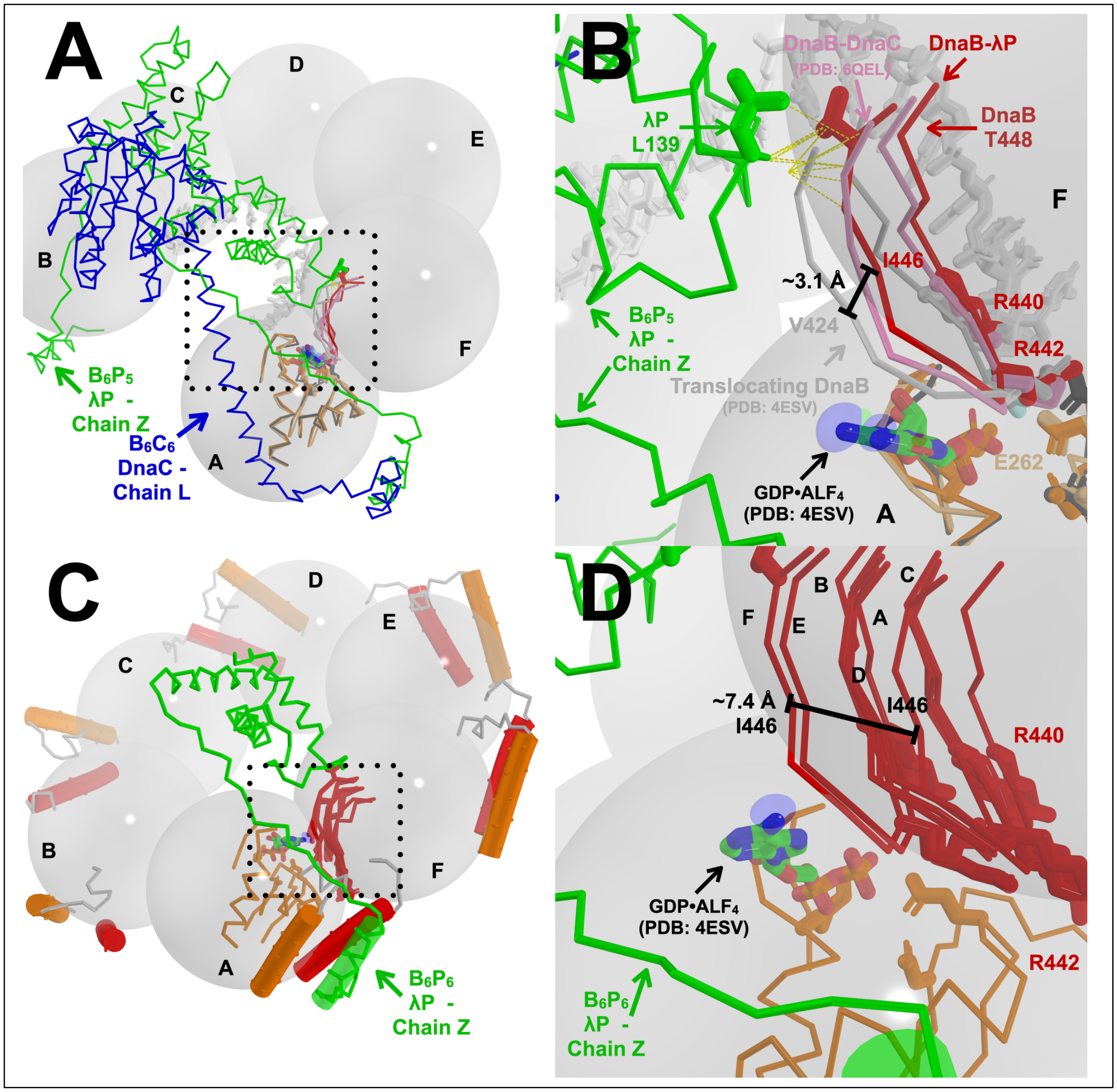

**Supplementary Figure 13. Misalignment of the ATPase centers in the DnaB Open Spiral.** A) Transition to the open spiral form of DnaB leads to a misalignment of the ATPase center at the interface between two CTDs. Schematic view of the $B_6P_5$ complex wherein the CTDs in the open spiral are represented by transparent and labeled grey spheres. Chain Z of the λP ensemble (green) and Chain L from *E. coli* DnaB•DnaC structure (blue, (PDB: 6QEL (9))) are shown in the PyMol ribbon representation. The *E. coli* DnaB•DnaC structure is aligned onto the CTD of chain A of the $B_6P_5$ complex. Portions of the $B_6P_5$ CTDs from chain A (light orange) and chain F (salmon) also appear as ribbons. B) A closeup (and rotated) view of the area within the dotted-line square in panel A. Colored in red is a ribbon representation of the arginine finger β-hairpin of chain F of the current $B_6P_5$ structure. The corresponding elements from the *E. coli* DnaB-DnaC and *B.st* DnaB-ssDNA (PDB: 4ESV, (4)) structures are colored in pink and grey. In $B_6P_5$, the arginine finger β-hairpin is 3.1 Å away from the position in 4ESV. The ssDNA from the translocating *B.st* DnaB-ssDNA appears in grey in the background. Several residues in λP and DnaB in the $B_6P_5$ complex are shown in stick format. C) The $B_6P_6$ complex is diagrammed and posed like in panel A. D) A closeup (and rotated) view of the area within the dotted-line square in panel C. In $B_6P_6$, the position of the chain F arginine finger β-hairpin is ~2 Å from the position in 4ESV. However, the six arginine finger β-hairpins in $B_6P_6$ are each sub-optimally positioned for catalysis. Their positions span a distance of 7.4 Å.

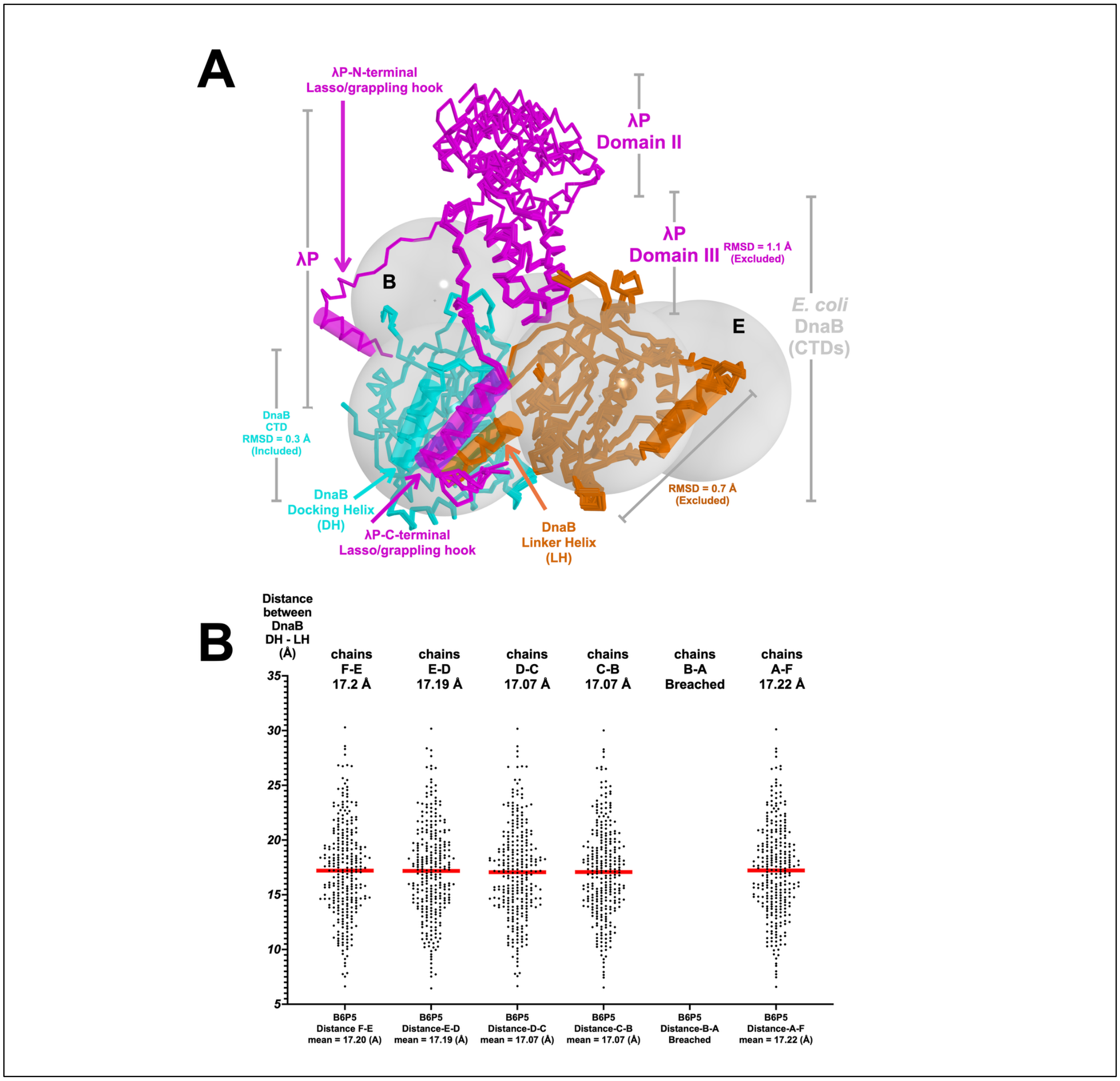

**Supplementary Figure 14. Superposition of the Five DnaB – λP Interfaces in the $B_6P_5$ complex.** The five $B_2P_1$ sub-structures in the $B_6P_5$ complex display a rigidity indicated by the structural alignment (panel A) and the distribution of pairwise distances (panel B) between each interface's DH and LH elements. A) Each instance of the $B_2P_1$ sub-structure was superimposed on the 'left' DnaB subunit, as in the frame of reference provided by this pose. The superposition calculation included residues 206 to 289 and 307 to 471; residues corresponding to the LH element (181-196) and the DH element (290-306) were excluded. The root-mean-square deviations of the superpositions average 0.3 Å on ~200 Cα atoms (labeled 'included'). The RMSD of structural domains excluded from the superposition are labeled 'excluded.' The 'left' DnaB subunit is colored in cyan, and the 'right' subunit is colored in orange. Each of the five λP chains is colored in magenta. DnaB CTDs are represented as spheres drawn on their respective centers of gravity. For reference, two of the DnaB CTDs are labeled by chain. B) Distribution of pairwise distances between the DH (residues 291:307) and LH (residues 182:198) elements in the $B_6P_5$ complex calculated across the five instances of the $B_2P_1$ sub-structure in the complex. This analysis excludes distances between chains B and A that line the breach.

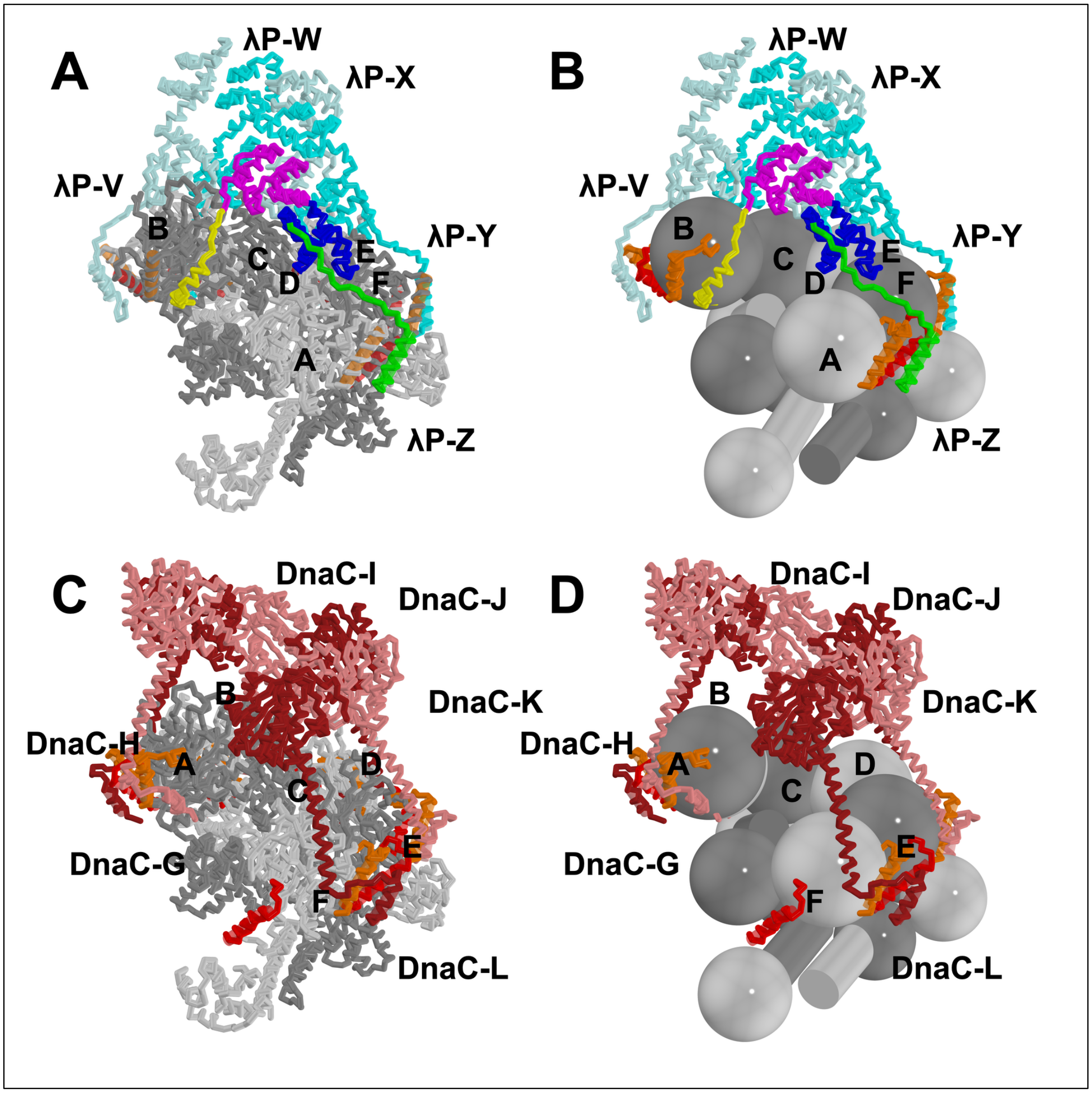

**Supplementary Figure 15. Comparison of Loader and Helicase Interfaces in the $B_6P_5$ and $B_6C_6$ complexes.** The λP loader ensemble (A, B) forms a more extensive interface with DnaB than DnaC (C, D). In panels A and B, the pose and representation of the $B_6P_5$ complex are identical to that in Figure 2, except that the DnaB (chains A: F) are alternately colored in dark gray and light gray. The DH and LH elements are colored and labeled orange and red. The five loader molecules (labeled V, W, X, Y, and Z) are depicted in the ribbon representation and colored in shades of blue, save for chain Z which is colored by domain (domain I: yellow, domain II: purple; domain III: blue, and domain IV: green). The $B_6C_6$ complex is illustrated in panels C and D. DnaB in the $B_6C_6$ complex is depicted as in panels A and B. The six DnaC monomers are alternately colored in red and pink.

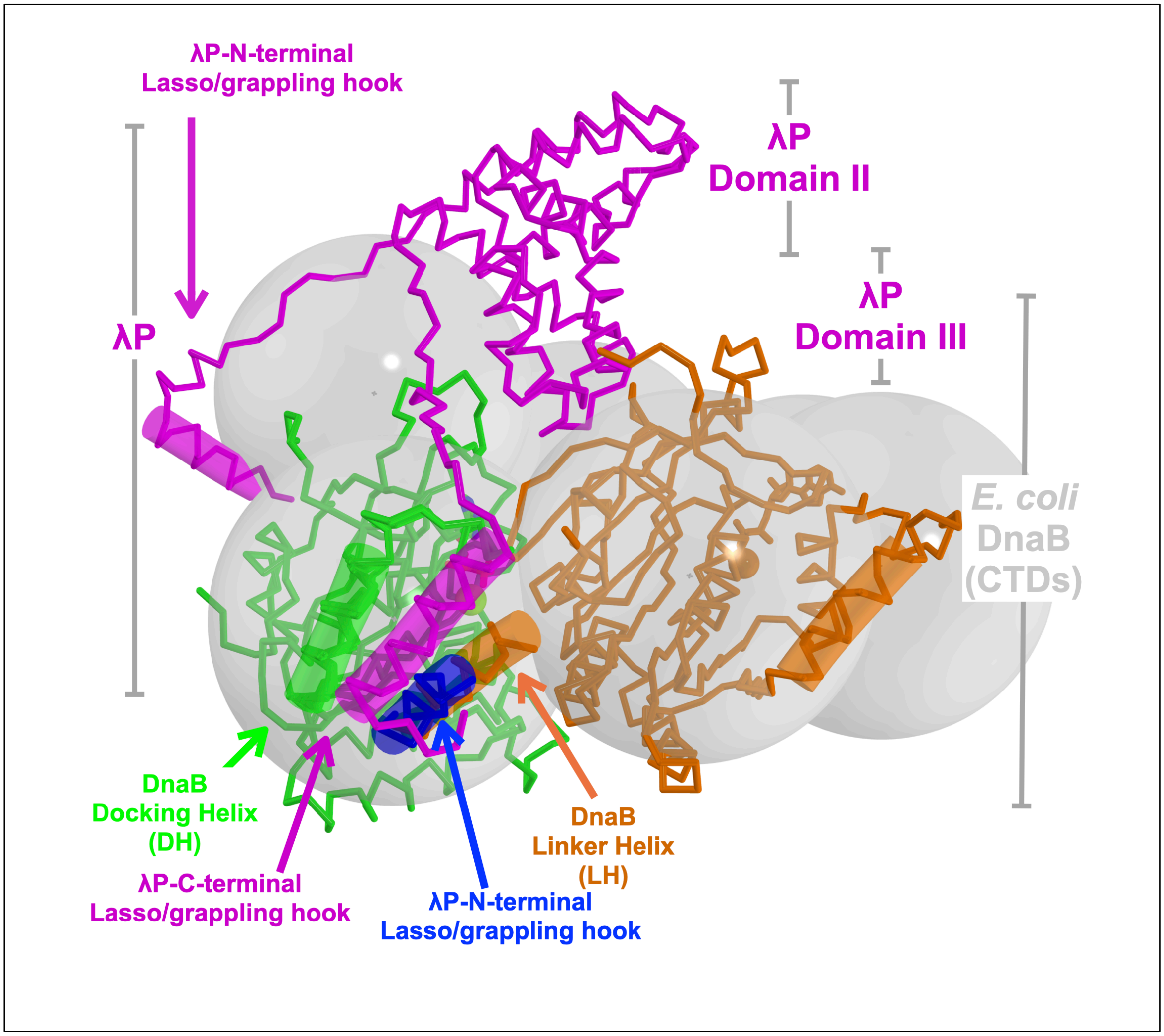

**Supplementary Figure 16. The N-terminal Lasso λP (chain Z) of $B_6P_5$ Complex Adopts a Unique Configuration Relative to the DnaB CTD.** The disposition of the chain Z N-terminal lasso relative to the chain B DnaB CTD overlaps with the lasso/grappling hook of DnaC (PDB: 6QEL, chain G, (9) (Figures 2 and 4). The DnaB subunit - chain B that contacts the N-terminal Lasso of λP (chain Z) was superimposed on DnaB - chain A. The superposition calculation included residues 206 to 289 and 307 to 471; residues corresponding to the LH element (181-196) and the DH element (290-306) were excluded. The RMSD of the superpositions was 0.46 Å over 224 Cα atoms. The 'left' DnaB subunit is colored in green, and the 'right' subunit is colored in orange. The chain B DnaB CTD is colored in cyan. DnaB CTDs are represented as spheres drawn on their respective centers of gravity.

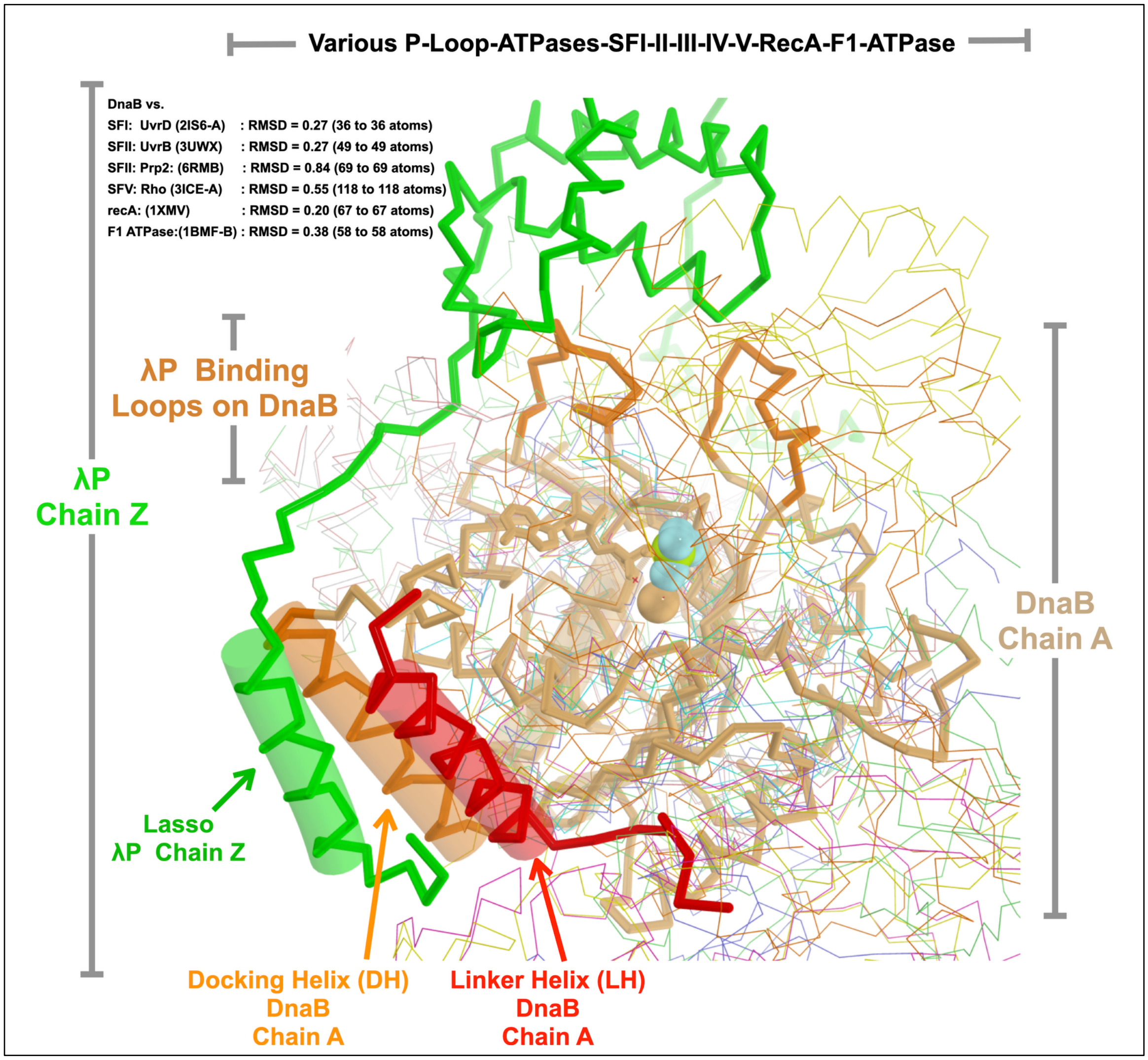

**Supplementary Figure 17. The λP binding site on DnaB consists of insertions to the RecA fold.** The λP ensemble contacts the DnaB hexamer via five widely dispersed segments on the DnaB protomer. Residues 290:307 (DH element), residues 386:396, and residues 421:431 are depicted in a darker orange color and are found on one CTD, while residues 182:199 (the LH element), colored in red, are found on the adjacent CTD. Each of these positions on DnaB is an insertion into the conserved RecA fold and provides points of contact for sites #1, #2, #3, and #4 between helicase and loader (site #5 is on the NTD, which is not part of the RecA fold). Chain A and LH elements of chain B from DnaB in the BP complex are depicted in the PyMol ribbon representation and colored in light orange and red, respectively. Superimposed on chain A of the DnaB are other members of the RecA family, including UvrD (helicase superfamily I, PDB: 2IS6, RMSD: 0.27 on 36 atoms (36)), UvrB (helicase superfamily II, PDB: 3UWX, RMSD: 0.27 on 49 atoms (37)), Prp2 (helicase superfamily II, PDB:6RMB, RMSD: 0.84 on 69 atoms (38)), Rho helicase (superfamily V, PDB: 3ICE, RMSD: 0.55 on 118 (39)), RecA (PDB: 1XMV, RMSD: 0.20 on 67 atoms (40)), and the F1-ATPase (PDB: 1BMF, RMSD: 0.38 on 58 atoms, (41)). Each of the above structures is shown in the PyMol ribbon representation, albeit in a smaller width than used for DnaB. Notably, the dark-orange-colored elements are found in none of the above RecA family structures.

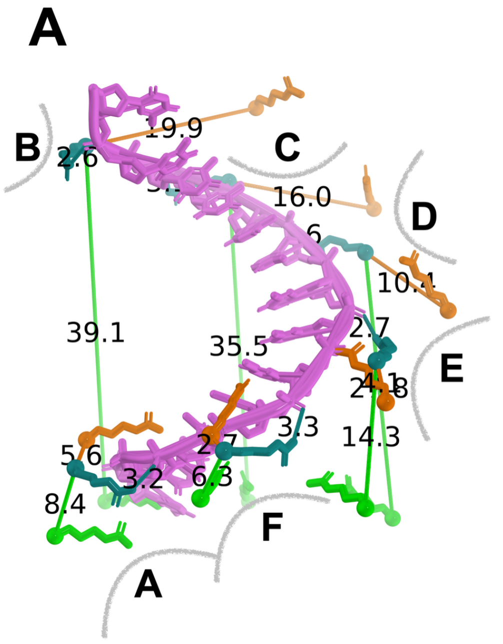

B

| | R403 ($B_6P_5$) | R403 ($B_6P_6$) |
|---|---|---|
| R381-A (PDB: 4ESV) | 5.6 | 8.4 |
| R381-F (PDB: 4ESV) | 2.7 | 6.3 |
| R381-E (PDB: 4ESV) | 4.1 | 14.3 |
| R381-D (PDB: 4ESV) | 10.4 | 27.8 |
| R381-C (PDB: 4ESV) | 16 | 35.5 |
| R381-B (PDB: 4ESV) | 19.9 | 39.1 |

Distances are in Å

**Supplementary Figure 18. DnaB's internal chamber ssDNA binding site is distorted in the $B_6P_5$ and $B_6P_6$ complexes.** Distortion was assessed by measuring the displacement of ssDNA binding residues in the 4ESV complex against equivalent residues in the $B_6P_5$ and $B_6P_6$ complexes. The results for one residue, 4ESV R381(*B.st*), and its equivalent in *E. coli* DnaB (R403) are shown graphically (A) and in tabular form (B). A) ssDNA from the B.st 4ESV complex is colored in light purple with R381 residues (deep teal) from chains A, F, E, D, C, and B in contact with the phosphate backbone. The corresponding residue (R403) from the $B_6P_5$ and $B_6P_6$ complexes is shown in orange and green, along with its displacement from a position optimal for contacts to ssDNA.

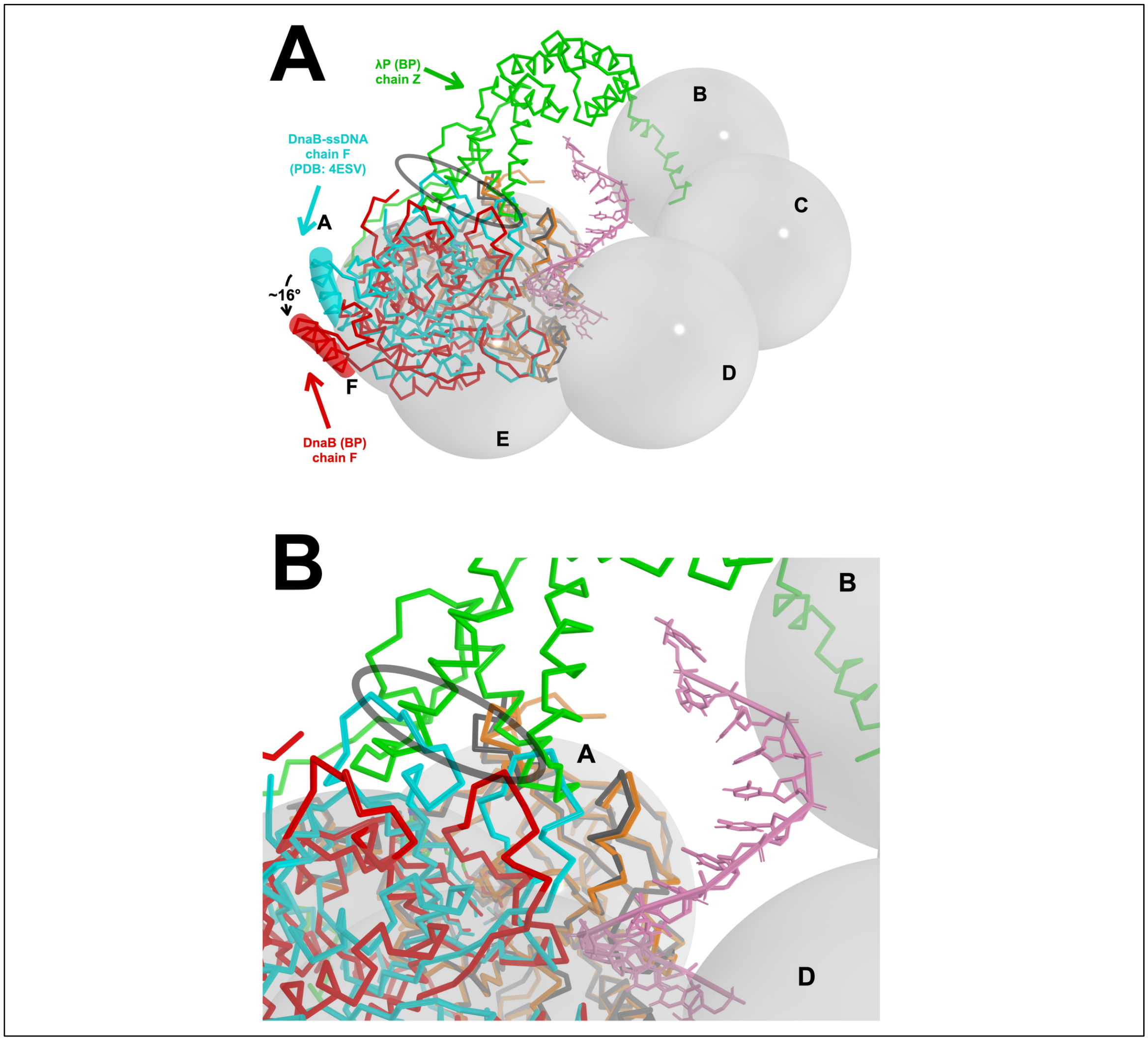

**Supplementary Figure 19. The $B_6P_5$ Complex Cannot Adopt the ssDNA Configuration of the Translocating Form.** A) Disposition of domain III of the λP loader in the $B_6P_5$ complex precludes adoption by DnaB of the ssDNA binding mode seen in the 4ESV translocation state. DnaB chain A (grey) of the 4ESV structure ssDNA ((4)) is superimposed on DnaB chain A (orange) of the $B_6P_5$ complex using residues 206:471 but omitting the DH (residues 290:306) and LH (181:196) elements from the calculation. In the foreground, chains F of DnaB from the $B_6P_5$ (red) and 4ESV (cyan) complexes are shown; these chains were also omitted from the superposition. The ssDNA from the DNA complex (pink) is depicted using the PyMol sticks representation. The black oval highlights the molecular clash between the λP loader and a CTD from the ssDNA complex. The open DnaB spiral within the $B_6P_5$ complex is represented by grey spheres labeled by chain. Chain Z of the λP ensemble (green) is shown in the PyMol ribbon representation. B) Closeup of panel A.

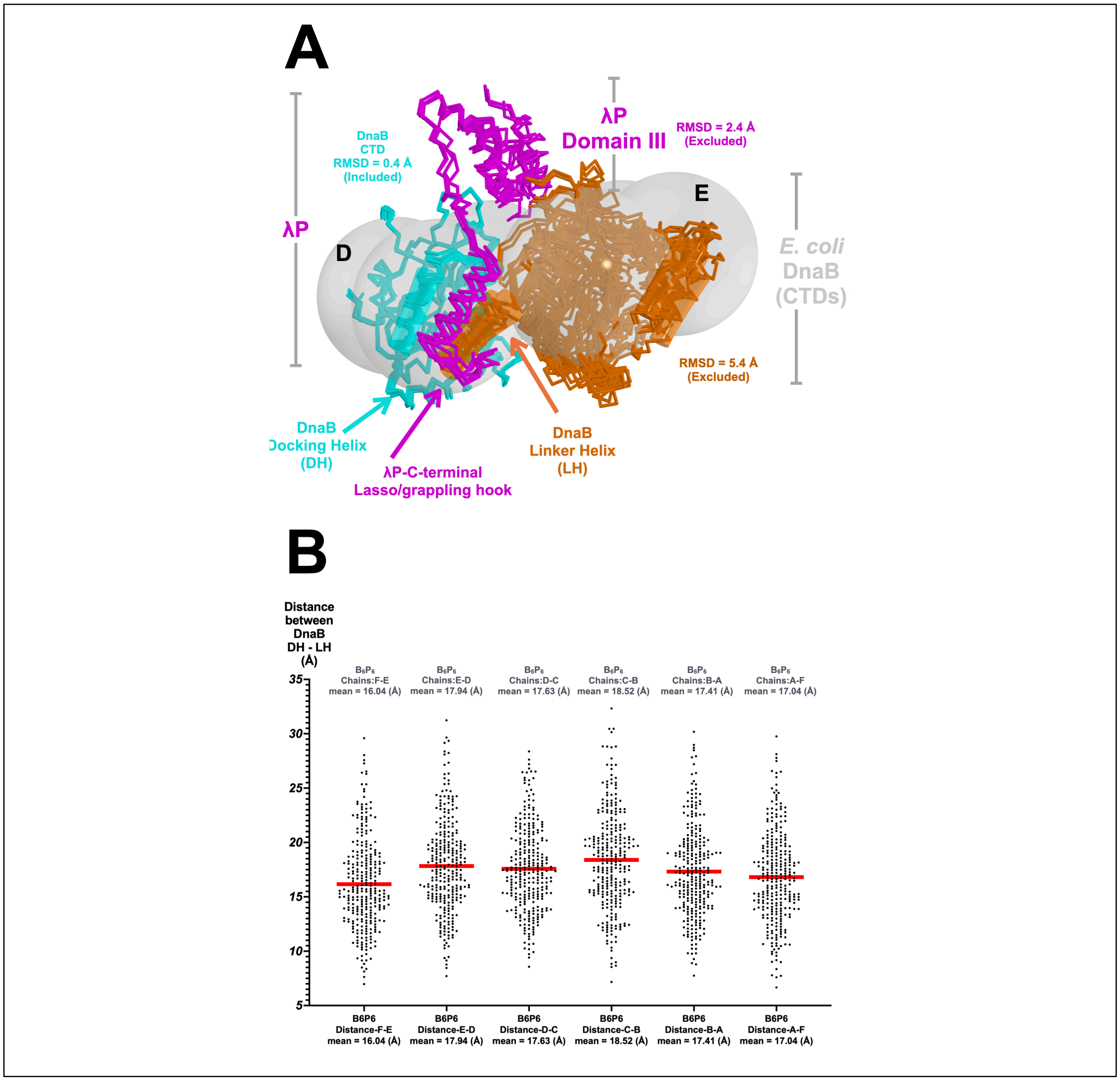

**Supplementary Figure 20. The $B_6P_6$ complex is inchoate and has not reached its final form.** This is indicated by the divergence in RMSD of superimposed $B_2P_1$ instances and the variability in the distances between the DH – LH elements. A) Each instance of the $B_2P_1$ sub-structure from the $B_6P_6$ complex was superimposed on the 'left' DnaB subunit in the frame of reference of this pose. The superposition calculation included residues 206 to 289 and 307 to 471; residues corresponding to the LH element (181:196) and the DH element (290:306) were excluded. The root-mean-square deviations of the superposition averaged 0.4 Å on ~200 Cα atoms (labeled ‘included’). The RMSD of structural domains excluded from the superposition are labeled ‘excluded.’ The ‘left’ DnaB subunit is colored in cyan, and the ‘right’ subunit is colored in orange. For reference, two of the DnaB CTDs are labeled by chain. Each of the five λP chains is colored in magenta. DnaB CTDs are represented as spheres drawn on their respective centers of gravity. B) Distribution of pairwise distances between the DH (residues 291:307) and LH (residues 182:198) elements in the $B_6P_6$ complex calculated across the six instances of the $B_2P_1$ sub-structure. .

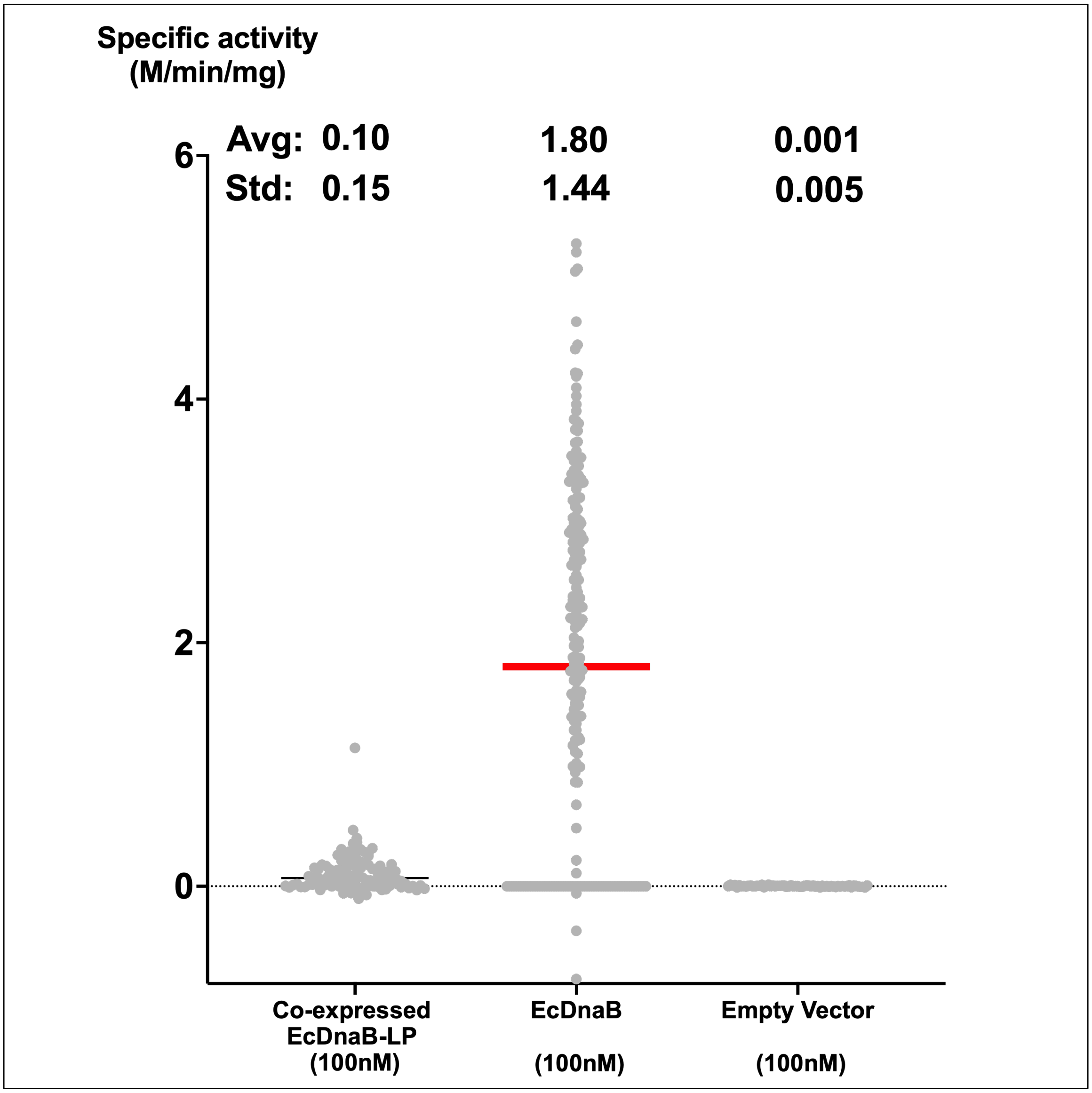

**Supplementary Figure 21. λP Suppresses the ATPase Activity of DnaB.** Full-length λP completely suppresses the ATPase activity of DnaB. The average and standard deviation of each measurement appear below each plot. The red line represents the mean value. The spread in the measurements in the EcDnaB sample captures variability in our preparations. The empty vector sample represents a bacterial protein extract prepared using the exact set of steps described above for DnaB, except that the pET24 vector that was used did not include either the DnaB or λP genes.

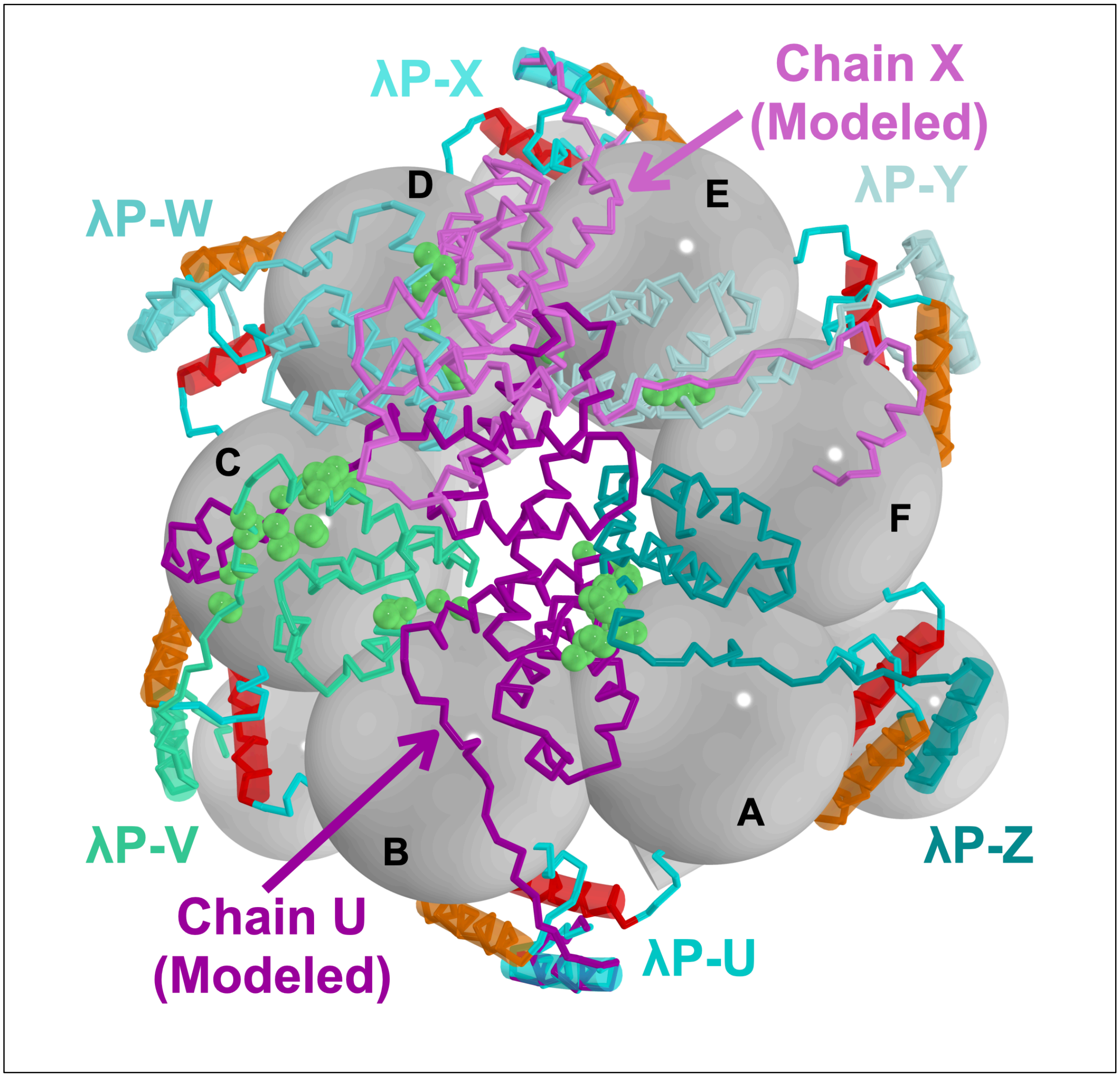

**Supplementary Figure 22. Consequences of Modeling the Two Disordered Chains of λP onto the $B_6P_6$ complex.** The complete chains U (dark purple) and X (light purple) of λP are incompatible with the positions of chains V, W, Y, and Z in the $B_6P_6$ complex. Incompatibility of the disordered domain IIIs of chain U and X with the $B_6P_6$ complex is indicated by the clash (close approach; within 1.5 Å) with chains W and Y, and V and Z. The light-green spheres show atomic positions in the above chains within 1.5 Å of chains U and X.

**Supplementary Movie 1. Transition of DnaB from the Closed Planar to the Ajar Planar Configuration.** DnaB is depicted in the ribbons format (A) and the sphere/cylinder design language (B). DnaB chains are colored in alternating shades of gray. DnaB's DH and LH helices are shown in the ribbon format and colored orange and red. The six chains of the λP loader are drawn as ribbons and colored in alternating cyan and teal. The animation starts from the closed planar DnaB configuration (PDB: 4NMN, (2)) and morphs into the planar ajar configuration (this work), with the appearance of the six λP chains. The morph depicts the transition and then animates back to the closed planar form. The animation repeats at two DnaB poses, the first looking down the NTD tier and the second looking down the CTD tier.

**Supplementary Movie 2. Transition of DnaB from the Ajar Planar to the Open Spiral Configuration.** DnaB is depicted in the ribbons format (A) and the sphere/cylinder design language (B). DnaB chains are colored in alternating shades of gray. DnaB's DH and LH helices are shown in the ribbon format and colored orange and red. The six chains of the λP loader are drawn as ribbons and colored in alternating cyan and teal. The animation starts from the ajar planar DnaB configuration (this work) and morphs into the open spiral configuration (this work). The transition is accompanied by the appearance of the five λP chains. For clarity, the six λP chains in the ajar planar form are not shown. The morph depicts the transition and then animates back to the ajar planar form. The animation repeats at two DnaB poses, the first looking down the NTD tier and the second looking down the CTD tier.

#### Supplementary Table 1. Expression Constructs of the λP Helicase Loader

| Expression Plasmid (Name, Insert, Tag) | Primers (F: forward primer, R: reverse primer) |
|---|---|
| pDB005-pRSFDuet-LambdaP-105-205-6X-N-His | PCR/Ligation<br>F:CATGCCATGGGTCATCACCATCACCATCACCAGTTTGTTGCATGGTGCCGG<br>R:CGCGGATCCTTACATGACAGGAAGTTGTTTTACTGGTTCAGGGATCGCCTCACCACGGTTAATTCT |
| pDB006-pRSFDuet-LambdaP-105-210-6X-N-His | PCR/Ligation<br>F:CATGCCATGGGTCATCACCATCACCATCACCAGTTTGTTGCATGGTGCCGG<br>R:CGCGGATCCTTATAGAGGTCTACCGCCCATGACAGGAAGTTGTTTTACTGGTTCAGGGATCG |
| pDB007-pRSFDuet-LambdaP-105-215-6X-N-His | PCR/Ligation<br>F:CATGCCATGGGTCATCACCATCACCATCACCAGTTTGTTGCATGGTGCCGG<br>R:CGCGGATCCTTAAGCCTGTGCACGATTTAGAGGTCTACCGCCCATGACAGG |
| pDB008-pRSFDuet-LambdaP-105-220-6X-N-His | PCR/Ligation<br>F:CATGCCATGGGTCATCACCATCACCATCACCAGTTTGTTGCATGGTGCCGG<br>R:CGCGGATCCTTATGCGATCTTCGCCAGAGCCTGTGCACGATTTAGAGG |
| pDB010-pRSFDuet-LambdaP-110-210-6X-N-His | PCR/Ligation<br>F:CATGCCATGGGTCATCACCATCACCATCACTGCCGGGAAGAAGCATCCGTTA<br>R:CGCGGATCCTTATAGAGGTCTACCGCCCATGACAGGAAGTTGTTTTACTGGTTCAGGGATCG |
| pDB011-pRSFDuet-LambdaP-110-215-6X-N-His | PCR/Ligation<br>F:CATGCCATGGGTCATCACCATCACCATCACTGCCGGGAAGAAGCATCCGTTA<br>R:CGCGGATCCTTAAGCCTGTGCACGATTTAGAGGTCTACCGCCCATGACAGG |
| pDB017-pRSFDuet-LambdaP-105-205-6X-C-His | PCR/Ligation<br>F:CATGCCATGGGTCAGTTTGTTGCATGGTGCCGGGAAGAAGCATC<br>R:CGCGGATCCTTAGTGATGGTGATGGTGATGCATGACAGGAAGTTGTTTTACTGGTTCAGG |
| pDB018-pRSFDuet-LambdaP-105-210-6X-C-His | PCR/Ligation<br>F:CATGCCATGGGTCAGTTTGTTGCATGGTGCCGGGAAGAAGCATC<br>R:CGCGGATCCTTAGTGATGGTGATGGTGATGTAGAGGTCTACCGCCCATGACAGGAAG |
| pDB019-pRSFDuet-LambdaP-105-215-6X-C-His | PCR/Ligation<br>F:CATGCCATGGGTCAGTTTGTTGCATGGTGCCGGGAAGAAGCATC<br>R:CGCGGATCCTTAGTGATGGTGATGGTGATGAGCCTGTGCACGATTTAGAGGTCTACC |

| | |
|---|---|
| pDB020-pRSFDuet-LambdaP-105-220-6X-C-His | PCR/Ligation<br>F:CATGCCATGGGTCAGTTTGTTGCATGGTGCCGGGAAGAAGC<br>ATC<br>R:CGCGGATCCTTAGTGATGGTGATGGTGATGTGCGATCTTCGC<br>CAGAG |
| pDB022-pRSFDuet-LambdaP-110-210-6X-C-His | PCR/Ligation<br>F:CATGCCATGGGTTGCCGGGAAGAAGCATCCGTTACCGCC<br>R:CGCGGATCCTTAGTGATGGTGATGGTGATGTAGAGGTCTACC<br>GCCCATGACAGGAAG |
| pDB023-pRSFDuet-LambdaP-110-215-6X-C-His | PCR/Ligation<br>F:CATGCCATGGGTTGCCGGGAAGAAGCATCCGTTACCGCC<br>R:CGCGGATCCTTAGTGATGGTGATGGTGATGAGCCTGTGCAC<br>GATTTAGAGGTCTACC |
| pDB065-pCDFDuet-EcDnaB-untagged | The *E. coli* DnaB gene was cut from pET24a-EcDnaB (NdeI and BamHi sites) and ligated into the corresponding sites of pCDFDuet. |
| pET24a-EcDnaB | (8) |
| pCDFDuet-Lambda-P | (8) |
| pCDFDuet-Lambda-P-C-His | PCR/Ligation<br>F:CATGCCATGGAAAACATCGCCGCACAGATG<br>R:CGCGGATCCTCAGTGATGGTGATGGTGATGTACACTTGCTCC<br>T |

**Supplementary Table 2: X-Ray Data Collection, Phasing, and Refinement**

| | λP-105 – 210-N-His (SAD) |
|---|---|
| PDB | 8V9S |
| SBGrid ID | TBA |
| **Data collection** | |
| Space group | $P3_121$ |
| Cell dimensions | |
| *a*, *b*, *c* (Å) | 49.731, 49.731, 70.772 |
| alpha, beta, gamma, (°) | 90, 90, 120 |
| Wavelength | 0.97918 (peak) |
| Resolution (Å) | 1.86 (CC1/2 = 0.678) |
| Highest resolution shell | 1.89 – 1.86 |
| $R_{sym}$ all data (in highest resolution shell) | 0.105 (0.617) |
| $\langle I / sigI \rangle$ all data (in highest resolution shell) | 18.99 (0.81) |
| Completeness (%) all data (in highest resolution shell) | 98.9 (79.4) |
| Average redundancy all data (in highest resolution shell) | 5.5 (2.5) |
| **Refinement** | |
| Resolution (Å) | 1.86 |
| No. reflections | 8718 |
| $R_{work}$ / $R_{free}$ | 0.225/0.243 |
| No. atoms | |
| Protein | 607 |
| Water | 76 |
| **B-factors** | |
| Protein | 26.73 |
| Water | 37.62 |
| **RMS deviations** | |
| Bond lengths (Å) | 0.011 |
| Bond angles (°) | 1.256 |
| **Validation** | |
| MolProbity score | 1.49 |
| Clash score | 9.2 |
| Rotamer outliers (%) | 0 |
| **Ramachandran plot (%)** | |
| Favored (%) | 98.61 |
| Allowed (%) | 1.39 |
| Outliers (%) | 0.00 |

By default, the resolution was determined using CC0.5 > 0.5 criteria.

**Supplementary Table 3: Cryogenic Electron Microscopy Data Measurement and Analysis and Coordinate Refinement**

| Dataset | P11-J107 $B_6P_5$ | P45-J50 $B_6P_5$ | P155-J148 $B_6P_6$ |
|---|---|---|---|
| PDB | 8V9T | 9OA1 | 9OA2 |
| EMDB | 43086 | 70269 | 70271 |
| **Data collection and processing** | | | |
| Microscope | Titan Krios | Titan Krios | Titan Krios |
| Voltage (kV) | 300 | 300 | 300 |
| Camera | Gatan K3 | Gatan K3 | Gatan K3 |
| Magnification | 81000 | 81000 | 81000 |
| Defocus range (um) | -1.0 to -2.5 | -1.0 to -2.5 | 0.6 to 2.7 |
| Exposure time (ms) | 2000 | 2000 | 2000 |
| Electron dose ($e^-/Å^2$) | 51.01 | 51.01 | 51.19 |
| Number of frames collected (no.) | 40 | 40 | 40 |
| Number of frames processed (no.) | 40 | 40 | 40 |
| Calibrated pixel size (Å) | 1.083 | 1.083 | 1.083 |
| Micrographs (no.) | 25595 | 7863 | 6479 |
| Total particle images (no.) | 12437695 | 7250518 | 2315940 |
| **Refinement** | | | |
| Particle per class (no.) | 1137545 | 1500880 | 572557 |
| Symmetry imposed | C1 | C1 | C1 |
| Map resolution (Å), 0.143 FSC | 2.84 | 2.66 | 3.85 |
| Map sharpening B-factor ($Å^2$) | 101.1 | 91.5 | 108.2 |
| Map versus model cross-correlation (CC (volume)) | 0.7364 | 0.7995 | 0.5212 |
| **Model Composition** | | | |
| Non-hydrogen atoms | 26373 | 28915 | 24854 |
| Protein residues | 3328 | 3647 | 3146 |
| ADP residues | 5 | 6 | 6 |
| **Model Refinement** | | | |
| Resolution limit (Å) | 2.84 | 2.66 | 3.85 |
| Number of chains | 11 | 11 | 12 |

| | | | |
|---|---|---|---|
| Number of residues | 3328 | 3647 | 3146 |
| **B factors (Å²)** | | | |
| Protein | 90.18 | 160.67 | 153.98 |
| Ligand (ADP) | 72.04 | 121.02 | 155.72 |
| **RMS deviations** | | | |
| Bond lengths (Å) | 0.003 | 0.002 | 0.003 |
| Bond angles (°) | 0.614 | 0.556 | 0.651 |
| **Validation** | | | |
| MolProbity score | 1.74 | 1.76 | 1.77 |
| Clash score | 4.79 | 5.25 | 7.95 |
| Rotamer outliers (%) | 2.03 | 2.11 | 1.69 |
| **Ramachandran plot (%)** | | | |
| Favored (%) | 96.21 | 96.46 | 97.04 |
| Allowed (%) | 3.64 | 3.32 | 2.73 |
| Outliers (%) | 0.15 | 0.22 | 0.22 |
